# P-element invasions in *Drosophila erecta* shed light on the establishment of host control over a transposable element

**DOI:** 10.1101/2022.12.22.521571

**Authors:** Divya Selvaraju, Filip Wierzbicki, Robert Kofler

## Abstract

To prevent the spread of transposable elements (TEs) hosts have developed sophisticated defence mechanisms. In mammals and invertebrates this defence mechanism operates through piRNAs. It is unclear how piRNA-based defences are established against invading TEs. According to the trap model, a TE insertion into a piRNA cluster, i.e. a distinct genomic locus, activates the host defence. Alternatively, siRNAs, generated by cleavage of dsRNA, may be the trigger for host control. To investigate this we introduced the P-element, one of the most widely studied eukaryotic transposons, into naïve lines of *Drosophila erecta*. We monitored the invasion in 3 replicates for more than 50 generations by sequencing the genomic DNA (using short and long-reads), the small RNAs and the transcriptome at regular intervals. A piRNA based host defence was rapidly established in 2 replicates but not in the third, where P-element copy numbers kept increasing for over 50 generations. We found that siRNAs emerged prior to piRNAs, supporting the view that siRNAs initiate host defence. However, neither insertions in piRNA clusters nor the formation of siRNAs were sufficient to stop the P-element. Instead the activation of the ping-pong cycle was shown to be crucial. We introduce a novel model, the crank-up model, which emphasizes activation of the ping-pong cycle as a critical event in establishing host control over a TE.

## Introduction

Transposable elements (TE) are sequences of DNA which selfishly spread in genomes. As this selfish activity enhances the transmission rate, TEs may spread in genomes even if this activity reduces host fitness [32, 72, 20]. TEs have been highly successful as they have invaded the genomes of virtually all eukaryotic species investigated so far [93]. TE content varies substantially among species, ranging from 3% in yeast to 77% in certain frogs [6]. Although some TE insertions could beneficially impact the host [13, 26], it is assumed that most TE insertions are neutral or deleterious [1, 68]. Theoretical work suggests that the selfish spread of TEs can reduce the fitness of host populations to such an extent that the survival of populations or even species is threatened [43, 44]. In agreement with this experimental populations invaded by a highly active TE went extinct after few generations [91]. Due to these deleterious effects host organisms have developed a broad range of sophisticated defence mechanisms, which frequently involve small RNAs [81]. In *Drosophila*, the host defence against TEs is based on piRNAs, small RNAs ranging in size of 23-29nt [28, 9]. These piRNAs bind to PIWI-clade proteins and mediate the repression of TEs at both the transcriptional and the post-transcriptional level [28, 9, 85, 51]. Most piRNAs are derived from discrete genomic source loci, termed piRNA clusters [9]. In *D. melanogaster* about 142 clusters were found, which account for around 3.5% of the genome [9, 97]. A central component of the piRNA pathway is the ping-pong cycle, where piRNAs, bound to the cytoplasmic proteins Aub and Ago3, direct the cleavage of TE transcripts [28, 9]. Cleavage by Ago3 yields novel piRNAs, which may in turn be loaded into Aub and vice versa. As a result, the ping-pong cycle amplifies the amount of piRNAs targeting a TE. Interestingly, cleavage by Ago3 (Aub) may also trigger a process termed ‘phasing’, where cleaved piRNA precursors are further processed into piRNAs by Zuc. These phased piRNAs are mostly bound by Piwi and mediate the transcriptional silencing of the TE in the nucleus. While the ping-pong cycle amplifies the amount of piRNAs, phasing is thought to increase the diversity of piRNAs targeting a TE [29, 65, 17]. piRNAs bound to PIWI-clade proteins have an important additional function: they are frequently maternally deposited into the egg and these deposited piRNAs are thought to define the position of piRNA clusters in the following generation [52, 31]. Furthermore, maternally deposited piRNAs likely initiate the ping-pong cycle in the next generation [52]. Therefore, once a host defence against a TE has been established, maternal piRNAs maintain the host defence against the TE in the next generation by defining the sites of piRNA producing loci and initiating the ping-pong cycle. However, in the case of a newly invading TE an important open question remains on how such a piRNA-based host defence gets established in the first place. There are two major hypothesis. The classical view, i.e. the trap model, holds that an insertion into a piRNA cluster triggers the production of piRNAs against an invading TE, which in turn direct the TE’s silencing [4, 60, 98, 27, 97, 73]. This view is supported by several lines of evidence. Artificial sequences inserted into piRNA clusters generate piRNAs against these sequences [67]. A single insertion in a piRNA cluster, such as X-TAS or 42AB, is sufficient to silence a reporter [35, 57]. However recently an alternative hypothesis was suggested: maternally inherited siRNAs, produced from Dicer-2 mediated cleavage of dsRNA, could trigger the conversion of a locus into a piRNA producing site [57]. Such dsRNA may be readily formed from sense and antisense transcripts of TEs [57]. Both models, the trap and the siRNA model, implicitly assume that the entire piRNA pathway, including the ping-pong cycle and generation of the downstream phased piRNAs, is activated once the piRNA based host defence has been triggered either due to a cluster insertion or the emergence of siRNAs. In agreement with this ping-pong signatures were observed for both, artificial sequences inserted into existing piRNA clusters and piRNA-producing loci initiated by siRNAs [67, 57]. To shed light on the establishment of host defences against an invading TE, we introduced the P-element, one of the most widely studied eukaryotic transposons, into naïve lines of *D. erecta*. The P-element was originally discovered as the cause of hybrid dysgenesis, where crosses of males having the P-element to females without led to various symptoms, such as atrophied ovaries. By contrast, the offspring of reciprocal crosses exhibits no such symptoms [40, 5]. The P-element is a 2907bp DNA transposon with 4 ORFs which encode a single protein, the transposase [5, 71]. Interestingly, the P-element is active solely in the germline but not in the soma. It is thought that this is a strategy by the TE to avoid deleterious fitness effects to hosts [11]. This tissue specific activity of the P-element is regulated by alternative splicing of it’s third intron (IVS3). Retention of IVS3 in the soma leads to a non-functional transposase [50]. Interestingly, it was recently proposed that the piRNA pathway regulates P-element activity by suppressing IVS3 splicing in the germline rather than by regulating the expression of the P-element [87]. However in contrast to this, reciprocal crosses among flies with and without the P-element suggest that P-element expression is markedly reduced in offspring with piRNA-based defence against the P-element [38, 66].

In addition to piRNAs, P-element activity may also be regulated by certain non-autonomous P-element insertions, containing internal deletions, such as the KP-element or D50 [7, 76]. Expression of these elements with internal deletions leads to non-functional transposases that may occupy the transposase binding sites and thus prevent functional transposases from mobilizing the P-element [53].

Here, we monitored the P-element invasion for over 50 generations in experimental *D. erecta* populations in three replicates. At regular intervals we sequenced genomic DNA (using long and short reads), the small RNAs and the transcriptome. The P-element spread rapidly in all three replicates. In two replicates copy numbers of the P-element stabilized around generation 20, due to activation of the piRNA pathway, including the ping-pong cycle. We observed siRNAs complementary to the P-element emerging prior to piRNAs, supporting the hypothesis that maternally deposited siRNAs initiate the host defence against an invading TE. The emergence of piRNAs led to down-regulation of IVS3 splicing but had little effect on P-element expression. In one replicate the host defence against the P-element was not established and P-element copy numbers kept increasing for over 50 generations. Despite the presence of siRNAs, piRNAs and piRNA cluster insertions, the ping-pong cycle targeting the P-element was not activated in this replicate until generation 70. Our data thus suggest that neither piRNA cluster insertions nor the presence of siRNAs against the P-element are sufficient to activate the whole host control. Therefore, we introduce a novel model, the crank-up model, which emphasizes the activation of the ping-pong cycle as the critical event which establishes host control over an invading TE.

## Results

### P-element invasion in *D. erecta*

To investigate the dynamics of TE invasions and the establishment of host control, we monitored P-element invasions in experimental populations of *D. erecta*, a species that does not have P-element insertions (i.e. naïve species) [10]. We utilised the *D. erecta* strain 01 as this strain is highly inbred and was used for generating the reference genome [21, 41]. We first confirmed the absence of the P-element in strain 01 using PCR and Illumina sequencing (supplementary figs S1, S2A). Next we introduced the P-element of *D. melanogaster* into the *D. erecta* strain 01 via micro-injection of a P-element carrying plasmid into embryos (ppi25.1; kindly provided by Dr. Erin Kelleher). The transformed flies were screened for presence of the P-element using PCR (supplementary fig. S2) and maintained in the lab for 3 generations before we mixed them with naïve *D. erecta* flies of strain 01. The experimental populations were maintained at a population size of *N* = 250 for 50 generations. We used non-overlapping generations, 3 replicates and a constant temperature of 25°C. As the populations are based on highly inbred lines with a low level of polymorphism, the influence of selection should be minimal (in contrast to previous works where the P-element invaded populations with high levels of standing genetic variation [47, 48]). We investigated the spread of the P-element by monitoring several key parameters: the abundance of P-element insertions, the extent of gonadal dysgenesis, the expression and splicing of the P-element and the amount of piRNAs against the P-element. For an overview of our experimental design see supplementary fig. S3.

To trace the spread of the P-element we sequenced the populations as pools (Pool-Seq [82]) at about each 10^*th*^ generation using Illumina paired-end sequencing (for an overview of all used Pool-Seq samples see supplementary table S1). We estimated the number of P-element insertions with DeviaTE [92], which normalizes the coverage of the P-element to the coverage of single copy genes. Initially, P-element copy numbers rapidly increased in all three replicates but the spread considerably slowed around generation 20 in replicates 1 and 4 (fig. 1A; supplementary figs S4-S6). By generation 48, each haploid genome carried about 27 and 37 P-element insertions in replicate 1 and 4 respectively (supplementary table S2). In contrast to this, P-element copy numbers continued to increase in replicate 2. By generation 48 each haploid genome accumulated 151 P-element copies in replicate 2 (i.e. a staggering 302 P-element insertions per fly; fig. 1A; supplementary table S2). At late generations (*≥* 34) P-element copy numbers are significantly higher in replicate 2 than in replicates 1 and 4 (Wilcoxon rank sum test *p* = 0.024; supplementary table S2). The effective transposition rate (*u*′) in replicate 2 is predominantly higher than in the other two replicates (supplementary table S2). Note that we are solely able to measure the effective transposition rate, i.e. the novel insertions gained through transpositions, minus the insertions lost via negative selection against the TEs (*u*′ = *u − x*). An analysis independent of DeviaTE, based exclusively on the fraction of raw reads mapping to the P-element, confirms that the P-element proliferates in replicate 2 while the invasion is largely controlled by generation 20 in replicates 1 and 4 (supplementary fig. S7; supplementary table S2).

**Figure 1:**
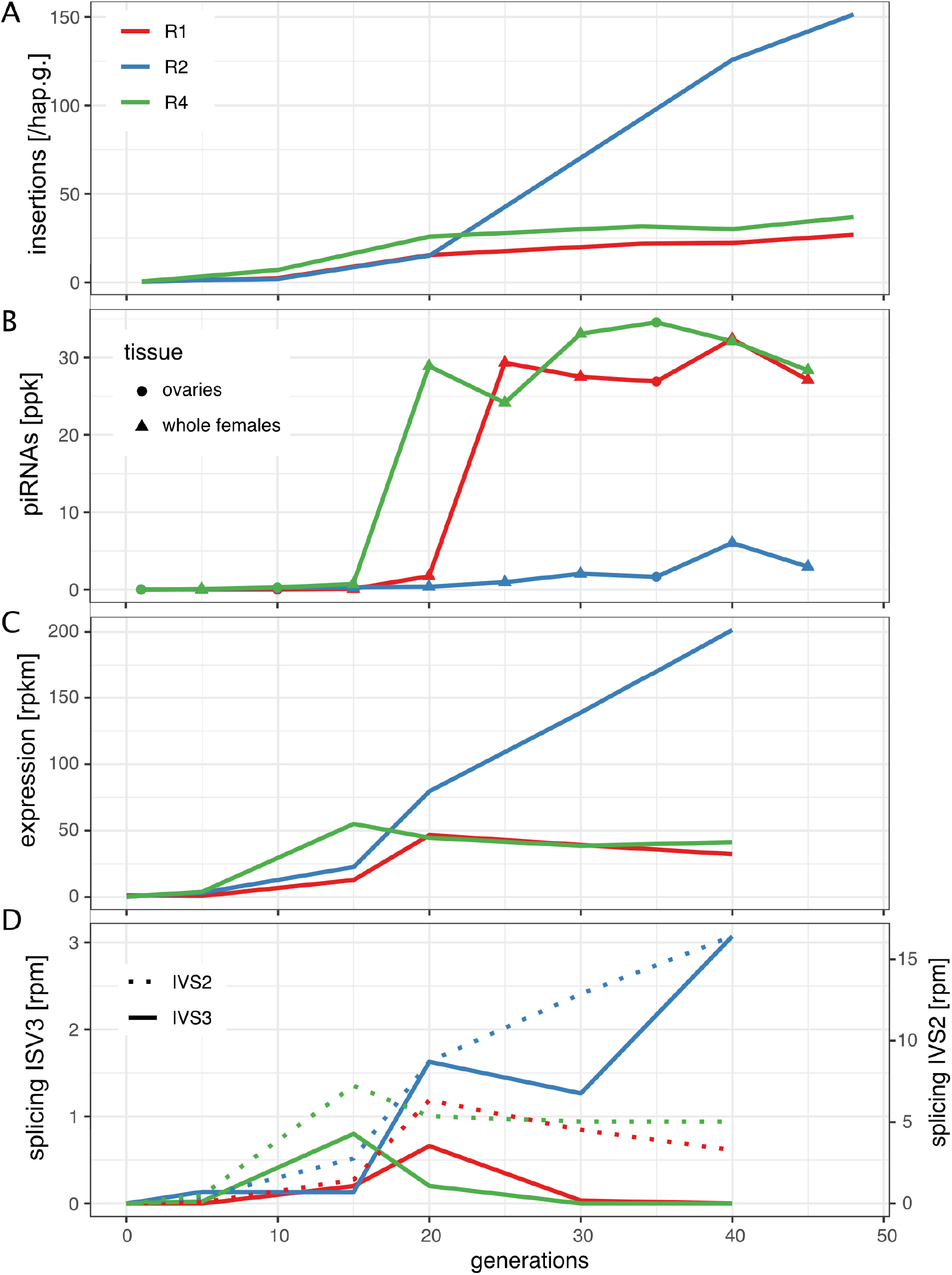
Invasion dynamics of the P-element in experimental *D. erecta* populations. Data are shown for 3 replicate populations (R1, R2, R4). A) Insertions per haploid genome (hap.g.) during the invasion. B) Abundance of piRNAs complementary to the P-element [ppk: P-element piRNAs per 1000 piRNAs]. C) Sense expression of the P-element in whole female flies. Naïve flies are shown at generation zero. D) Splicing of the second (IVS2) and the third (IVS3) intron of the P-element. Naïve flies are shown at generation zero [rpm spliced reads per million mapped reads].

In *Drosophila* TE activity is largely controlled by piRNAs, small RNAs ranging in size between 23 and 29nt [9, 28]. To test whether piRNAs against the P-element emerged in our experimental populations we sequenced small RNAs at every 5th generation during the experiment. Initially, we aimed to sequence ovarian RNA but due to the huge workload associated with repeated dissections of large numbers of ovaries, we sequenced whole bodies of female flies at later generations (for an overview of all sequenced small RNA libraries see supplementary table S3). Nevertheless, since piRNAs are usually solely found in ovaries of many *Drosophila* species, including *D. erecta* [54, 95], we reasoned that it should be feasible to compare the piRNA abundance between ovaries and whole females when reads are normalized to a million piRNAs. To investigate this, we sequenced small RNAs from ovaries and whole bodies of naïve *D. erecta* flies using three biological replicates. As miRNAs are abundant in the soma and the germline [61] normalization to a million miRNA yielded diverging estimates between ovaries and whole female flies (supplementary fig. S8). However, normalization to a million piRNAs yielded similar estimates of the piRNA abundance among the two tissues (supplementary fig. S8). For the remainder of this work we thus normalized the abundance of small RNAs to a million piRNAs.

Only very few piRNAs complementary to the P-element were found in the naïve *D. erecta* strain01 (one read in the ovaries, and three reads in the whole bodies; fig. 3). In the experimental populations, piRNA copy numbers rapidly increased in replicates 1 and 4 around generation 20 but remained at a significantly lower level in replicate 2 (fig. 1C; Wilcoxon rank rum test at generations *≥* 25, *p* = 0.00067). In replicates 1 and 4 most piRNAs have a length between 25 and 27nt and are antisense to the P-element, as expected for piRNAs (supplementary fig S9) [88, 9]. Furthermore, these piRNAs show a pronounced U-bias at the first nucleotide, as described for piRNAs bound to Aub or Piwi (supplementary fig. S10; [9, 79] The piRNAs are distributed all along the sequence of the P-element (supplementary fig. S11). Our data thus suggest that the P-element in replicates 1 and 4 around generation 20-25 is largely controlled by piRNAs, whereas the abundance of piRNAs may be insufficient to stop the P-element invasion in replicate 2 (fig. 1).

We next asked how piRNAs act to control the P-element invasion in replicates 1 and 4. Reciprocal crosses among flies with and without the P-element showed that P-element expression is markedly reduced (4 to 10 fold) in flies with maternally inherited P-element piRNAs as compared to flies lacking these piRNAs [38, 66]. This suggests that piRNAs regulate the expression of the P-element. On the other hand, a recent work proposed that piRNAs do not reduce the transcript levels of the P-element but rather act to regulate the splicing of the third intron (ISV3) [87].

To investigate how piRNAs control the P-element, we performed stranded RNA-Seq at about each 10^*th*^ generation (for an overview of the RNA-Seq data see supplementary table S4). While very few RNA-Seq reads map to the P-element in naïve flies (0-1 reads), large numbers of reads align to the P-element in our experimental populations (supplementary table S4). The position of the P-element introns is identical in *D. erecta* and *D. melanogaster* (supplementary fig.S12, S13). In replicates 1 and 4 the expression (sense) of the P-element in whole flies increases until generations 15-20 and remains at a high level thereafter (fig. 1C; supplementary fig. S12). In replicate 2 the expression of the P-element increases until generation 40. Since piRNAs are solely expressed in the germline of most *Drosophila* species, it may be argued that somatic P-element expression could mask a locally reduced P-element expression in ovaries when whole flies are sequenced. We therefore also sequenced P-element expression in ovaries at generations 10 and 35 (supplementary fig. S13). In ovaries, the P-element expression increased from generation 10 to 35 (supplementary fig. S13). At generation 35, P-element expression in ovaries was even higher than in whole flies (normalized to rpkm; supplementary figs. S12, S13). Our data thus suggest that the emergence of piRNAs does not cause a marked decrease in P-element expression. In comparison, the level of splicing of IVS3 showed a pronounced response to the emergence of piRNAs (fig. 1D). In replicates 1 and 4 the level of splicing of IVS3 increased until generations 15-20 but splicing of IVS3 stopped at later generations (fig. 1D; supplementary figs. S12, S13). Only in replicate 2, where few piRNAs against the P-element emerged, did splicing of IVS3 continue to increase. In the ovaries, we also found that splicing of IVS3 stopped by generation 35 in replicates 1 and 4 but not in replicate 2 (supplementary figs. S13). In contrast to IVS3, the number of spliced reads for the other two introns of the P-element (IVS1, IVS2) remained at a high level in all replicates (fig. 1D; supplementary fig. S14). A linear model suggests that piRNAs have a significant negative effect on the splicing level of IVS3 and IVS1 but no effect on the splicing level of IVS2 nor on the expression level of the P-element (supplementary table S5; *p <* 0.01). Overall, our data support the hypothesis of Teixeira et al. [87] that P-element piRNAs primarily act to repress the splicing of the P-element’s third intron, thus preventing the expression of functional P-element transcripts.

A typical phenotypic effect associated with P-element activity are atrophied ovaries (gonadal dysgenesis: GD) in the offspring of flies without an effective piRNA based defence against the P-element. Since atrophied ovaries are largely observed at high temperatures we performed GD assays at 29°C and not at the experimental conditions (25°C; for an overview of GD assays see supplementary fig. S15). We observed a substantial number of atrophied ovaries, well above background levels, in all three replicates in the early stages of the invasion (supplementary fig. S16, supplementary table S6). In replicates 1 and 2 at generation 20 we found 100% atrophied ovaries. We were unable to estimate GD beyond generation 34 in replicate 2 due to an insufficient number of eclosed female flies. Of note, replicate 2 could not be maintained beyond generation 62 since not enough female flies eclosed at the experimental conditions (25°C). We were able rescue the population by back-crossing females of replicate 2 to naïve males. It is likely that the accumulating load of deleterious P-element insertions was driving replicate 2 to extinction (see also Lama et al. [49]).

### Insertions in piRNA clusters

We next investigated the reasons as to why the P-element was silenced in replicates 1 and 4 but not in replicate 2. Prevailing opinion states that a TE invasion is silenced when a copy of the invading TE jumps into a piRNA cluster, which is thought to activate the piRNA pathway (the trap model). One important aspect that is frequently neglected in functional discussions of the trap model is that a TE insertion in a piRNA cluster will initially be solely present in a single individual of the population (i.e. 1 out of 250 flies in our experiment) and thus the TE is initially silenced only in a single individual. In order to inactivate the TE throughout the population, silencing needs to spread to all individuals of a population. For this, three scenarios are feasible.

First, a cluster insertion is positively selected and sweeps through the population until all individuals carry the same cluster insertion (sweep model) [8] Under this sweep model we expect to find at least one fixed cluster insertion in replicates 1 and 4 around generation 20 but not in replicate 2. Finding TE insertions in piRNA clusters is challenging, as piRNA clusters are highly repetitive and thus difficult to assemble. To facilitate the discovery of P-element insertions in clusters, we relied on a recent long-read assembly of *D. erecta*. The annotation of piRNA clusters was based on small RNA data from ovaries of naïve *D. erecta* flies and a previously described algorithm [47, 48]. To distinguish germline (dual-strand clusters) from somatic clusters (uni-strand clusters), we also sequenced small RNAs from embryos of naïve *D. erecta* flies. To identify P-element insertions in the repetitive piRNA clusters with high-confidence, we sequenced the experimental populations at multiple generations using ONT long-read sequencing (for an overview of the long-read data see supplementary table S7). We did not find a single fixed cluster insertion around generation 20 in any sample from all three replicates (supplementary table S8). A cluster insertion in contig 508 at generation 18 of replicate 2 had the highest population frequency (G18-L1: *f* = 0.14; fig. 2; supplementary table S8). In fact, an analysis of the population frequencies of P-element insertions based on Illumina short read data suggests that not a single P-element insertion is fixed in the experimental populations (supplementary figs S17, S18, S19). Our findings are thus not in agreement with the sweep model.

**Figure 2:**
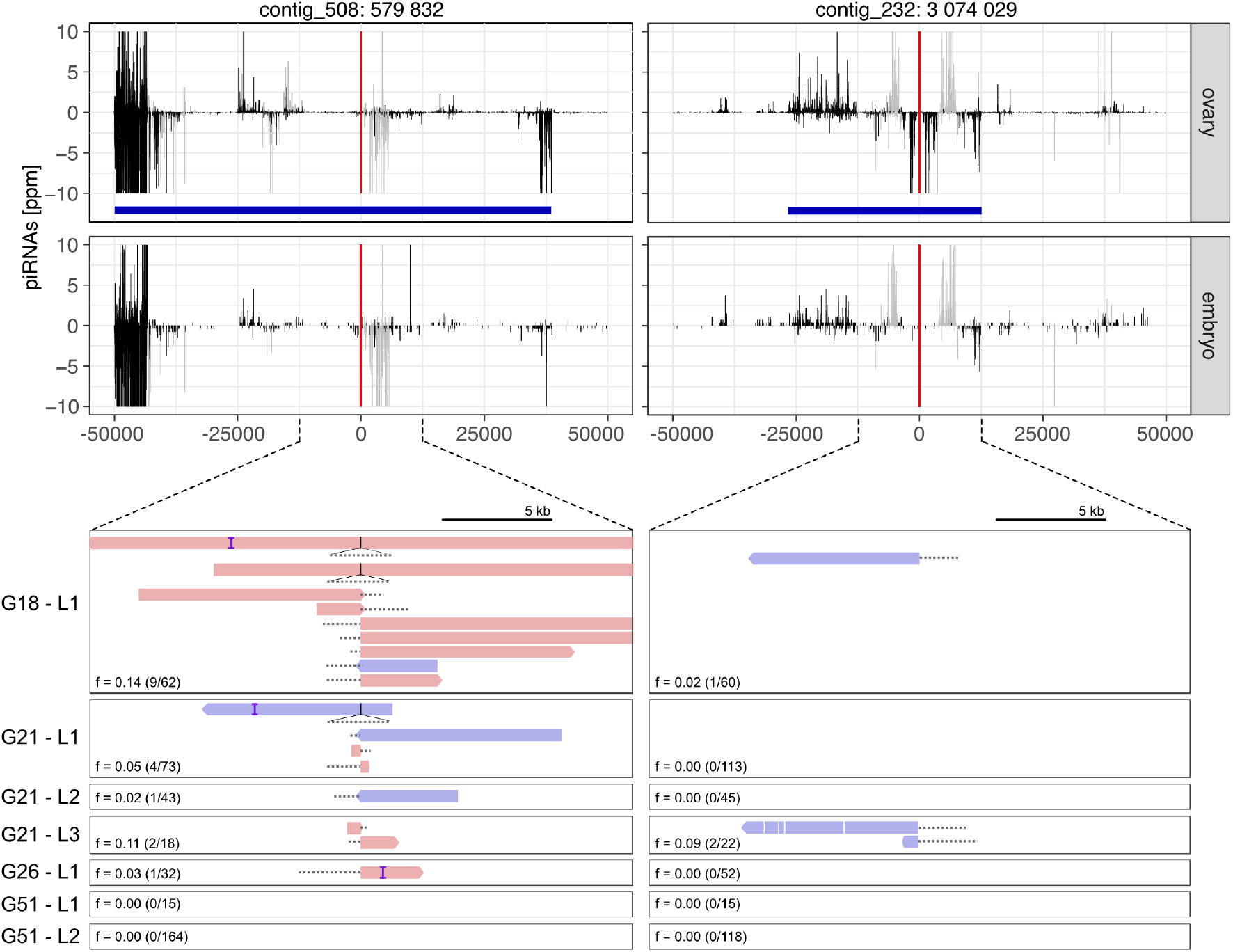
At least two P-element insertions in germline clusters are present in replicate 2 at early stages of the invasion (generations 18-26). Top panels show the abundance of piRNAs in a 100kb window around the insertion sites in both ovaries and embryos. Sense piRNAs are on the positive y-axis and antisense piRNAs on the negative y-axis. Bottom panels show the support of the P-element insertions by ONT long-reads at different generations (*Gxx*) and in different sequencing libraries (*Lx*). Sense reads are shown in red and antisense reads in blue. Regions of the reads aligning to the P-element are shown as dashed lines (true to scale). For each library we indicate the population frequency of the P-element insertion (*f*), the number of reads supporting the insertion and coverage at the insertion site (in brackets: reads/coverage). Note that both insertions are likely full-length P-elements (almost the complete P-element is covered by some reads) and that both insertions are likely lost (or at very low frequency) by generation 51.

Second, it was proposed that due to a high TE activity, many different cluster insertions emerge independently, such that a TE invasion will be stopped by a high number of segregating cluster insertions (shotgun model) [47, 43, 36]. Each individual in a population will thus carry a distinct sets of cluster insertions. Surprisingly, computer simulations under this scenario show that a TE invasion is stopped when each diploid individual carries, on average, about four cluster insertions, though it was assumed during the simulations that a single insertion per diploid is sufficient to silence the TE [43]. Recombination and random assortment among segregating cluster insertions will lead to a distribution of cluster insertions in populations, where some individuals will end up with several cluster insertions and others with just a few or even none. The TE will be active in the individuals without a cluster insertion. Solely when diploids carry an average of about four cluster insertions, the vast majority of the offspring will end up with at least one cluster insertion. Under this model, we expect to find around four cluster insertions at generation 20 in replicates 1 and 4 and fewer in replicate 2. Based on our ONT long-read data, we find that all replicates only carry between 3 to 29% of the required number of cluster insertions (*R*1 = 10%, *R*2 = 3 *−* 29%, *R*4 = 6%; supplementary table S9). Therefore, our data is not in agreement with the shotgun model. This is also consistent with our previous work, where we found an insufficient number of cluster insertions in experimental *D. simulans* populations being invaded by the P-element [47, 48].

Finally, it was noted that dispersed TE insertions may generate piRNAs [64, 84]. Since the deletion of large piRNA clusters did not lead to an activation of TEs, it was proposed that these dispersed TEs have an important role in the silencing of TEs [23]. The conversion of a regular TE insertion into a piRNA producing locus (the so called ‘paramutation’) may be triggered by maternally deposited piRNAs [19, 52, 31]. In the case of an invasion of a novel TE, this raises an important question on the origin of the very first piRNAs which could trigger such paramutations. One option is that an insertion into a piRNA cluster triggers the origin of the very first piRNAs against an invading TE. Once such initial piRNAs have emerged, increasing numbers of regular TE insertions, in different individuals, may be converted into piRNA producing loci as the invasion progresses (paramutation model). Under this model, we expect to find at least one cluster insertion, possibly at a low population frequency, in replicates 1 and 4 but not in replicate 2. While we found some cluster insertions at early generations (around generation 20) in replicates 1 and 4 (supplementary table S8) we also found two cluster insertions in replicate 2 (fig. 2). These cluster insertions in replicate 2 are likely reliable, as they are supported by several long-reads from different strands in multiple sequencing libraries of distinct generations (fig. 2). As piRNAs are found in both the ovaries and embryos of naive flies and piRNAs align to both strands, the two insertion sites are likely present in germline clusters (dual-strand clusters [60, 17]). An analysis of genomic short read data also supports the idea that cluster insertions are present in all replicates, including replicate 2, at early generations (albeit different ones; supplementary figs S18). Consequently, our data suggests that cluster insertions in replicate 2, which were present from generations 18 to 26, were not sufficient to establish the host defence mechanism against the P-element. Nevertheless, the cluster insertions could still be necessary precondition for activating the host defence.

In summary, we are able to rule out the idea that a fixed cluster insertion or several segregating cluster insertions are responsible for establishing host control over the P-element in our experimental populations. We cannot rule out that cluster insertions are necessary to trigger the generation of the first piRNAs against an invading TE, which leads to paramutations that convert increasing numbers of TE insertions into piRNA producing loci as the invasion progresses. However, our data suggest that cluster insertions and a few first piRNAs are insufficient to establish host control over the P-element.

### 0.1 siRNAs emerge before piRNAs

In addition to piRNAs, siRNAs (20-22nt) may also contribute to silencing of TEs [14, 16, 2]. In contrast to piRNAs, which are solely found in the germline of most *Drosophila* species, siRNAs are found both in the germline and the somatic cells [54]. Generation of siRNAs occurs by cleavage of dsRNA which could, for example, be formed by sense and antisense transcripts of a locus [57]. The siRNA pathway is distinct from the piRNA pathway, relying on entirely different sets of enzymes (e.g. Dicer2, Ago2) [88, 16]. However, recent work revealed an unexpected link between the piRNA and siRNA pathway [57]. It was suggested that siRNAs could initiate the conversion of TE insertions into piRNA producing loci [57]. siRNAs are an attractive mechanism for establishing host control over an invading TE, as they provide a simple means to distinguish self (genes) from non-self [57]: whereas dispersed TE insertions could readily generate sense and antisense transcripts, that may form dsRNA (and thus siRNAs), genes are rarely transcribed from both strands. In agreement with this, we found that in our RNA-Seq libraries about 7.41% of all P-element transcripts are antisense. As a control, solely 0.25% of the reads aligning to all *D. erecta* transcripts are antisense, which is significantly lower than for the P-element (Wilcoxon rank sum test *W* = 0, *p* = 7.4*e −* 07). Importantly, antisense transcripts of the P-element were found in all replicates already at generation 5 (supplementary fig. S20). Furthermore, antisense transcripts were found in RNA extracted from ovaries and whole flies (supplementary fig. S20). Therefore, the substrate necessary for generating siRNAs is already present at early generations of our experimental populations.

If siRNAs activate piRNA producing loci against the P-element [57], we expect siRNAs to emerge before piRNAs in the experimental populations. In contrast, if insertions into piRNA clusters trigger the host defence, we expect piRNAs to emerge independently of siRNAs. To investigate this, we monitored the length distribution of small RNAs during the invasions at every 5th generatio. We assumed that small RNAs with a length between 20-22nt and 23-29nt correspond to siRNAs and piRNAs, respectively. In naïve flies and at early generations of the experimental populations, few small RNAs aligning to the P-element were found and the length distribution of these RNAs was non-specific (fig. 3). However, at around generation 10-15 siRNAs peaks emerged in all replicates. Apart from the distinct length (about 21nt) these small RNAs have additional features typical for siRNAs, such as the balanced strand bias (fig. 3) and a less pronounced 5’-U bias than the piRNAs (supplementary figs. S21, S10, [16, 37, 25]). These siRNAs were distributed over the entire sequence of the P-element (supplementary fig. S22). The piRNA peaks, with lengths between 23-29nt, emerged only at later generations in replicates 1 and 4 but not in replicate 2 (fig. 3). In replicate 2 the abundance of siRNAs reached similar levels as the abundance of piRNAs in replicate 1 and 4 (supplementary fig. S23). However, in replicate 2 the abundance of siRNAs was much lower in the ovarian sample than in whole flies (generation 35; supplementary fig. S23), suggesting that many siRNAs are of somatic origin. Nevertheless, abundant siRNAs were also present in the ovarian samples of all replicates (generation 35; fig. 3).

**Figure 3:**
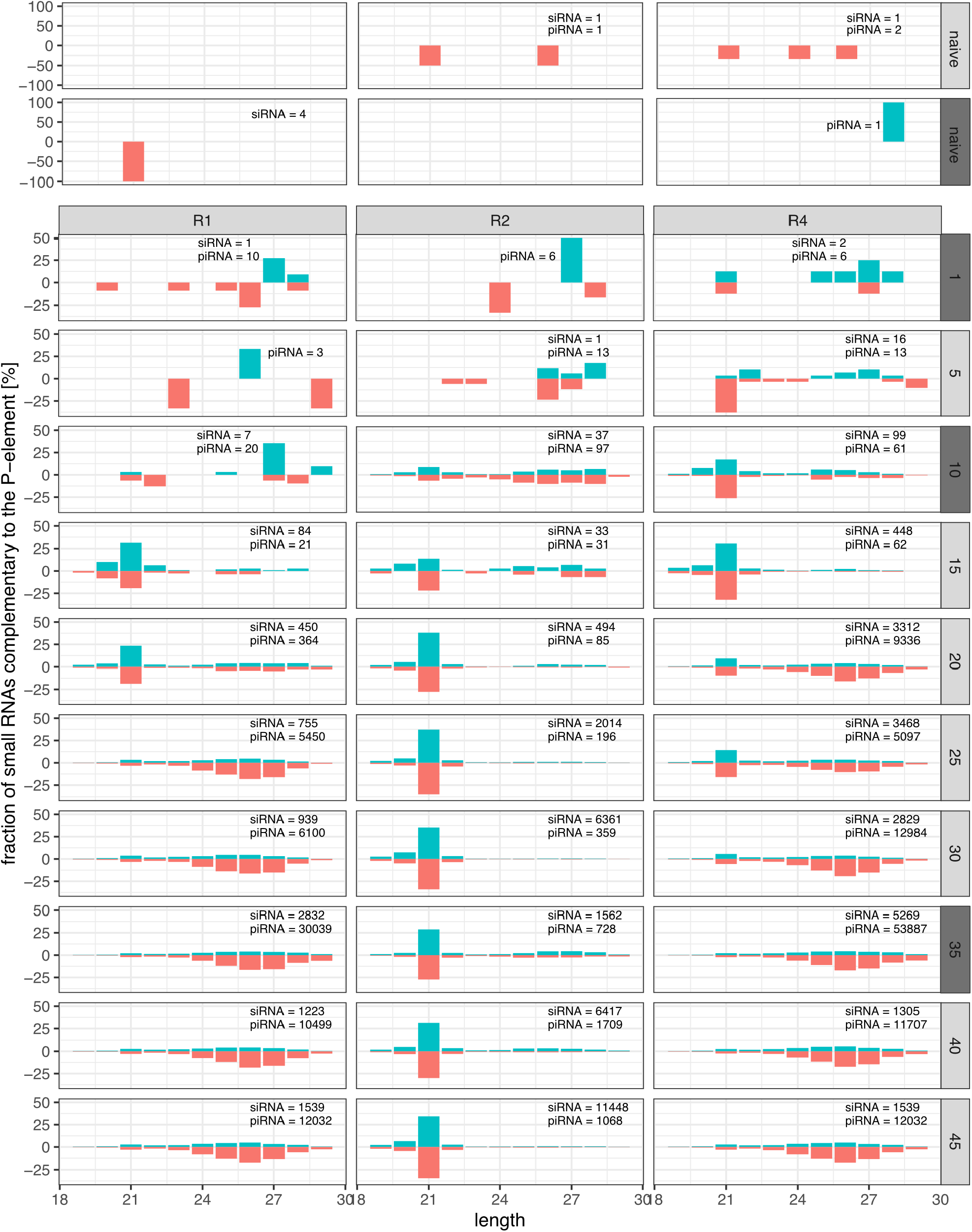
Length distribution of small RNAs mapping to the P-element during the invasions. Data are shown for naïve flies and the three replicates (top panel) at all five generations (right panel). We extracted small RNAs, either from whole bodies of female flies (light-grey panels) or from ovaries (dark-grey panels). For each sample we show the percentage of the small RNAs mapping to a given size category. In each sample the total number of reads accordingly adds to 100%. Sense RNAs are on the positive y-axis and antisense RNAs on the negative y-axis. The total number of siRNAs (20-22nt) and piRNAs (23-29nt) mapping to the P-element are shown.

In summary, we found that i) sense and antisense transcripts of the P-element are present at early generations of the invasion, ii) siRNAs are present in ovaries (where the host defence is typically acting) and iii) siRNAs emerged before piRNAs. Our data are thus consistent with the hypothesis of Luo et al. [57] that siRNAs may trigger the silencing of a TE. However, our data also suggest that siRNAs are insufficient to trigger the host defence alone, as abundant siRNAs against the P-element were also found in the replicate 2.

### Ping-pong is not active in replicate 2

The ping-pong cycle is an important mechanism which amplifies the abundance of piRNAs, slices TE transcripts and initiates transcriptional silencing of TEs in the germline. [9, 28, 83, 65]. We were interested in whether the ping-pong cycle is active against the P-element in our experimental populations, especially in replicate 2. The ping-pong cycle acts via the two PIWI clade proteins Aub and Ago3 [9, 28]. piRNAs bound to Aub usually mediate the slicing of sense transcripts of TEs, while piRNAs bound to Ago3 act on antisense transcripts. The cleavage products of Aub are loaded onto Ago3 and vice versa. Since the cleavage site of Aub is shifted by 10nt from the cleavage site of Ago3, a characteristic peak is observed (at position 10) when plotting the distance between the 5’ ends of sense and antisense piRNAs (i.e. the ping-pong signature [9, 28]). A pronounced ping-pong signature is thus characteristic for an active ping-pong cycle. In ovaries sampled at generation 35 we observed a clear ping-pong signature in replicates 1 and 4 but not in replicate 2, which suggests that the ping-pong cycle is inactive in replicate 2 (fig. 4). This absence of the peak at position 10 in replicate 2 was observed at all generations with sufficient amounts of piRNAs, irrespective of whether the small RNAs were extracted from whole flies or ovaries (fig. 4B; supplementary fig. S24). For most TE families, antisense piRNAs are largely bound to Piwi and Aub, whereas sense piRNAs are frequently associated with Ago3 [9, 83]. Piwi and Aub bound piRNAs frequently show a strong U-bias at position 1, whereas Ago3 bound piRNAs exhibit an A-bias at position 10 [9, 28, 15]. We find the pronounced U-bias at position 1 for antisense piRNAs of the P-element in all three replicates, in agreement with expectations for piRNAs (fig. 4; supplementary fig. S26). We also find the A-bias at position 10 for sense piRNAs in replicates 1 and 4 but not in replicate 2, where we find a U-bias at position 1 instead (fig. 4; supplementary fig. S25). This suggests that the ping-pong cycle is inactive in replicate 2, because Ago3 loaded piRNAs are absent.

**Figure 4:**
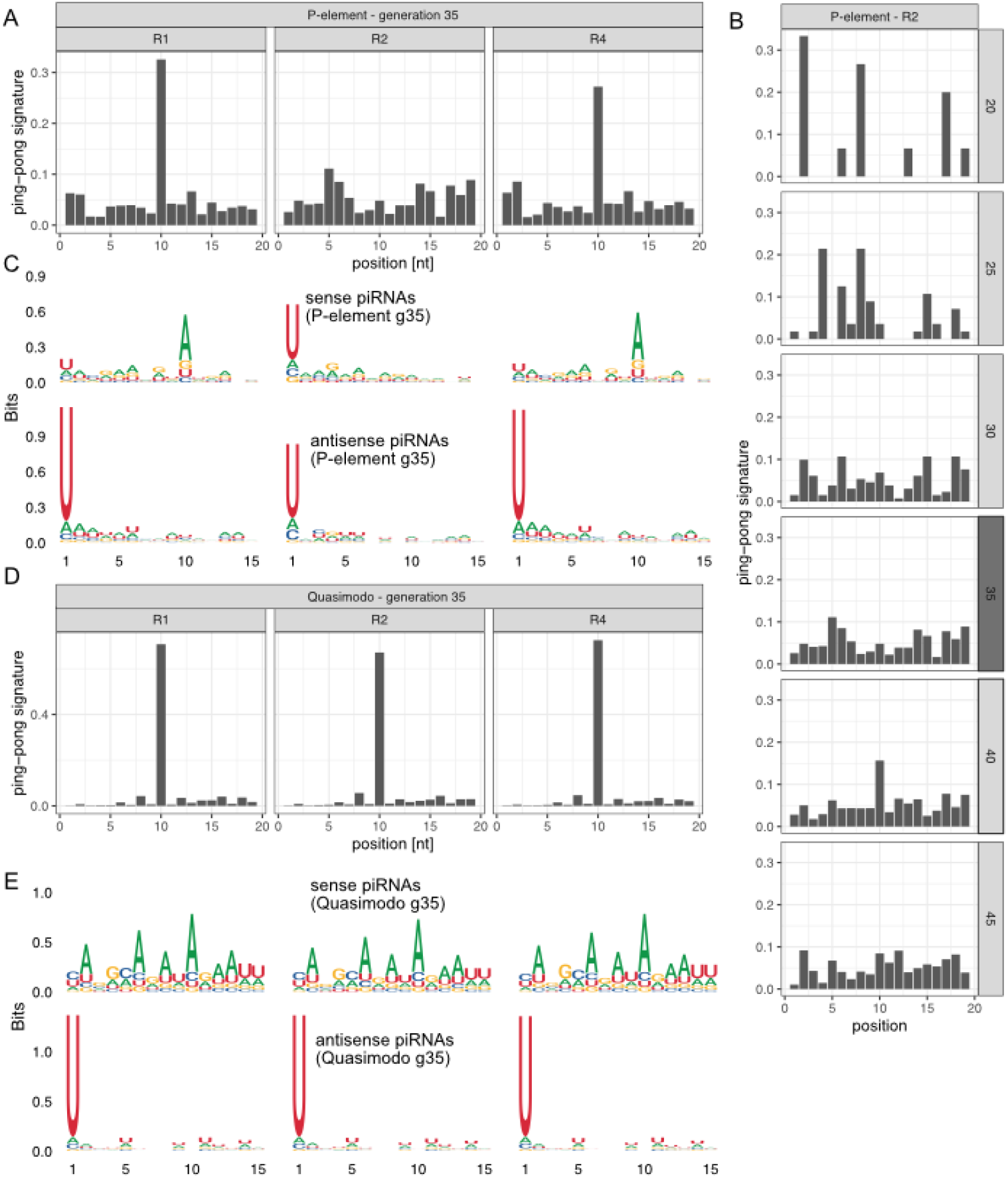
The P-element ping-pong cycle is inactive in replicate 2. A) The ping-pong signature (peak at position 10) of the P-element in ovaries at generation 35. Note that the ping-pong signature is missing in replicate 2. B) The ping-pong signature of the P-element in replicate 2. Data are shown for all generations with sufficient piRNAs. Small RNA extracted from ovaries and whole flies are shown in dark and light grey, respectively. C) Motifs of sense and antisense piRNAs (23-29nt) complementary to the P-element in ovaries at generation 35. Sense piRNAs in replicate 2 do not show an A-bias at position 10. D) Quasimodo displays a pronounced ping-pong signature in all replicates (ovaries, generation 35). E) Motifs of sense and antisense piRNAs of Quasimodo (ovaries, generation 35).

Our findings raise the question of whether the ping-pong cycle is defective in replicate 2. To answer this question, we investigated the ping-pong signatures and the small RNA motifs of other TEs in *D. erecta*. One major difficulty is that a reliable TE annotation is not available in *D. erecta*. By aligning *D. erecta* genomic reads to *D. melanogaster* TEs, we noticed that Quasimodo (LTR) and BS (non-LTR) show a high and continuous coverage (albeit with many divergent SNPs). This suggests that full length copies of these two TEs are present in *D. erecta*. BS and Quasimodo show clear ping-pong signatures in all three replicates (fig. 4D; supplementary fig. S27). Furthermore, the U-bias at position 1 of antisense piRNAs and the A-bias at position 10 of sense piRNAs is also present in all three replicates (fig. 4E; supplementary fig. S27). As follows, our data suggests that the ping-pong cycle is fully functional in all three replicates, including replicate 2.

In summary, we show that the ping-pong cycle against the P-element is active in replicates 1 and 4 but not replicate 2, although the ping-pong cycle is fully functional in all replicates.

### Differences among replicates

Our data raises the important question as to why the ping-pong cycle was inactive against the P-element in replicate 2. Sense and antisense transcripts of TEs are the substrate of the ping-pong cycle. These transcripts are sliced by Aub and Ago3, thereby generating novel piRNAs [9, 17]. However, we showed that all replicates have a similar abundance of sense and antisense transcripts of the P-element (supplementary fig. S20). Furthermore, given that replicate 2 has the highest expression level of the P-element (fig. 1C), a lack of substrate cannot explain the absence of ping-pong in replicate 2.

It is feasible that there is a trade-off between the defence against TEs and viruses [78]. A virus infection may, for example, preoccupy important enzymes from the siRNA or piRNA pathway, such that they are no longer available for establishing a defence against an invading TE. We thus investigated the amount of small RNAs mapping to different *Drosophila* viruses but found similarly low numbers of reads aligning to viruses in all replicates (supplementary table S10).

In a recent landmark study Moon et al. [66] identified the gene Chk2 as a crucial factor for triggering the ping-pong cycle against the P-element. We thus speculated that expression of this gene may be compromised in replicate 2. However, FBtr0141271, the *D. erecta* ortholog of Chk2, has a very similar expression level among the three replicates at all time points (supplementary fig. S28).

It is, however, possible that we have not yet identified all genes required for triggering the ping-pong cycle. Thus we asked whether any of the *D. erecta* transcripts are differentially expressed among the replicates. Orthology to *D. melanogaster* genes was established with BLAST. Between the base population and the evolved lines (at generations 30 and 40) we mostly found that the P-element and some genes involved with circadian rhythm (tim, Pdp1) were differentially expressed (fig. 5A). The differential expression of circadian rhythm genes likely reflects slight differences in sampling times between evolved and naïve populations. Thus, we asked if any genes are differentially expressed between replicates with (1 and 4) and without (2) active P-element ping-pong. Except for the P-element, which is most highly expressed in replicate 2, we did not detect any significant differences among these replicates (fig 1C; fig. 5B).

**Figure 5:**
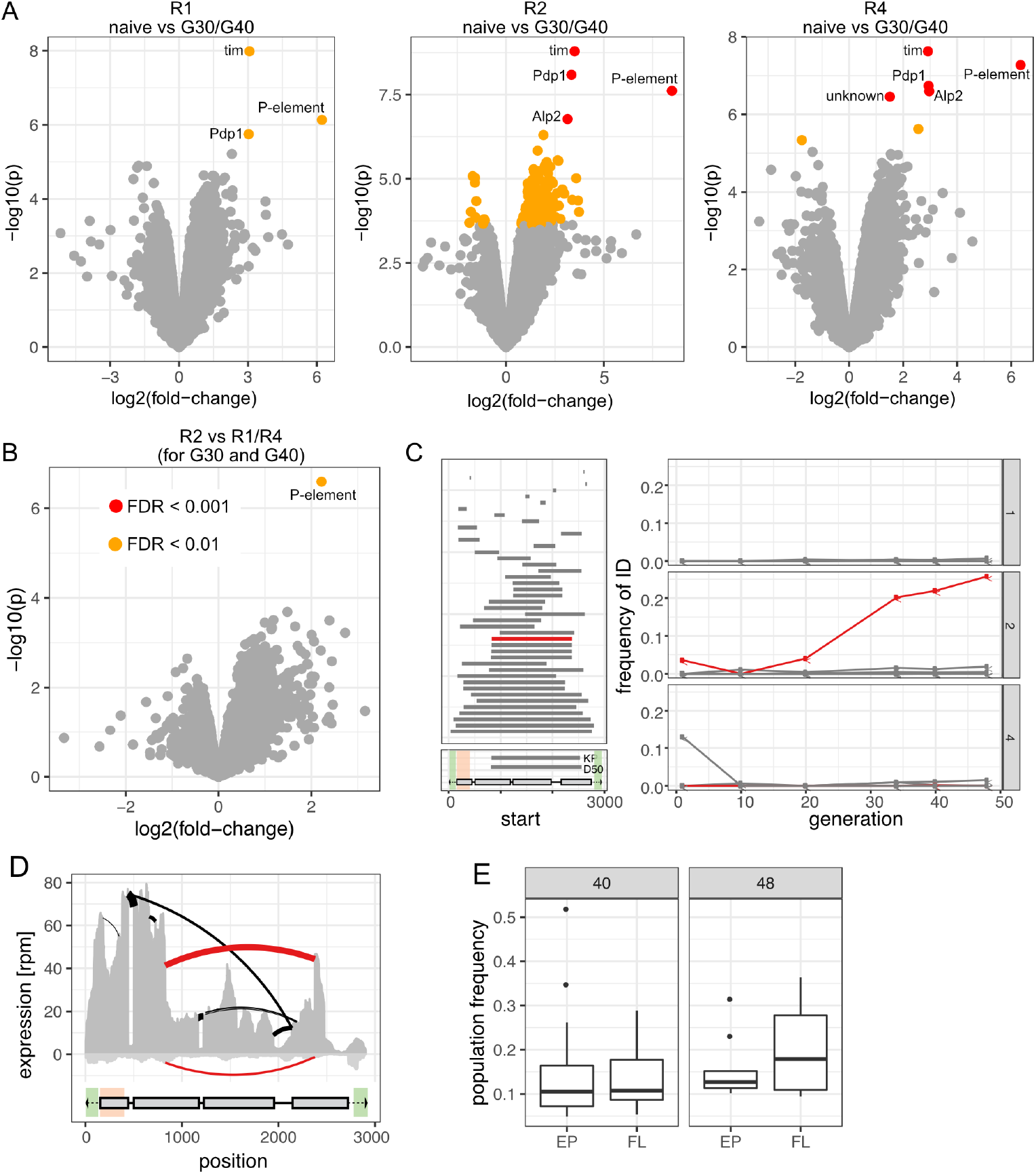
Differences among replicates with (R1,R4) and without (R2) P-element ping-pong. A) Volcano plots highlighting expression differences for TEs and *D. erecta* transcripts between naïve flies and invaded flies (generations 30 and 40). Data are shown for all three replicates (R1, R2, R4). B) Volcano plot highlighting expression differences between replicates with and without P-element ping-pong at generation 30 and 40. C) Overview of P-element insertions with internal deletions in the experimental populations (left panel). The lower left panel shows the composition of the P-element and the deleted regions of D50 and the KP-element. The DNA-binding domain (orange) and regions necessary for mobilizing the P-element (green) are indicated [58]. The right panel shows the frequency of the IDs (relative to all P-element insertions) in the three replicates. Note that the EP-element (red), i.e. a P-element variant with deletion of a similar region than for the KP-element, is increasing in frequency in replicate 2 but not in the other replicates. D) Sashimi plot showing that the EP-element (red arcs) is transcribed in replicate 2 at generation 35 in the sense (positive y-axis) and the antisense orientation (negative y-axis). Note that gaps in the mRNA (i.e. arcs) may result from splicing of mRNA-precursors as well as expression of P-elements with an internal deletions. E) Population frequency of the EP-element and the full-length P-element (FL) in replicate 2 at generations 40 and 48.

We next asked if differences in the sequence of the P-element might be responsible for the observed differences among the replicates. In terms of base substitutions, the sequence of the P-element is highly similar among all replicates. Only a few rare SNPs were found in all replicates (supplementary table S11). We were particularly interested in whether the abundance of internal deletions (IDs) of the P-element varies among replicates, as some IDs may repress P-element activity [7, 76]. IDs arise frequently during propagation of the P-element. The P-element propagates via a “cut-and-paste mechanism”, where gaps resulting from excision of a P-element are repaired using the sister chromatid as template [22]. Interruption of this gap-repair is thought to generate P-element insertions with IDs [22]. We used our previously published tool DeviaTE [92] to identify the location and abundance of IDs within the P-element. Based on split-reads (reads mapping to the breakpoints of an ID), DeviaTE quantifies the abundance of IDs relative to the total abundance of P-element insertions, but it is not possible to identify the genomic location of the IDs. We found 43 P-element IDs in the experimental populations (fig. 5C). The vast majority of the IDs (42/43) occurred in a single replicate, confirming our previous finding that IDs of the P-element are usually replicate specific [92]. Most of the IDs remained at a low frequency. However, one ID, where nucleotides 827-2375 are deleted (henceforth “EP-element”; for erecta P-element) rose to a frequency of 25.8% in replicate 2 (fig. 5C; supplementary figs S4-S6).

In *D. melanogaster* some insertions with IDs, like D50 or the KP-element, repress P-element activity [7, 76]. We were interested whether the EP-element might be one such repressor of P-element activity. The proteins encoded by the EP- and the KP-element are quite similar. Both deletions lead to a premature stop codon, where the resulting protein has a length of 207aa and 208aa for the KP- and the EP-element, respectively. Furthermore, for the EP-element, the first 206 codons are identical to the full-length P-element (which has 751 codons; [24]), while for the KP-element the first 199 codons are identical to the full-length element. The EP-element retains the DNA-binding domain (the first 88 codons [53]) but probably does not produce a functional transposase (the vast majority of the codons are missing). Similarly to the KP-element, the EP-element is therefore likely a repressor of P-element activity.

This raises the question on why the EP-element was rising to a high frequency in replicate 2. In principle two hypothesis are viable. The EP-element may be positively selected (as it reduces deleterious P-element activity) or it could be preferentially mobilized. These two hypothesis can be distinguished by investigating the population frequency of the different P-element insertions. Positive selection increases the population frequency of beneficial insertions. If EP-elements are positively selected their population frequency should be higher than the frequencies of FL insertions. In contrast, preferential mobilization leads to many novel insertions, and novel insertions initially have a low population frequency (1*/*2*N*). If EP-elements are preferential mobilized their population frequency should therefore, on average, be lower than the frequency of FL insertions. To address this question, we linked the information obtained from short and long-read sequencing. Long-read sequencing provides the identity of P-element insertions (e.g. distinguish between EP-elements and full-length insertions), while short-read sequencing provides estimates of the population frequency of P-element insertions. Based on the long-read data from generation 51 and the short-read data from generations 48 and 40, we found that EP-elements have a lower (albeit not significantly) population frequency than FL insertions (fig. 5E; Wilcoxon rank sum test *p*_48_ = 0.31, *p*_40_ = 0.64). The high abundance of the EP-element in replicate 2 can thus most likely be explained by preferential mobilization of the EP-element. This is also in agreement with previous works suggesting that P-elements with internal deletions may be more readily mobilized than FL insertions [34, 47, 86].

In summary, out of the investigated factors (viral load, fraction of antisense transcripts, Chk2 expression, expression pattern of known genes, SNPs and internal deletions of the P-element), solely the abundance of an internal deletion, the EP-element, was substantially higher in replicate 2 than in the other replicates.

### Influence of the genome vs. maternally deposited piRNAs

The presence of the EP-element in replicate 2 is the only notable difference among replicates with (replicates 1, 4) and without (replicate 2) P-element ping-pong. We were thus wondering if the EP-element could somehow interfere with the ping-pong cycle in replicate 2. To test this hypothesis, we performed reciprocal crosses among flies sampled from the replicates. Reciprocal crosses among replicates enable us to distinguish between the influence of the genome (which is identical among offspring of the reciprocal crosses) and the influence of the maternally transmitted factors such as piRNAs (which differs among the offspring of reciprocal crosses; fig. 6A). Flies for the crosses were sampled from the experimental populations between generations 67 and 70. Note that we had to back-cross females of replicate 2 with naïve males at generation 62 since the fecundity in this replicate was rapidly deteriorating (see above). For each cross we performed 3 subreplicates. We sequenced the small RNAs of the female parents and of the female F1 offspring (fig. 6A). Our small RNA data show that even by generation 67 all six samples of replicate 2 do not exhibit a ping-pong signature (fig. 6B), although the ping-pong cycle is fully functional in all subreplicates (supplementary fig. S29). Given that we found 74 P-element insertions in piRNA clusters in replicate 2 by generation 51 (based on ONT long-read sequencing) these data confirm that cluster insertions may not be sufficient to trigger the host response against an invading TE. However, the offspring of crosses among replicates only shows ping-pong signatures if the female is sampled from replicates 1 or 4 but not if females were sampled from replicate 2. As the genome is largely identical among offspring of the reciprocal crosses (e.g. R1xR2 vs R2xR1) these results suggest that maternally transmitted factors are responsible for the absence of ping-pong in replicate 2. Although, any maternally transmitted component, such as imprinting or the abundance of some protein, could be responsible, we suspect that maternally transmitted piRNAs are responsible, as we did not find expression differences for any transcript among the replicates and it has been shown that maternally deposited piRNAs are important for initiating the ping-pong cycle in the next generation [52]. Maternally transmitted piRNAs in replicate 2 thus seem to have a composition that is incompatible with the establishment of ping-pong. Our results also suggest that the EP-element is not responsible for the absence of ping-pong in replicate 2.

**Figure 6:**
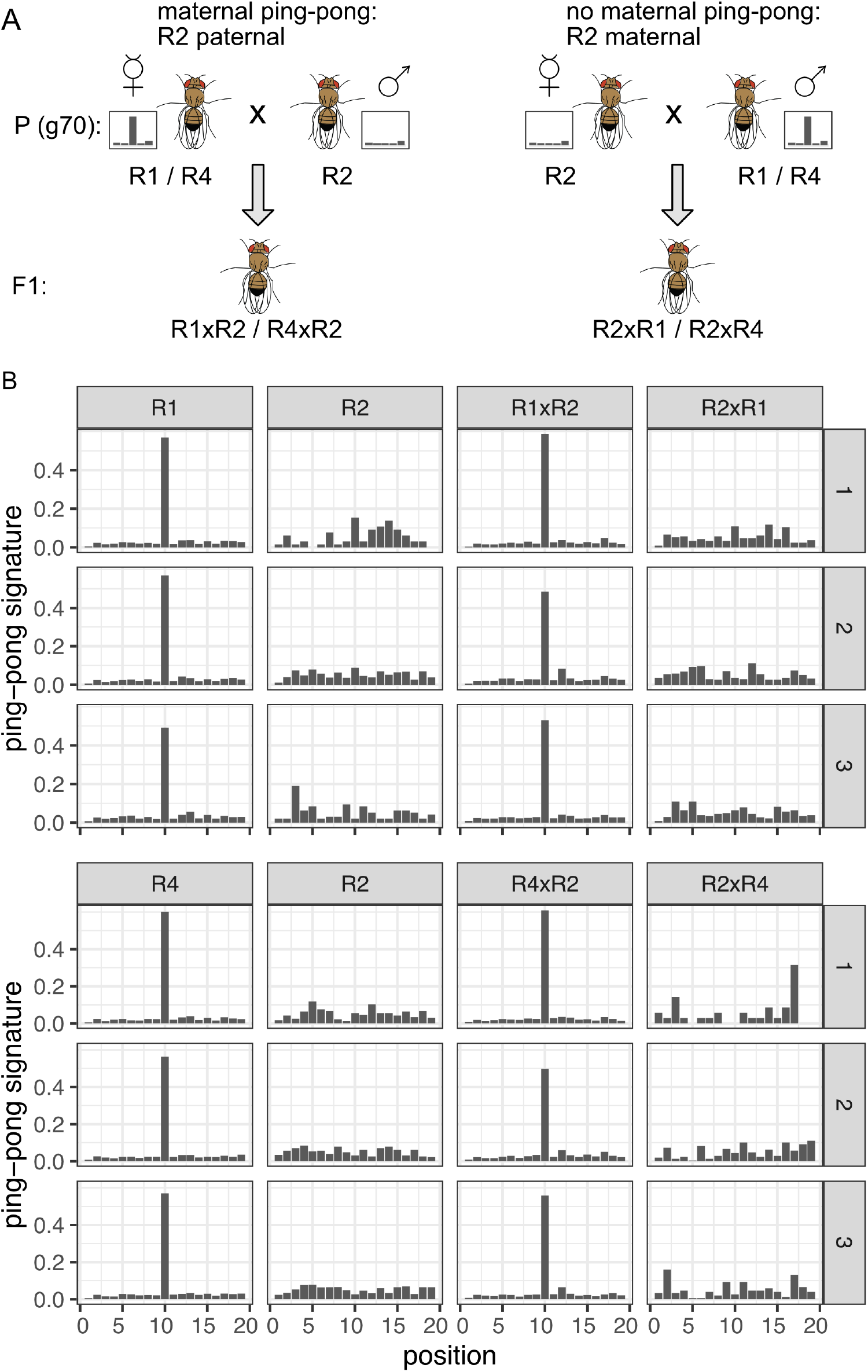
Maternally deposited piRNAs, not the genomic composition, are responsible for the absence of P-element ping-pong in replicate 2. A) Crossing scheme for testing the influence of maternal piRNAs. We performed reciprocal crosses among replicates with (R1,R4) and without (R2) ping-pong signature against the P-element. Note that in the F1 offspring of the reciprocal crosses (eg.: R1xR2 vs R2xR1), the genomic background is largely identical, while the composition of the maternally deposited piRNAs differs. The flies for this experiment were sampled around generation1770 from the experimental populations B) Ping-pong signatures for the P-element in all three replicates (R1, R2, R4) and in the F1 offspring of the reciprocal crosses among the replicates (e.g. R1xR2: R1-female x R2-male). Small RNA was extracted from whole female flies and three sub-replicates (right panel) were used for each sample.

Since the genome, and thus also the P-element insertions, is identical among reciprocal crosses, these results confirm our conclusion that differences in the abundance and quality of piRNA producing loci among replicates can not be responsible for the absence of the ping-pong cycle in replicate 2.

## Discussion

### Invasion dynamics in *D. erecta*

Here we introduced the P-element into a naïve *D. erecta* strain and monitored the ensuing invasions in several replicates using pooled genomic sequencing, ONT long-read sequencing, RNA-Seq, small RNA-Seq and gonadal dysgensis assays. Making this current study the most comprehensive investigation of a TE invasion to date.

It is perhaps not surprising that the P-element is able to multiply in *D. erecta*. The P-element is found in many species of the *willistoni* and *saltans* group [18]. In the last century it first invaded natural populations of *D. melanogaster* and then later populations of *D. simulans* [39, 45], indicating that the P-element is able to multiply in a broad range of species. Mobility assays further show that the P-element can be mobilized in all tested Drosophilids but not in Tephritidae [70]).

Overall, we found that the dynamics of the P-element invasions in *D. erecta* are very similar to other species. First, the P-element has a similar, albeit slightly higher, effective transposition rate in *D. erecta* (*u*′ = 0.217; average of first 20 generations; supplementary table S2;) than in *D. simulans* (*u*′ = 0.15; hot temperature; [47]). Second, the positions of the introns of the P-element are conserved between *D. melanogaster* and *D. erecta* (supplementary figs. S12, S13). Third, similar to other species, internal deletions of the P-element arise rapidly in *D. erecta* (fig.5; [48, 47, 7]). Fourth, we showed that the P-element is inducing gonadal dysgenesis in *D. erecta* similarly as in *D. melanogaster* and *D. simulans* [40, 33]. Gonadal dysgenesis is likely the consequence of developmental arrest of germline stem cells in ovaries induced by excessive P-element activity [66]. The presence of gonadal dysgenesis therefore suggests that the P-element is active in the germline stem cells of *D. erecta* ovaries, akin to *D. melanogaster* and *D. simulans* [47, 48, 91]. Finally, we found that piRNAs are likely not regulating the expression level of the P-element but rather the splicing of its introns, especially of IVS3 (fig. 1). However, we can not fully rule out that piRNAs are just repressing P-element expression in the germline stem cells, i.e. the cells where the P-element is thought to be active [66]. Germline stem cells contribute little bulk RNA to RNA-samples extracted from ovaries or whole flies, as performed here. We thus lack the sensitivity to detect expression differences solely taking place in a few germline stem cells. However we consider this unlikely since no mechanism is known on how the piRNA pathway could detect tissues where a TE is active and then selectively repress TE expression in these tissues. Furthermore, a previous work in *D. melanogaster*, which investigated the P-element regulation in FACS-sorted primordial stem cells, also suggests that piRNAs inhibit splicing of the third P-element intron (IVS3) and not the expression level of the P-element [87]. Interestingly, this regulation of TE activity by repression of splicing does not seem to be a general mechanism of piRNAs, as for some TEs the expression level is repressed by piRNAs (e.g. Burdock; [87]). Regulation of expression could be an important complementary strategy, as otherwise any TE without introns, such as Bari1 [12], might escape host control by the piRNA pathway. The question remains as to how the piRNA pathway determines whether to inhibit splicing or expression of a given TE family and whether these are exclusive regulation mechanism. For example, it would be interesting to identify whether a modified P-element, where IVS3 is removed, can be controlled by the piRNA pathway (e.g. by inhibiting P-element expression).

On the whole, we find that the invasion dynamics of the P-element as well as the host response to the invasion are similar among *D. erecta, D. melanogaster* and *D. simulans*. However, unlike the described P-element invasions in other species [47, 48, 91], we found that the P-element invasion in *D. erecta* cannot be stopped in one replicate (R2). We think that such a replicate offers a unique opportunity to gain insights into the establishment of host control over an invading TE.

### A revised model for the silencing of an invading TE

According to the prevailing theory, the trap model, an invading TE is silenced when one copy of the TE jumps into a piRNA cluster, which then triggers the production of piRNAs which silence all TE copies in trans (fig. 7; [4, 59, 98, 27, 97]). In this model, the critical event that establishes host control over the invading TE is an insertion of the invading TE in a piRNA cluster (fig. 7). It is thought that a cluster insertion eventually activates the entire piRNA pathway, including the ping-pong cycle and the generation of Zuc-dependent trail piRNAs. For several reasons our results are incompatible with this notion. First we found that a cluster insertion is insufficient alone to trigger activation of the piRNA pathway. In replicate 2, where the P-element invasion escaped host control, we found several P-element insertions within piRNA clusters already at around generation 20 (supplementary table S9) and many more by generation 51. Two cluster insertions are of high quality and supported by multiple long-reads (fig. 2). It may be argued that these clusters insertions are not fully functional in the germline. For example the level of piRNA expression in the germline might be too low. This however raises the difficult question of which properties piRNA clusters require such that a single insertion into the cluster can establish host control over the invading TE. Such clusters could, for example, require a certain length, piRNA expression level, H3K9me3 modificats, or *Rhino* and *Kipfler* binding properties [19, 3, 42]. It is even possible that the capacity to silence TE invasion varies within a piRNA cluster. Ideally, to address this question it will be necessary to insert an artificial sequence into many diverse positions in various piRNA clusters and test if the insertion represses a reporter (e.g similarly to Luo et al. [57] and Josse et al. [35]). Our reciprocal crosses among replicates also support our conclusion that differences in the quality of cluster insertions among replicates are insufficient to explain why the invasion was silenced in some replicates but not in another, as the offspring of reciprocal crosses has largely the same genome (fig. 6). Nevertheless, the cluster insertions may contribute to the generation of the very first piRNAs against an invading TE. These initial piRNAs may be a necessary, albeit insufficient, precondition for establishing host control over an invading TE (fig. 7). Secondly, our densely sampled small RNA data revealed that siRNAs against the P-element emerged prior to piRNAs, which supports a recent finding by Luo et al. [57]. The authors propose that maternally transmitted siRNAs may initiate the conversion of a proto-cluster (e.g. a repetitive region with a P-element insertion) into piRNA producing locus. Transposon invasions may lead to the generation of sense and antisense transcripts that form dsRNA, which is then further processed by Dicer-2 into siRNAs [16, 57]. Given that sense transcripts are not the limiting factor (supplementary fig. S20), the generation of a few antisense transcripts for an invading TE could trigger the generation of siRNAs which, in turn, might initiate silencing of the invading TE. Antisense transcripts may be quite abundant. For example, we found that about 7% of all P-element transcripts are antisense. Unlike a random insertion into a piRNA cluster, which is likely a rare and highly stochastic event, the generation of siRNAs against an invading TE may be a much more predictable and repeatable event. Due to this high abundance of antisense transcripts formation of siRNAs may happen very early during a TE invasion. In agreement with this, siRNAs against the P-element emerged in all replicates after 10-15 generations (fig. 3). This siRNA-based mechanism for silencing an invading TE, is also able to distinguish self (genes) from non-self (TEs) as genes rarely generate antisense transcripts (supplementary fig. S20). However, we found that the generation of siRNAs against the P-element was also not sufficient to activate the piRNA pathway against the P-element, as large numbers of P-element siRNAs were found in replicate 2 but host-control was not established. siRNAs may thus lead to the generation of the first piRNAs against an invading TE which could again be a necessary, but insufficient, step for establishing host control over an invading TE (fig. 7). Thirdly we found that the activation of ping-pong is necessary to silence an invading TE. Although, some first piRNAs against the P-element were present in replicate 2 they were insufficient to stop the P-element invasion (fig. 1). The level of piRNAs in replicate 2 was found to be at a lower level than in other replicates, likely due to the absence of the ping-pong cycle. Although fully functional in all replicates, the ping-pong cycle was inactive for the P-element in replicate 2 for at least 67 generations (fig. 4, 6). Our data thus suggests that activation of the ping-pong cycle is necessary for stopping a TE invasion. Furthermore, activation of the ping-pong cycle seems to be independent event from the generation of the very first piRNAs against an invading TE. Activation of the ping-pong cycle is thus a crucial but stochastic event which does not necessarily occur once some first piRNAs against an invading TE emerged (fig. 7). The activation of the ping-pong cycle, subsequent to the generation of the first piRNAs against an invading TE, is also in agreement with previous work. Which has shown that during TE invasions, the size of the ping-pong signature (which is thought to reflect the abundance of piRNAs processed by the ping-pong cycle) is initially weak but increases with time [47, 48, 91]. Based on these findings we propose an updated model, named the crank-up model, surmising how the host establishes control over an invading TE (fig. 7). Likely due to the formation of dsRNA from both sense and antisense transcripts of a TE, siRNAs are produced which initiate the conversion of some TE insertion into piRNA producing loci and thus the production of the first piRNAs against the invading TE. TE insertions in piRNA clusters may contribute to the generation of the first piRNAs targeting an invading TE. Note, that we are deliberately avoiding the term primary piRNAs to prevent confusion with Piwi-bound piRNAs, which were initially thought to act upstream of the ping-pong pathway but are now believed to be generated downstream of the ping-pong cycle in a Zuc-dependent manner [83, 30, 17]. In our model, the critical step that establishes host control over an invading TE is the activation of the ping-pong cycle (fig. 7). This is in contrast to the trap model, where an insertion into a piRNA cluster is the critical step that activates host control. We call this view the crank-up model, in reference to old cars that needed to be cranked up laboriously to activate the engine but once started the inertia of the engine kept the combustion cycle running. Similarly, it seems difficult to activate the ping-pong cycle but once activated, maternally deposited piRNAs are thought to keep the ping-pong cycle running [52]. It is however unclear what triggers the activation of the ping-pong cycle. One possibility is that the concentration of piRNAs targeting a TE needs to exceed a certain threshold [66].

**Figure 7:**
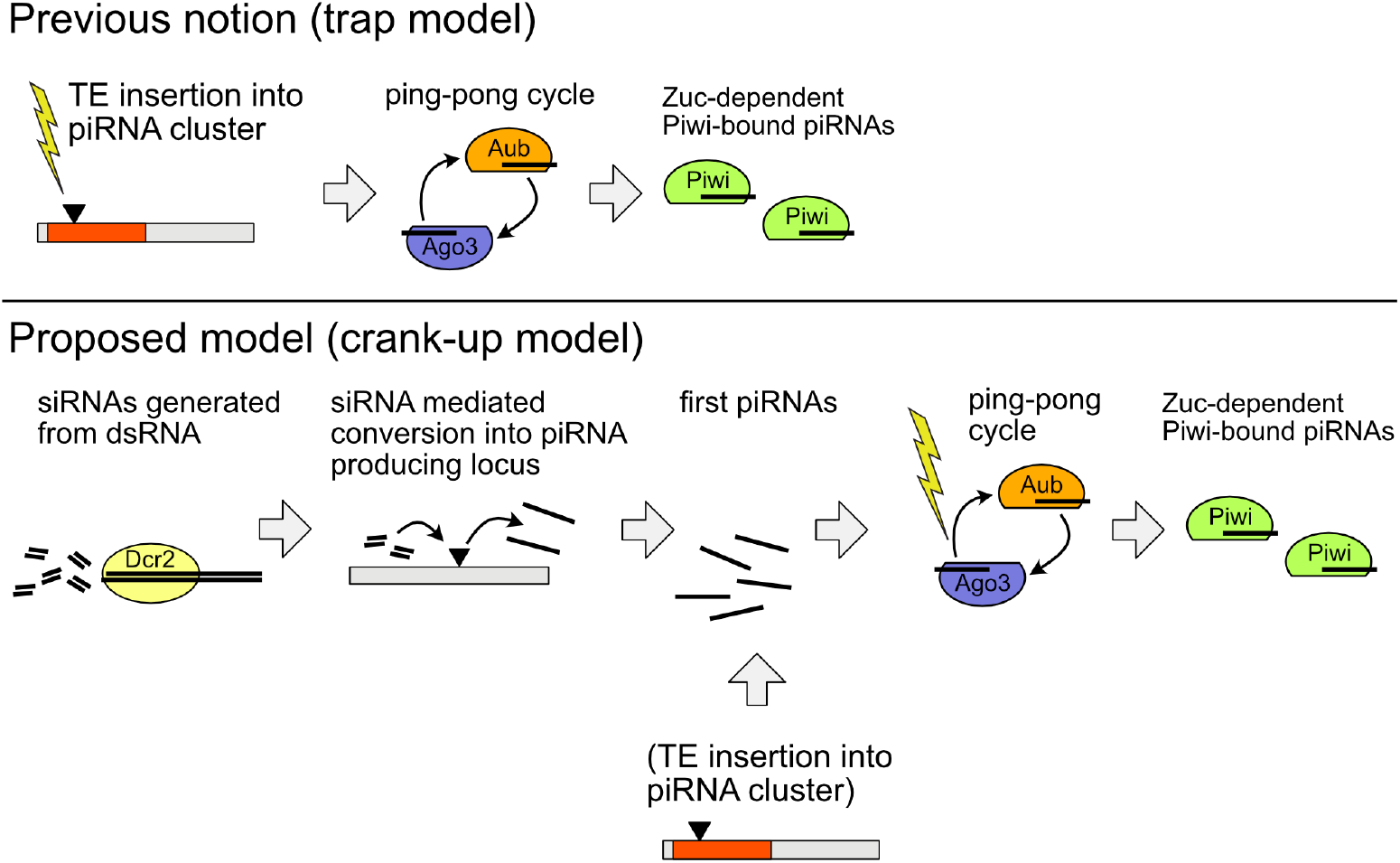
Two models for the silencing of TE invasions in the germline. The critical events, mainly responsible for establishing host silencing are indicated by yellow flashes. According to the prevailing view, i.e. the trap model, a TE insertion into a piRNA cluster is the critical event that establishes host control over an invading TE (yellow flash). A cluster insertion is thought to trigger the piRNA pathway, including the ping-pong cycle and the generation of Zuc-dependent trail piRNAs. In our updated model, the crank model, we propose that siRNAs initiate the emergence of piRNAs and that the critical event establishing host control over an invading TE is the activation of the ping-pong cycle (yellow flash). siRNAs are generated from Dcr2 mediated cleavage of dsRNA from TEs. The siRNAs then drive the conversion of a TE insertion into a piRNA producing locus, which leads to the emergence of the very fist piRNAs against a TE [57]. TE insertions in piRNA clusters may also contribute to the emergence of the first piRNAs. In this model, the first piRNAs against an TE are readily generated during each TE invasion. The critical step, ultimately responsible for triggering silencing of the TE, is the activation of the ping-pong cycle.

### Absence of ping-pong in replicate 2

It is an important open question as to why the ping-pong cycle could be activated in replicates 1 and 4 but not in replicate 2. By comparing replicates both with and without ping-pong against the P-element, we could rule out an influence of the expression of a key gene involved in triggering ping-pong (Chk2 [66]), the abundance of antisense transcripts against the P-element, the viral load [78] and the general expression level of known *D. erecta* transcripts (fig. 5; supplementary fig. S20; supplementary table S10). With the reciprocal crosses among replicates, we could further preclude an influence of the genomic composition, as the genomes are largely identical among the reciprocal crosses (fig. 6). As a consequence, we can rule out any influence of particularly potent cluster insertions, that may just be present in replicates 1 or 4 but not in replicate 2. Furthermore, the reciprocal crosses also rule out any influence of the prominent internal-deletion found solely in replicate 2 (i.e. the EP-element). These crosses instead suggest that maternally transmitted information, likely piRNAs (or siRNAs), is responsible for the absence of ping-pong in replicate 2 (fig. 6). The importance of maternally deposited piRNAs is also highlighted by a previous work which found that the ping-pong cycle is greatly enhanced if maternally deposited piRNAs are present [52]. We found some initial piRNAs against the P-element in replicate 2 but it is not clear if they are maternally deposited. Also, it is not clear if any maternally deposited piRNAs are sufficient to activate the ping-pong cycle. It is, for example, feasible that only piRNAs generated by the ping-pong cycle are effective in enhancing the ping-pong cycle in the next generation.

An alternative explanation is that the activation of the ping-pong cycle requires a certain minimum concentration of piRNAs Moon et al. [66] and that the piRNA concentration in replicate 2 is below this threshold. It is however unclear why this threshold concentration could not be reached in replicate 2, although this replicate has more piRNA cluster insertions and a higher P-element expression level (which will lead to more dsRNA and thus siRNAs) than the other two replicates (fig. 1C; supplementary table S9).

Since the ping-pong cycle is rapidly established within the first 20-25 generations in replicates 1 and 4 but not by generation 70 in replicate 2 (supplementary fig. S24), we believe that the small RNAs maternally deposited in replicate 2 might interfere with the activation of the ping-pong cycle. We speculate that the maternally deposited small RNAs in replicate 2 could have a peculiar composition that is not favorable for activating the ping-pong cycle. The emergence of such an unfavorable small RNA composition could be a highly stochastic process. It is however entirely unclear which features could characterize such a composition and which stochastic events drive experimental populations into a dead-end where the invading TE can not be controlled by the host. It is also unclear if the establishment of such unfavorable small RNAs is reversible, i.e. if the ping-pong cycle could be activated in future generations in replicate 2.

### Consequences for natural populations

Here we showed that host control over an invading TE could not be established in one experimental population. This raises the question as to whether this could also happen in natural populations. The invasion dynamics in natural populations may be different than in experimental populations. Whereas experimental populations are largely panmictic, more population subdivision is expected in natural populations. As a result, a TE invasion could be unchecked in a few local sub-populations but largely under host control in the rest. Migration between the sub-populations could then help to spread host-control to all sub-populations. An unchecked invasion, as described here, could thus solely be a threat to isolated populations or largely panmictic species. A related open question is how often unchecked TE invasions occur. In our three experimental populations, host defences were not established in a single replicate. In our previous work in *D. simulans*, the host defence was established in each of the six investigated experimental populations [47, 48]. Hence, it is our belief that unchecked TE invasions are rather rare (less than one in ten). An important question is what could happen to natural populations if host defence is not established. The TE will likely attain unusually high copy numbers in such (local) populations without host control For example, the experimental population without host control had many more P-element insertions (151 per haploid genome) than the populations where host control was established (27 *−* 37 per haploid genome). Furthermore, the fecundity was rapidly declining in the experimental population without host defence, likely due to the accumulating deleterious effects of many different P-element insertions. We could solely rescue this population by back-crossing the females to naïve males not having the P-element. Similarly, it was reported that very few replicate populations survived P-element invasions in *D. melanogaster* at 27°C, a temperature where the P-element is highly active [91]. This raises the possibility that TE invasions might drive natural populations, or possibly even species, to extinction. This problem could be especially severe when species are already stressed, e.g. due to climate change, or when an environmental change increases the activity of the invading TE. Rising temperatures will for example increase P-element activity[66]).

## Material and Methods

### Strains and Transformation

We obtained the reference strain of *D. erecta*, 14021-0224.01, from the UCSD Drosophila Species Stock Center (now the National Drosophila Species Stock Center, Cornell University). This strain is derived from a wild type strain, inbred for eight generations [21]. To introduce the P-element into *D. erecta*, the plasmid ppi25.1 - which contains a full length insertion of the canonical P-element (kindly provided by E. Kelleher; GenBank:X06779; [71]) - was injected into early embryos by Rainbow Transgenic Flies Inc (https://www.rainbowgene.com/; Camarillo, CA, USA). We established 12 lines by crossing the transformed adults (2 males and 3 females). The lines were screened for the presence of the P-element with PCR using four different primer pairs as described previously [33]. Out of the 12 lines tested, 7 contained the P-element (supplementary fig. S2). The transformed lines were maintained at 20°C (the P-element has a reduced activity at low temperatures [66, 47]) for 3 generations before setting up the experimental populations.

### Experimental populations

To establish the experimental populations, we crossed five males from five P-element lines with 5 naïve virgin females (in total we crossed 25 males with 25 females) and allowed them to mate for 3 days. After mating, we mixed the 50 crossed flies with 200 naïve *D. erecta* flies. The resulting 250 flies constitute the base population. For an overview of the experimental design see supplementary fig. S3We maintained 3 replicates of the experimental populations for 50 generations at 25°C at a census size of *N* = 250 using non-overlapping generations.

### Illumina sequencing of genomic DNA

Flies from each generation were stored in pure EtOH at -80C°C. At about each 10^*th*^ generation DNA was extracted from pools of 60 flies using a high salt extraction protocol [63]. Sequencing libraries were generated with about 330ng DNA and the NEBNext Ultra II FS DNA Library Prep Kit in combination with NEBNext Multiplex Oligos (New England Biolabs, Ipswich, MA, USA). We sequenced 2×125bp reads with an Illumina HiSeq 2500 at the VBCF-NGS facility (https://www.viennabiocenter.org/vbcf/next-generation-sequencing/). For an overview of all sequenced genomic samples see supplementary table S1.

### Abundance and diversity of TE insertions

We estimated the abundance and diversity of the P-element with DeviaTE [92]. The short reads were aligned with BWASW (v0.7.17 [56]) to the consensus sequences of TEs in *D. melanogaster* (which contains the P-element [74]) and three single copy genes of *D. erecta*: trafficjam, rpl32 and rhino (from FlyBase release 2017 05). The abundance of TEs is estimated by normalizing the coverage of TEs to the coverage of the single copy genes. DeviaTE also provides the position and frequency of internal deletions and SNPs of the P-element.

To identify the genomic position and population frequency of P-element insertions, we used PoPoolationTE2 (v1.10.04 [46]). We trimmed reads to a size of 75bp at the 3’-end (which increases the inner distance and thus the accuracy of population frequency estimates with PoPoolationTE2) and aligned the reads with BWASW (v0.7.17 [56] as single-ends to a fasta file consisting of a long-read based assembly of *D. erecta* [41] and the sequence of the P-element (X06779.1 [71]). Using PoPoolationTE2 we restored the paired-end information (se2pe), generated a ppileup file (ppileup --map-qual 15), identified signatures of TE insertions (identifySignatures --mode separate, --min-count 1, --signature-window fix100), determined the strand of the insertions (updateStrand --map-qual 15 --max-disagreement 0.4), estimated the population frequencies of TE signatures (frequency), filtered TE signatures (filterSignatures filterSignatures --min-count 1 --max-otherte-count 1 --min-coverage 10 --max-structvar-count 1) and finally paired TE signatures to obtain a list of P-element insertions (pairupSignatures). The number of reads mapping to the P-element (rpm) was also estimated with PopoolationTE2 (stat-reads). We estimated the transposition rate as *u* = *−*1*−*(*n*_*k*+*t*_*/n*_*k*_)^(1*/t*)^, where *t* is the time (in generations) between two measurements of TE copy numbers *n*_*k*_ and *n*_*k*+*t*_ (either rpm or abundance normalized to single copy genes).

### RNA sequencing

We sequenced RNA either from whole female flies or ovaries. To obtain ovaries, we kept flies with an age between 7-15 days on sugar-agar supplemented with yeast for two days. Ovaries were dissected on PBS. We used 30 flies for the extractions of all samples except generation 10 where solely 10-15 flies were used. To obtain embryos, flies (*<*7 days old) were allowed that lay eggs for 30 minutes on apple-juice agar plates with yeast paste. Embryos were transferred to a fine net, washed in cold water and dechorionated with 50% bleach for 2 minutes. Embryos were washed twice with an wash buffer (140mM NaCl, 0.03% TritonX100) and frozen in liquid nitrogen. We added 200 *µ*l TRIzol^®^Reagent (Invitrogen, Carlsbad, CA), to all samples, ground them and added another 800 *µl* Trizol. After incubation for 10 minutes, samples were vorted and 200 *µl* Chloroform was added. After spinning 500 *µl* of the upper phase was transferred to new tubes. 550 *µl* of isopropanol was added and then incubated for 10 minutes. The samples were spun at 4°C. The RNA pellet was cleaned with freshly made 80% EtOH and then dissolved in Nuclease-free water (Solis BioDyne, Estonia). The concentrations of the RNA samples were measured using Qubit (Invitrogen, Carlsbad, CA).

Small RNA and RNA samples were sequenced using Fasteris (https://www.fasteris.com/en-us/. After 2S RNA depletion the small RNA samples were sequenced using Illumina NextSeq with a read length of 75bp. For an overview of all sequenced small RNA samples, see supplementary table S3. The RNA samples were treated with DNase before they were sequenced on the Illumina NovoSeq machine, with a read length of 2×100bp. For an overview of all sequenced RNA samples see supplementary table S4.

### Analysis of small RNA data

Adaptor sequences were removed with cutadapt (v2.6 [62]) and reads with a length between 18 and 35 nt were retained. We aligned small RNA reads to a fasta file containing the *D. erecta* tRNAs, miRNAs, mRNAs, snRNAs, snoRNAs, rRNAs (r1.3; http://flybase.org/) and the consensus sequences of TEs from *D. melanogaster* [v9.42; plus Mariner GenBank: M14653.1 [74]] using novoalign (v3.09.00; http://www.novocraft.com/ -F STDFQ -o SAM -o FullNW -r RANDOM). The abundance of different small RNAs, the distribution of piRNAs within the P-element, the length distribution of the piRNAs and the ping-pong signal were computed using previously described Python scripts [47]. The motifs of small RNAs were computed with a novel script (smallRNA-U-bias.py) and the R-package ggseqlogo [89].

To identify piRNA clusters, we mapped the small RNA data from naïve ovaries to a long-read assembly of *D. erecta* [41] with novoalign (see above). Unambiguously mapped reads with a size between 23 and 29nt were counted within bins of 500bp, normalized to the number of miRNAs (see above) and then piRNA clusters were identified with an algorithm based on local scores, as described previously [47, 48] (–threshold 10). Clusters smaller than 2000bp were ignored.

We estimated the abundance of viruses in the small RNA libraries by aligning the small RNA data to a collection of *Drosophila* viruses maintained by the Obbard-lab (https://obbard.bio.ed.ac.uk/index.html; June 22; [69, 90] with novoalign (v3.03.02; -F STDFQ -o SAM -o FullNW -r RANDOM). We quantified the abundance of viruses using a custom script (viral-expression.py).

Statistical analysis and visualization of the data were performed in R [75].

### Analysis of expression data

We aligned the RNA data with gsnap (version 2014-10-22; [96]; -N 1) to a fasta file consisting of the transcripts of *D. erecta* (r1.3; FlyBase) and the consensus sequences of TEs in *D. melanogaster* [74]. The coverage and splicing level of the P-element was visualized in R, based on the results of two scripts, which normalized the samples to a million mapped reads (mRNA-coverage-senseantisense.py, mRNA-splicing-senseantisense.py). The raw counts of reads mapping to all transcripts and TEs was estimated with a separate script (mRNA-expression.py). Significant differences in expression levels were identified with edgeR based on the raw counts (v3.38.1; glmQLFit test; [77]). Volcano plots were generated in ggplot2 [94] using the results of edgeR. The orthologous sequence of the *D. melanogaster* gene Chk2 was identified by aligning NM 001259171.2 to *D. erecta* transcripts with BWASW (v0.7.17; [56]).

### Long-read sequencing

To identify P-element insertions in piRNA clusters, we employed Oxford Nanopore long-read sequencing. We used 60 flies to extract high molecular weight DNA with a phenol-chloroform extraction method [80]. We used 1*µg* DNA to prepare long-read libraries with the Ligation Sequencing kit SQK-LSK109 (Oxford Nanopore Technologies; Oxford). Libraries were run on R9 flow cells for 72 hours.

For an overview of all sequenced long-read samples, see supplementary table S7.

The long-reads were aligned with minimap2 (v2.10-r761; -x map-ont -Y -c) [55] to a fasta file containing the long-read assembly of *D. erecta* and the sequence of the P-element (see above). We identified reads supporting P-element insertions (Pele-insertion-finder.py) in the resulting paf-files. This script also resolved duplex-reads - i.e. reads where parts of a genomic region are sequenced twice in a tandem arrangement with one copy being reverse complemented - by truncating the read at the first base of the reverse complemented sequence. Next, we filtered reads for insertions in piRNA clusters (find-lr-bedinsertion.py; using the piRNA clusters identified above). We identified the final set of P-element insertions in piRNA clusters by grouping reads supporting a cluster insertion at similar positions (distance of less than 20nt; group-cluster-insertions.py --pos-tol 20).

To estimate the population frequency of different P-element variants, we filtered for ONT long-reads supporting either the insertion of a full-length (FL) P-element or an EP-element (filter-FL-EP.py; for the EP-element, the bases 827-2375 are deleted). We linked the identity of the P-element insertion inferred from ONT long reads to the population frequency estimates obtained with PoPoolationTE2 (see above). For each P-element insertion we used the frequency estimate of the nearest insertions (popfreq-FL-EP.py). Insertions without proximal frequency estimate were ignored (the maximum distance was 20nt).

### Crosses among replicates

At generation 67 - 70 we performed reciprocal crosses between flies from replicate 2 with flies from replicates 1 and 4. We crossed 15 virgin females from replicate 2 with males from replicates 1 and 4 and vice versa. For each cross we set up 3 subreplicates. The parental females and the F1 females were used for RNA extraction and small RNA sequencing. For an overview of the small RNA data see supplementary table S3.

### Gonadal dysgenesis assays

About 150-200 flies from every fifth generation were allowed to lay eggs at 29°C for 3 days. The progeny was kept at 29°C until the pupal stage and was than moved to 25 °C. Eclosed flies were transferred to apple-juice agar plates with live yeast paste and kept at 25°C for three days. The flies were dissected in 1x PBS solution and the size of the ovaries was scored using the following classification: clearly visible ovarioles or eggs (clear), ovarioles barely visible, atrophy in one ovary (weak), no ovarioles or eggs could be detected (absent). The percentage of dysgenic ovaries was computed as 100 *∗* (*absent* + (*weak/*2))*/*(*clear* + *weak* + *absent*)

## Data availability

All scripts used in this work are available at https://sourceforge.net/p/te-tools/code/HEAD/tree/ (folders “ere” and “longread”).

All data have been made publicly available at the European Nucleotide Archive (xxx).

## Author Contributions

RK conceived this work. DS performed the experiments and prepared sequencing libraries. FW extracted small RNAs from embryos and contributed to writing. RK and DS analyzed the data. RK and DS wrote the manuscript.

## Acknowledgement

We thank Erin Kelleher for discussions and providing the plasmid ppi25.1. We thank Kirsten-André Senti for advice and helpful discussions. We are grateful to Matthew Beaumont for valuable comments on the manuscript. We thank Marie Fablet, Almorò Scarpa, Stephane Ronsserray and Christian Schlöotterer for comments. We thank all members of the Institute of Population Genetics for feedback and support. This work was supported by an Austrian Science Foundation (FWF) grant (P35093) to RK.

