## supplement for "P-element invasions in *Drosophila erecta* shed light on the establishment of host control over a transposable element"

December 2022

### Supplementary figures

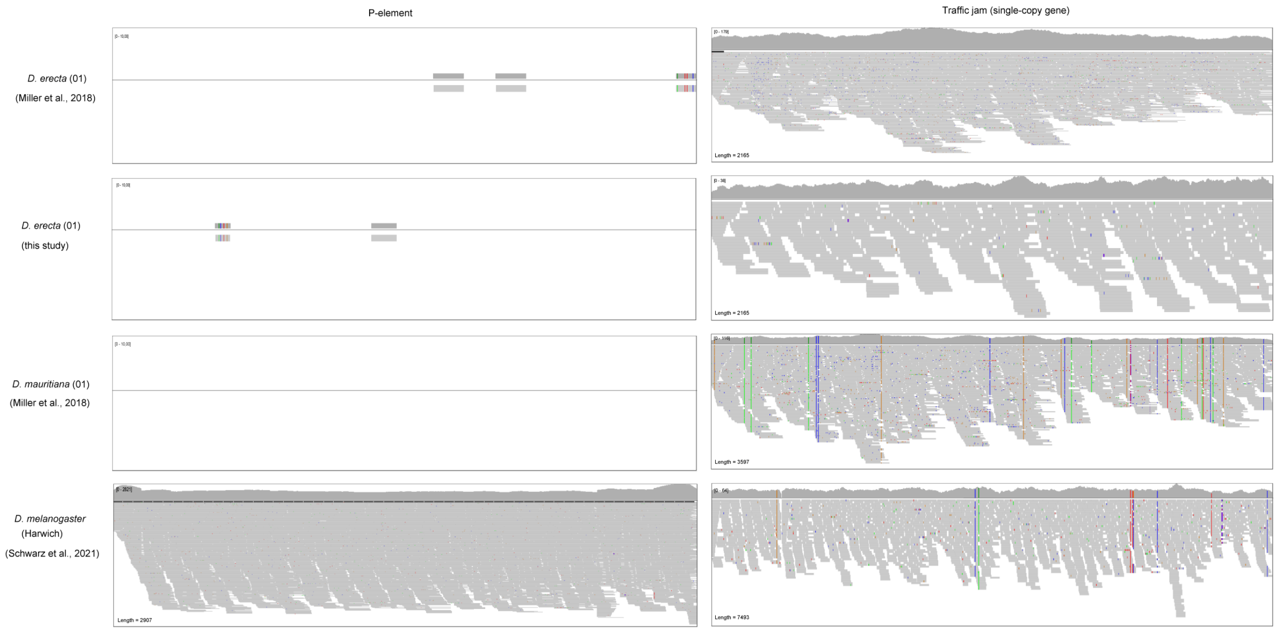


Figure 1: Abundance of reads aligning to the P-element and the single-copy gene *traffic jam* in *D. erecta* (strain 01), *D. mauritiana* (strain 01) and *D. melanogaster* (Harwich). Previous works showed that the P-element is present in the *D. melanogaster* strain Harwich but absent in *D. mauritiana* [Kofler et al., 2015, Brookfield et al., 1984, Srivastav et al., 2019]. In agreement with this many reads align to the P-element in *D. melanogaster* but none in *D. mauritiana*. Although many reads align to the single-copy gene solely 2-3 reads align to the P-element in *D. erecta*, suggesting that the P-element is absent in *D. erecta*.

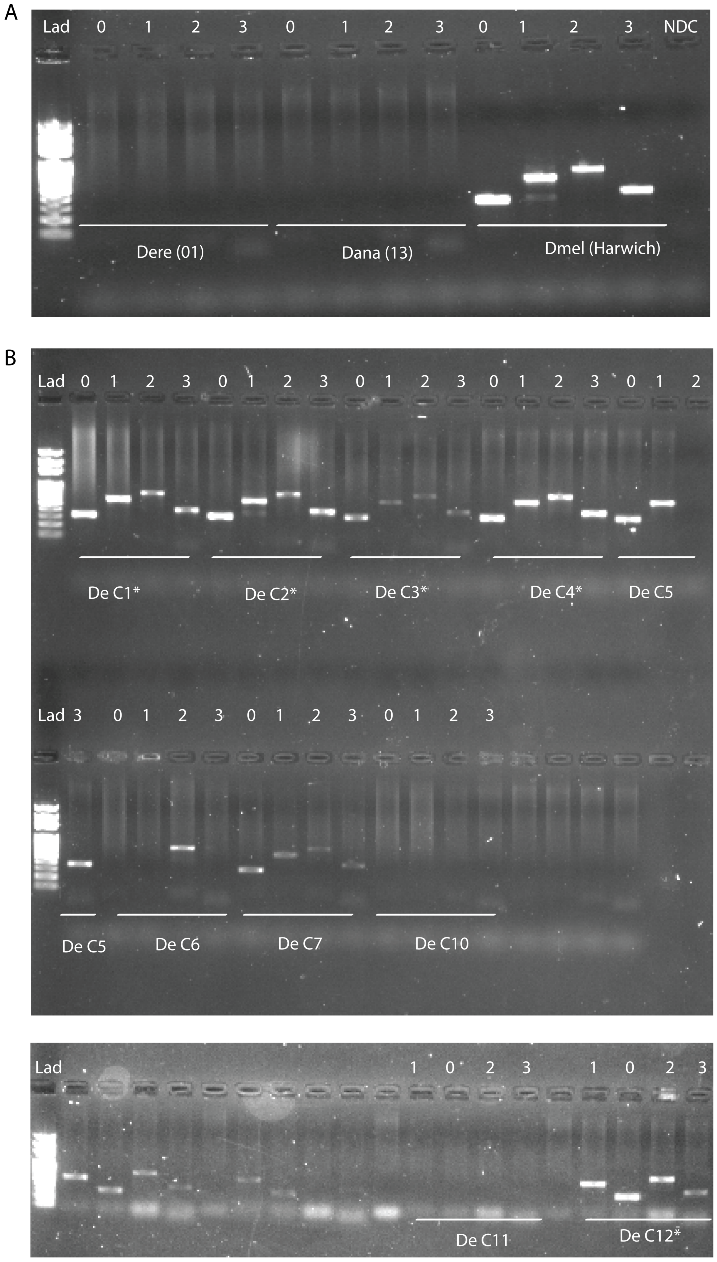


Figure 2: Presence of the P-element in different *D. erecta* lines. We tested each line (species) with four different primer pairs (number on top), where a primer pair was designed for each of the four ORFs of the P-element [Hill et al., 2016]. A) The P-element is absent in the *D. erecta* strain 01. The *D. melanogaster* strain Harwich was used as a positive control and *D. ananassae* as negative control [Daniels et al., 1990, Srivastav et al., 2019]. B) Presence of the P-element in 12 transformed *D. erecta* lines. The plasmid ppi25.1 was microinjected into naive *D. erecta* 01 embryos. Surviving G0 flies were mated and 12 lines were established. The lines used for setting up the experimental populations are marked with a star (*).

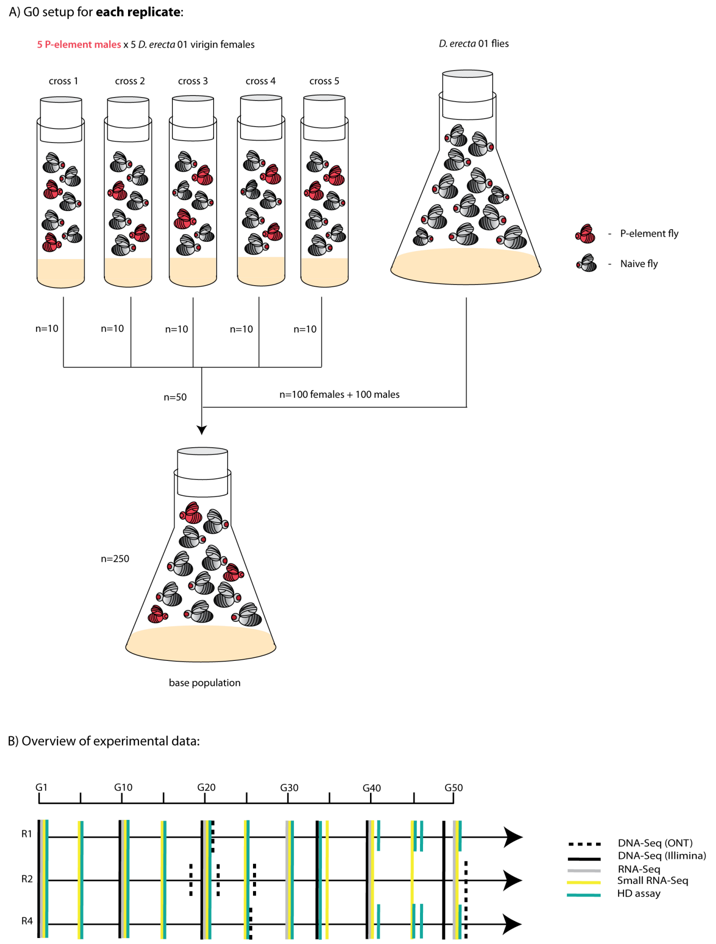


Figure 3: Overview of our experimental design. A) Strategy to setup the base population B) Overview of the experimental data generated in this work.

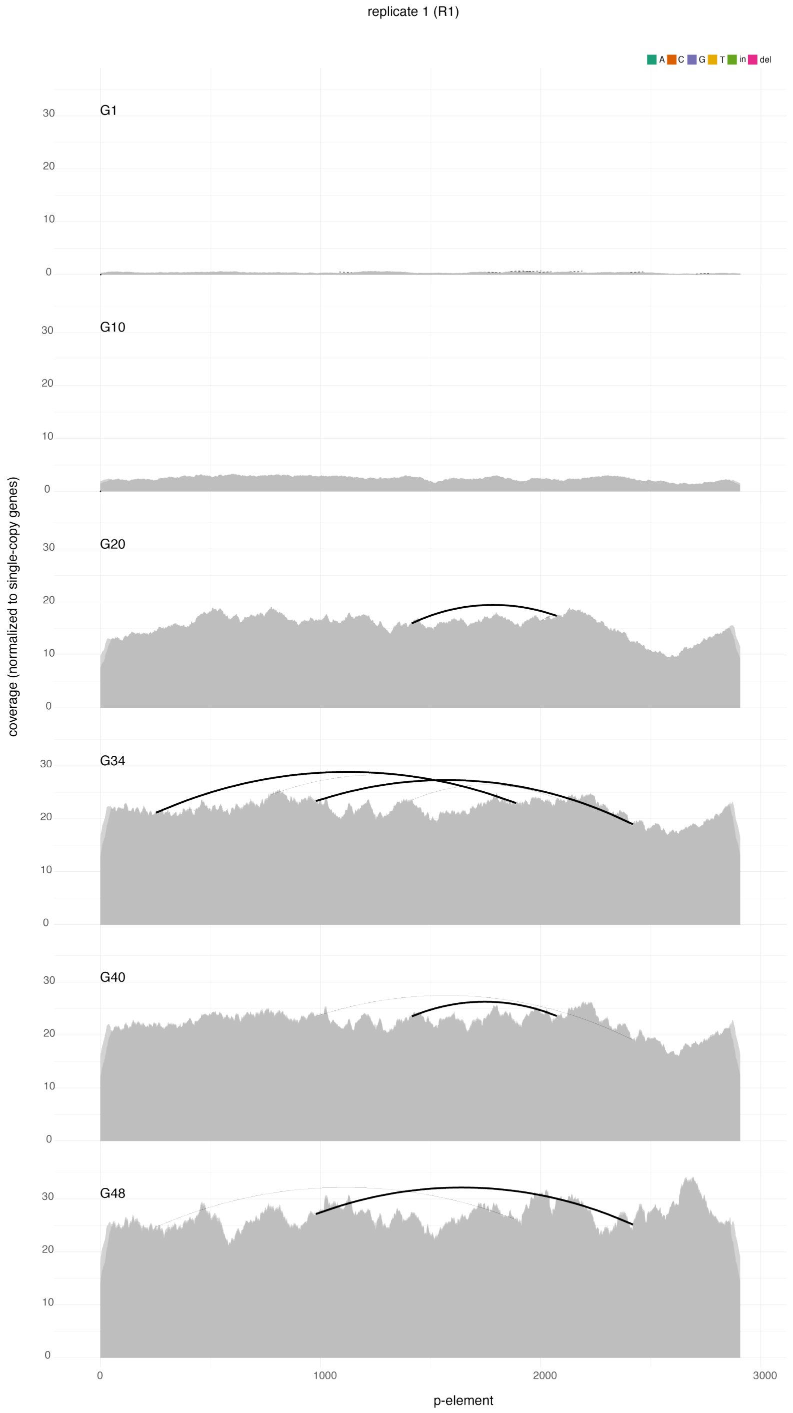


Figure 4: Abundance and diversity of the P-element during the invasion in replicate 1 visualized with DeviaTE [Weilguny and Kofler, 2019]. Single-nucleotide polymorphisms (SNPs) and small internal deletions (indels) are shown as colored lines. The absence of colored lines highlights that the P-element has no SNPs or solely SNPs segregating at a very low frequency. Large internal deletions are shown as black arcs.

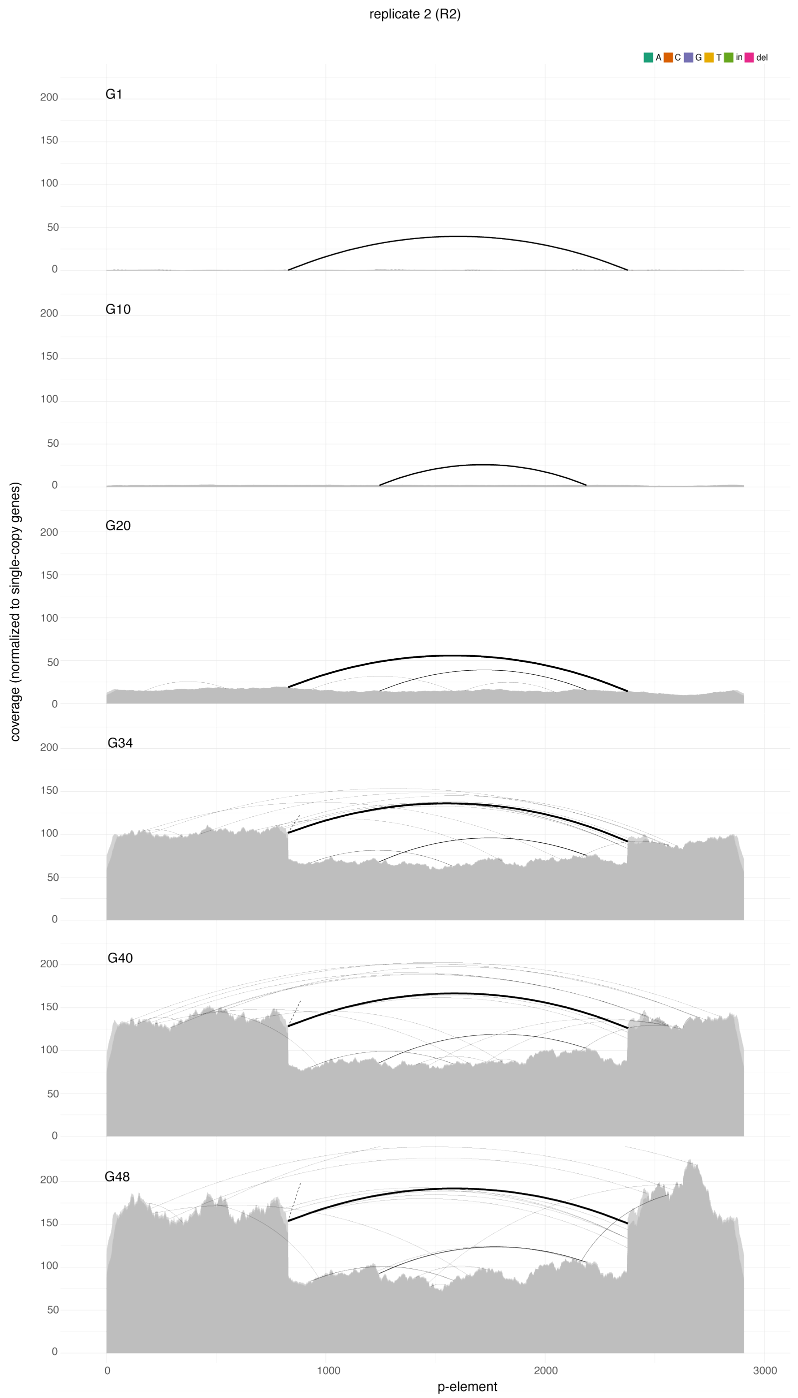


Figure 5: Abundance and diversity of the P-element during the invasion in replicate 2.

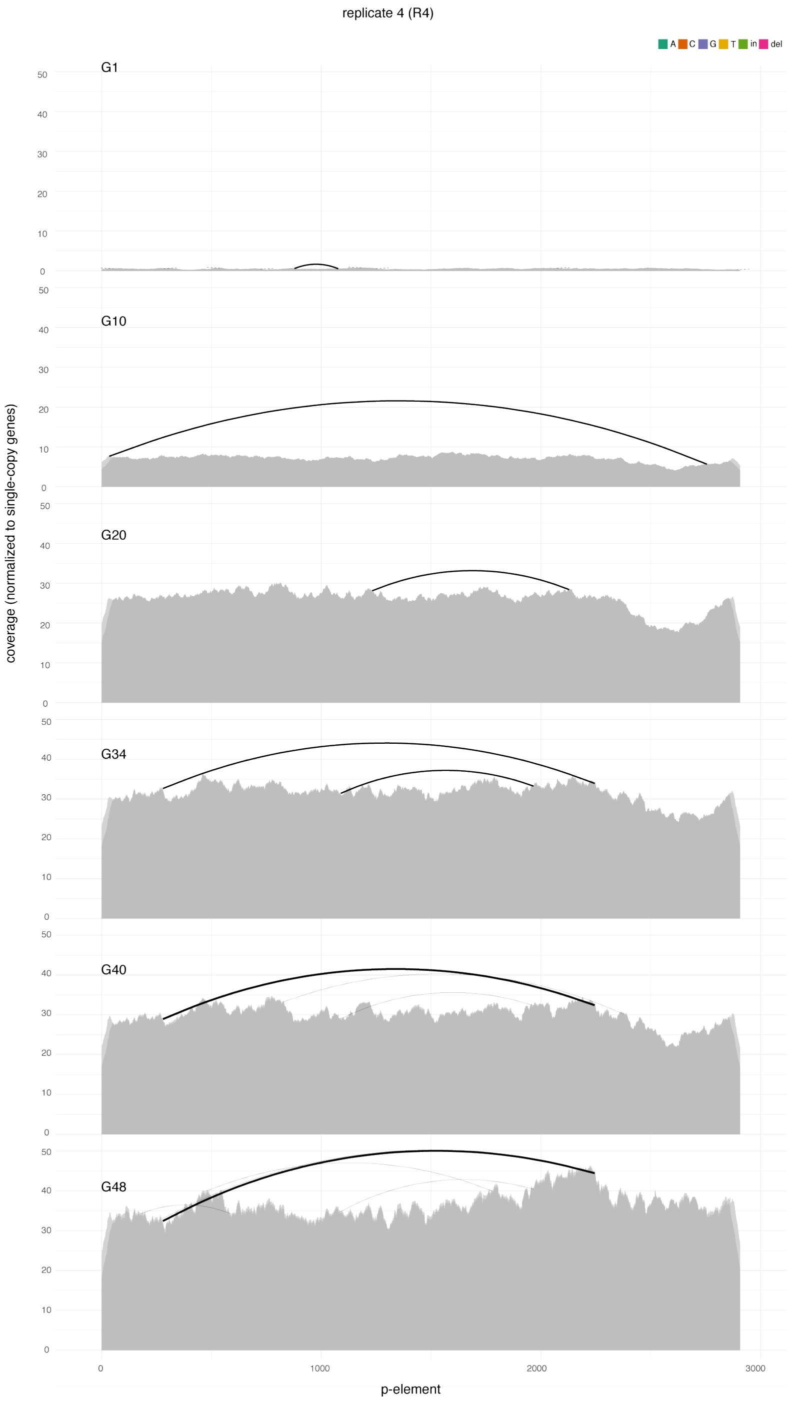


Figure 6: Abundance and diversity of the P-element during the invasion in replicate 4.

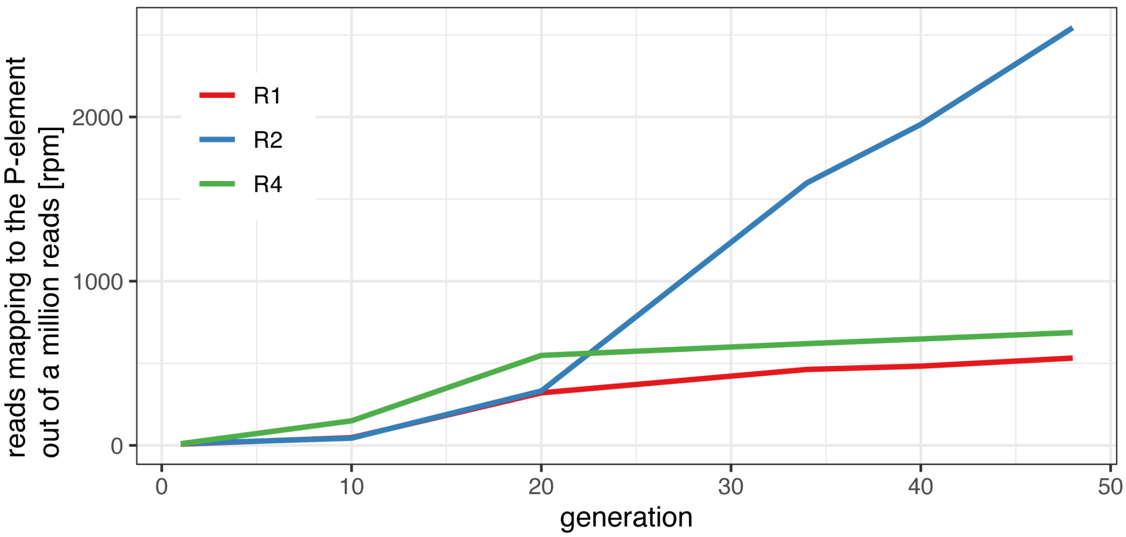


Figure 7: Abundance of P-element insertions during the invasion in reads mapping to the P-element out of a million reads (reader per million; rpm). Data are shown for three replicates (R1, R2, R4).

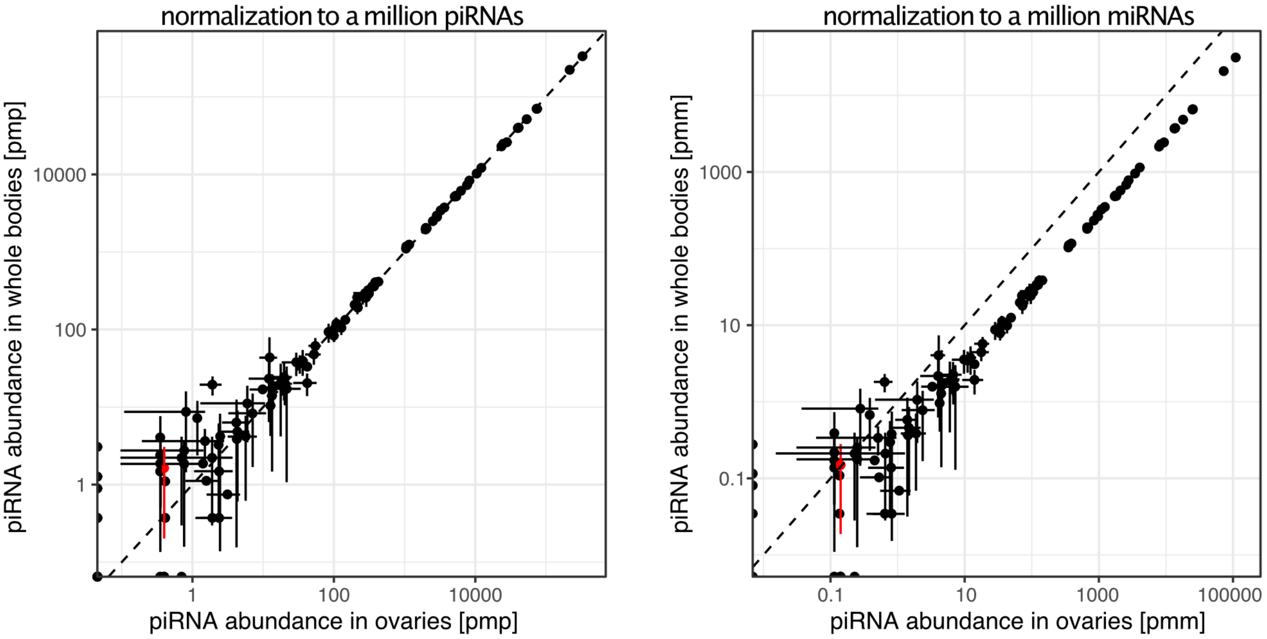


Figure 8: Effect of the normalization method on the piRNA abundance in ovaries and whole female flies. The abundance of piRNAs for each TE family (each dot is a TE family) was either normalized to a million piRNAs (left) or to a million miRNAs (right). We sequenced three replicates from ovaries and whole female flies of naive *D. erecta* flies (not having the P-element). For each replicate RNA was extracted from different flies. The error bars show the standard deviation among the three replicates and the dashed lines indicate the diagonal, i.e. the perfect correlation. The abundance of piRNAs complementary to the P-element is shown in red. Note that sequencing of small RNAs from ovaries and whole female flies yield similar results if the normalization to a million piRNAs is used. pmp piRNAs per million piRNAs, pmm piRNAs per million miRNAs

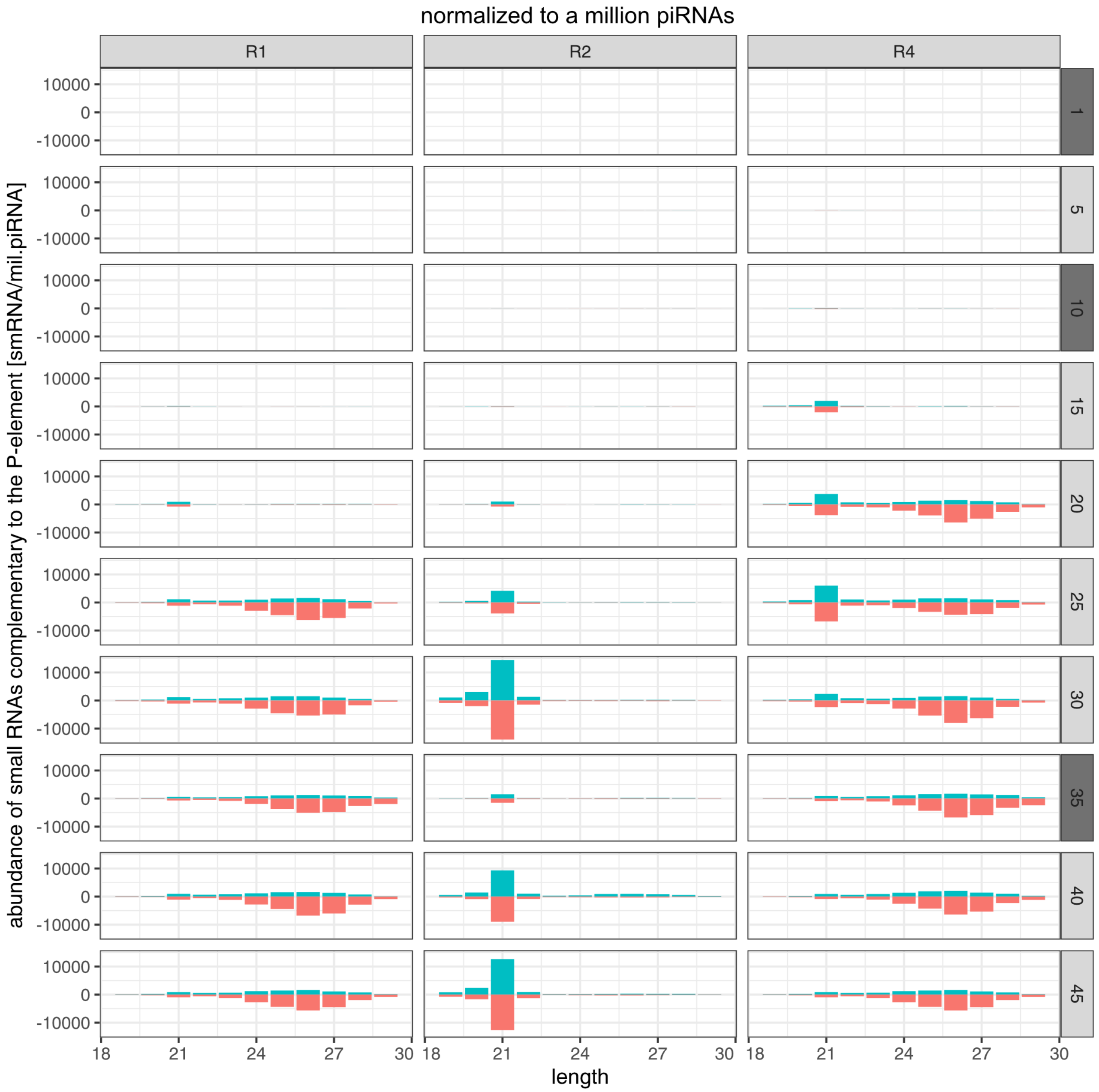


Figure 9: Length distribution of small RNAs mapping to the P-element. The abundance of small RNAs was normalized to 1 million piRNAs (smRNAs/mil.piRNA). Data are shown for three replicates (top panels) and several generations during the invasion (right panel). Sense RNAs are on the positive y-axis and antisense RNAs on the negative y-axis. small RNA data were either generated for whole bodies of female flies (light grey panels) or ovaries (dark grey panels).

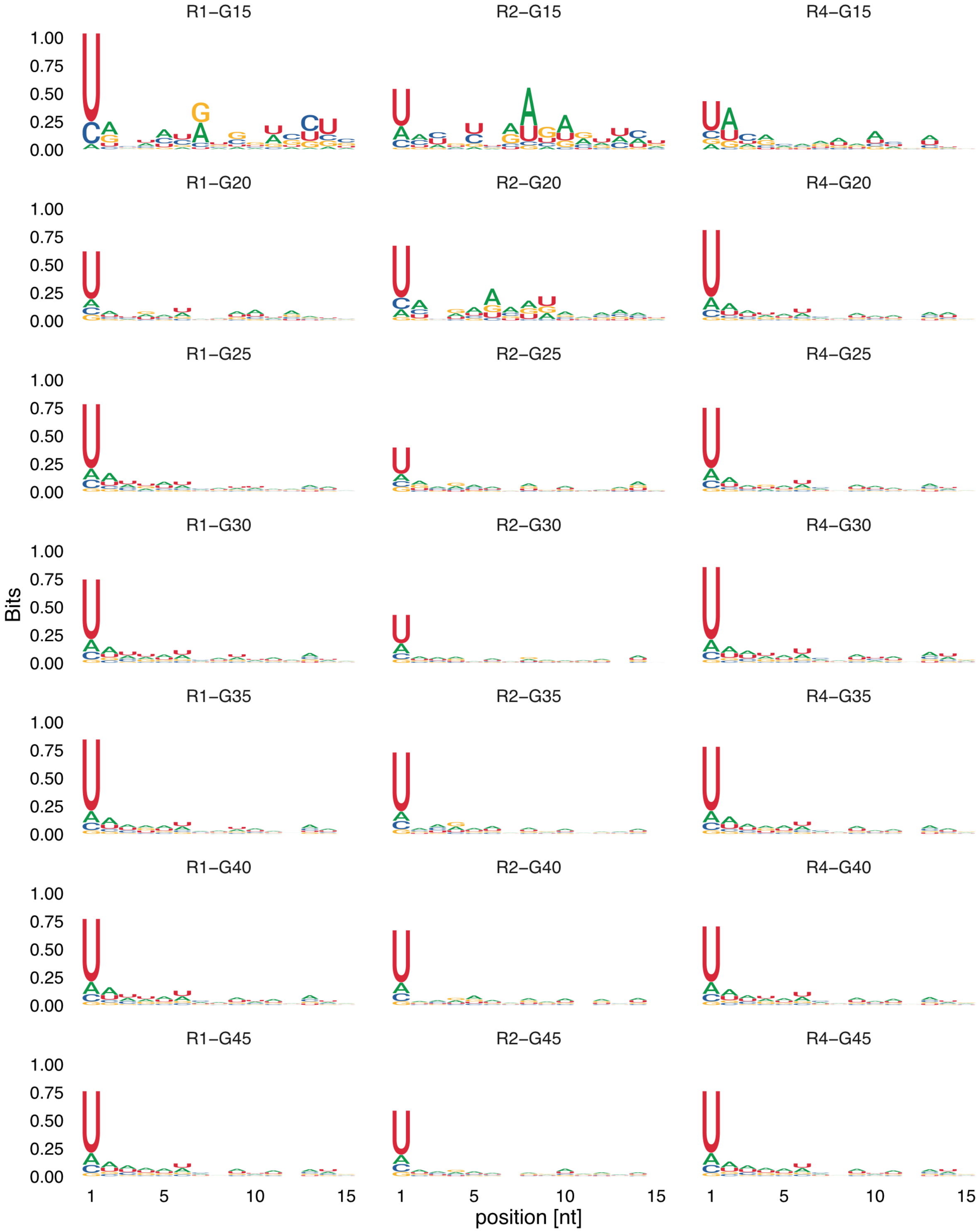


Figure 10: DNA motifs of P-element piRNAs (23-29nt). Data are shown for three replicates (R1, R2, R4) and multiple generations during the invasion (G15-G45). Prior to generation 15 the abundance of piRNAs was too small for computing motifs.

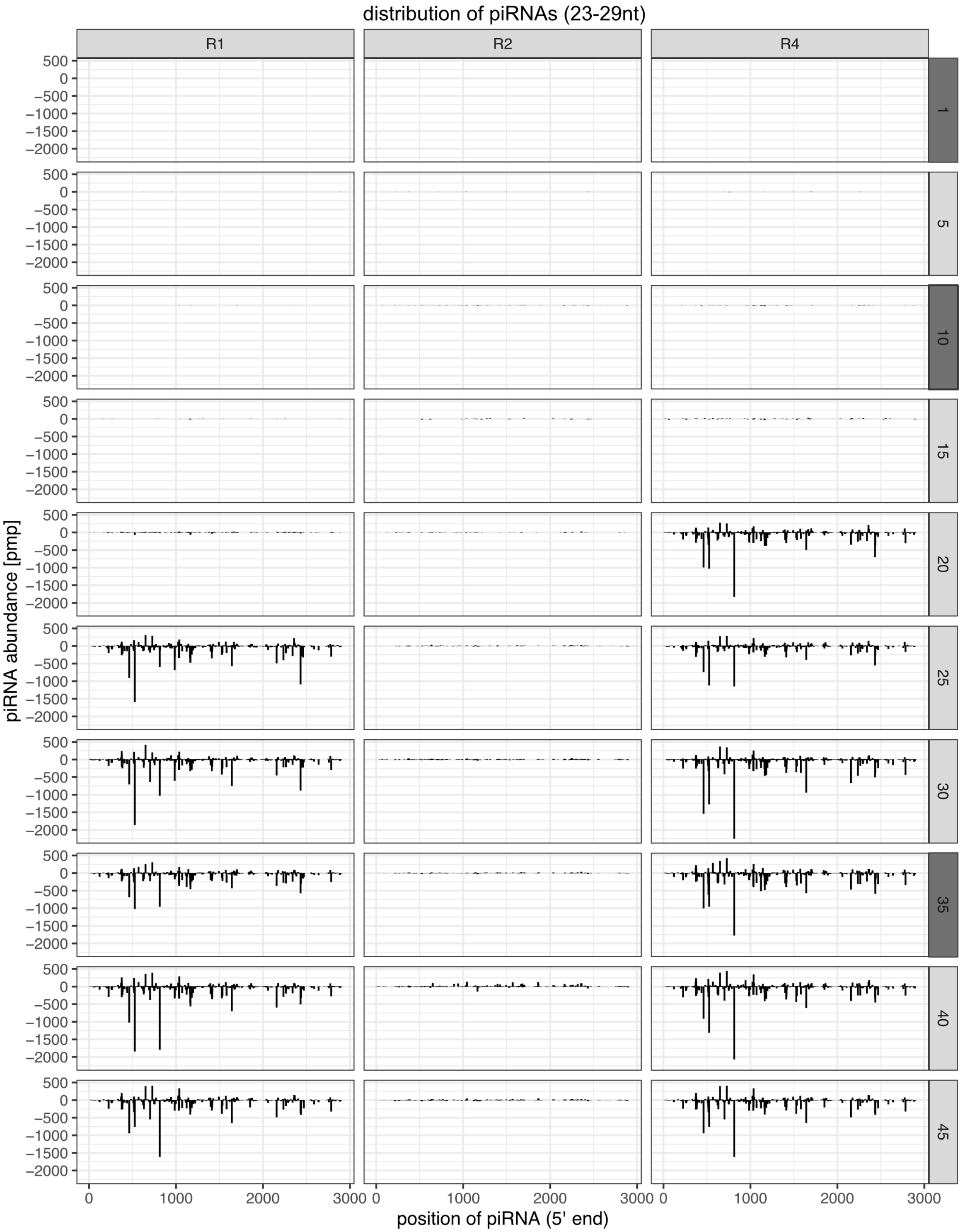


Figure 11: Distribution of piRNAs (23-29nt) along the P-element. Only the 5' positions of piRNAs are shown and the piRNA abundance is normalized to one million piRNAs. Replicates are at the top panel and the generations at the right panel. Sense piRNAs are shown on the positive y-axis and antisense piRNAs on the negative y-axis. small RNA data were either generated for whole bodies of female flies (light grey panels) or ovaries (dark grey panels).

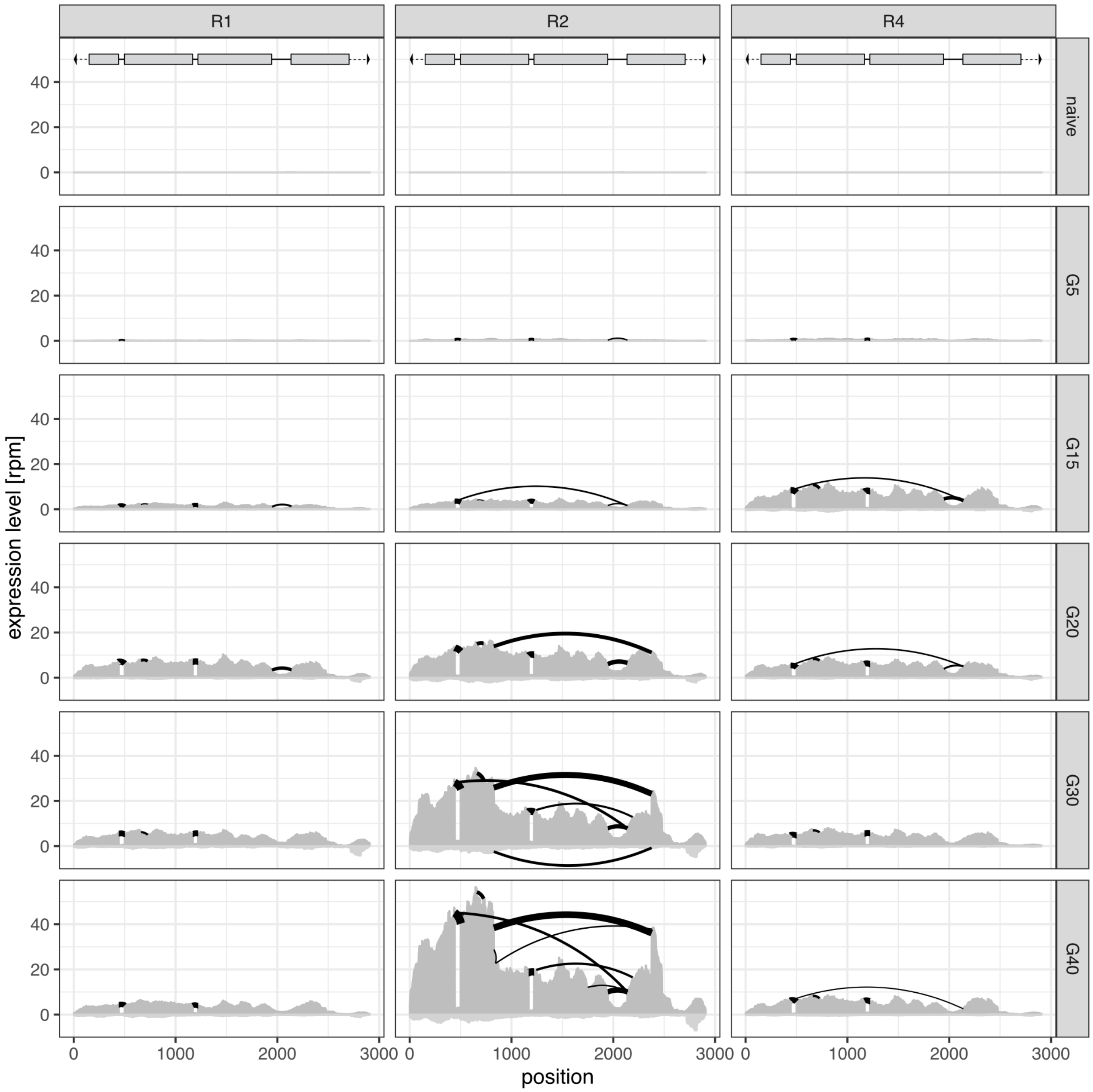


Figure 12: Sashimi plots showing the expression level and splicing of the P-element in female flies (whole body). Data are shown for different replicates (top panel) and generations during the invasion (right panel). Both the expression and splicing level (width of the black arcs) were normalized to a million mapped reads. Sense expression and gaps of sense transcripts (splicing or internal deletions) are shown on the positive y-axis whereas antisense expression and gaps of antisense transcripts are shown on the negative y-axis. The structure of the P-element is shown at the top, where terminal inverted repeats (TIRs) (black triangles), the four exons (grey rectangles) and introns (black lines) are shown.

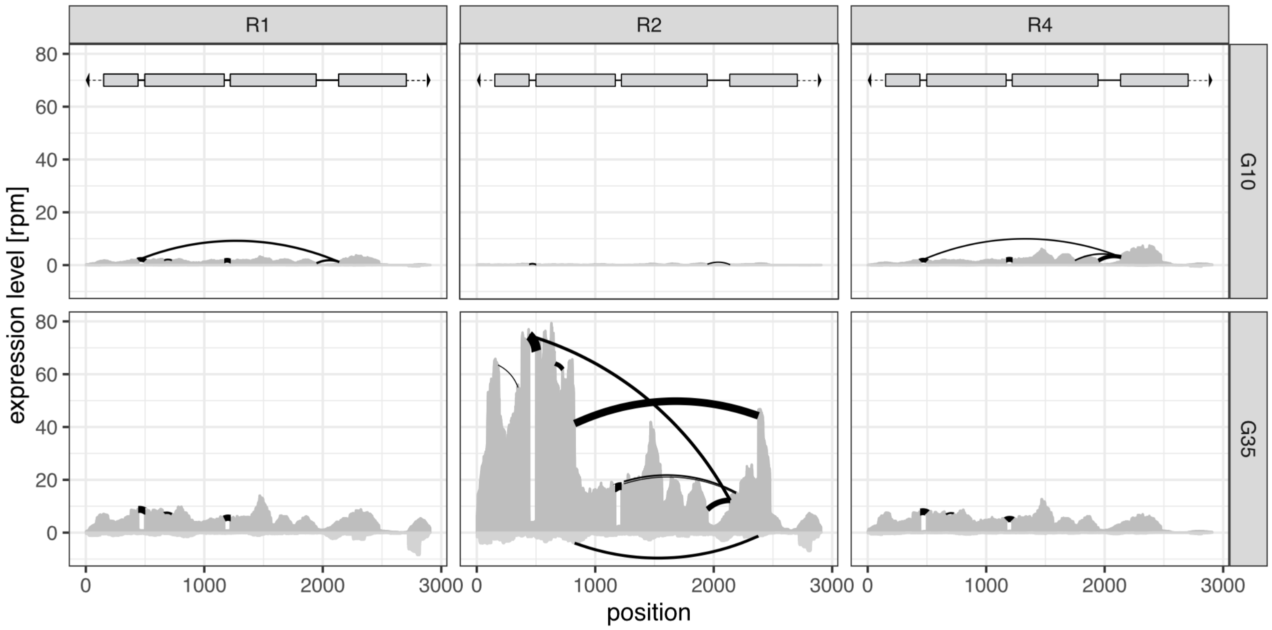


Figure 13: Sashimi plots showing the expression level and splicing of the P-element in ovaries. Data are shown for different replicates (top panel) and generations during the invasion (right panel). Both the expression and splicing level (width of the black arcs) were normalized to a million mapped reads. Sense expression and gaps of sense transcripts (splicing or internal deletions) are shown on the positive y-axis whereas antisense expression and gaps of antisense transcripts is shown on the negative y-axis. The structure of the P-element is shown at the top, where TIRs (black triangles), the four exons (grey rectangles) and introns (black lines) are shown.

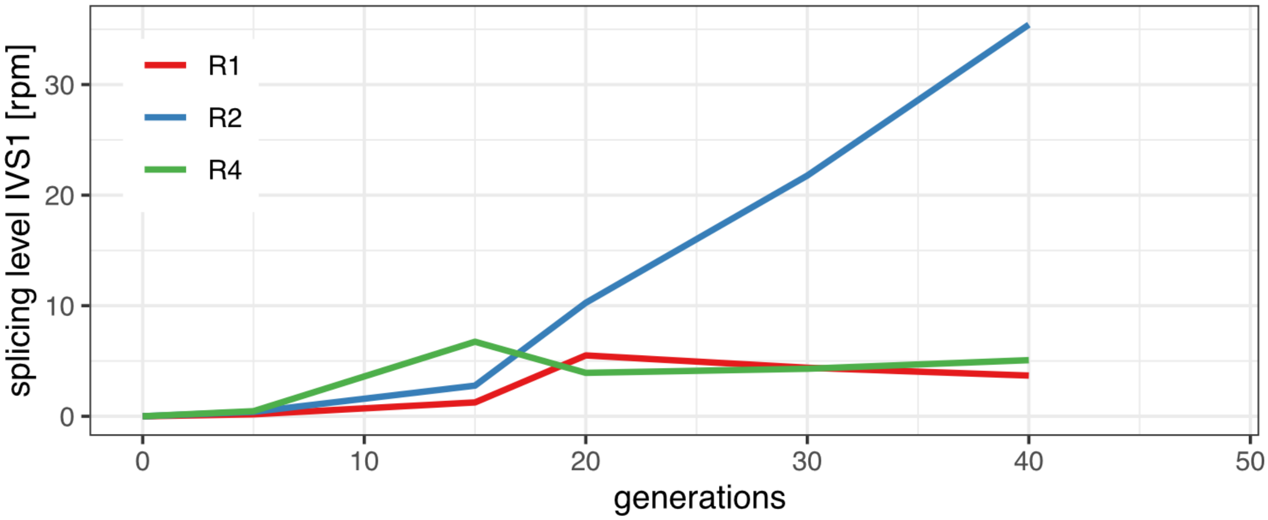


Figure 14: Splicing level of the first intron of the the P-element (IVS1) in the experimental populations.

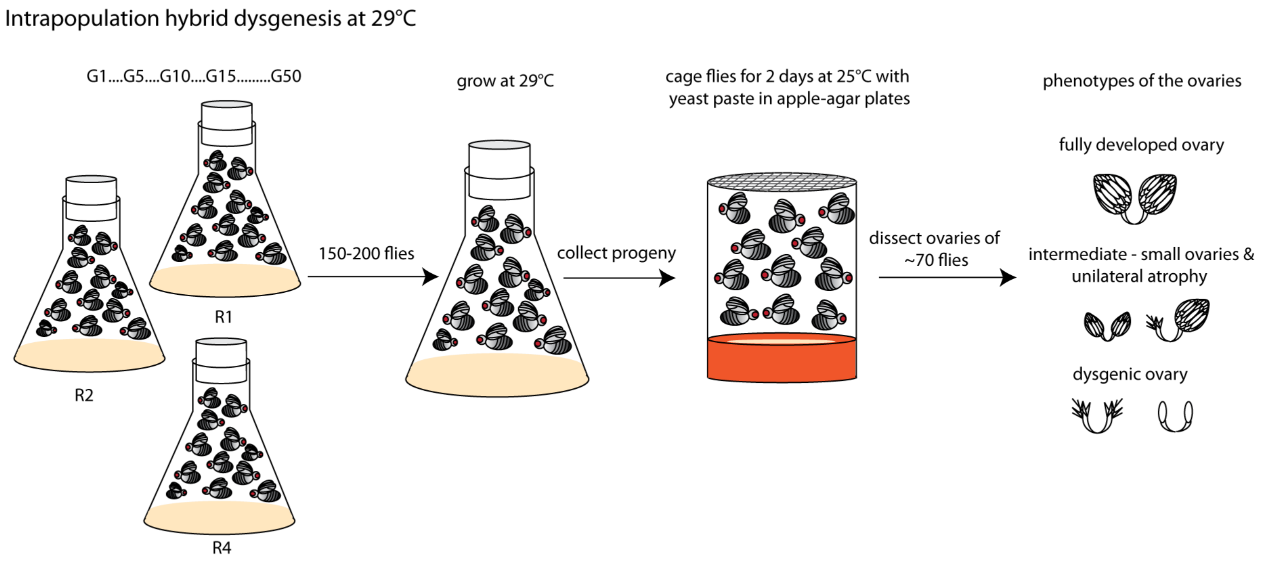


Figure 15: Overview of the gonadal dysgenesis assays performed for the three replicate populations (R1, R2, R4).

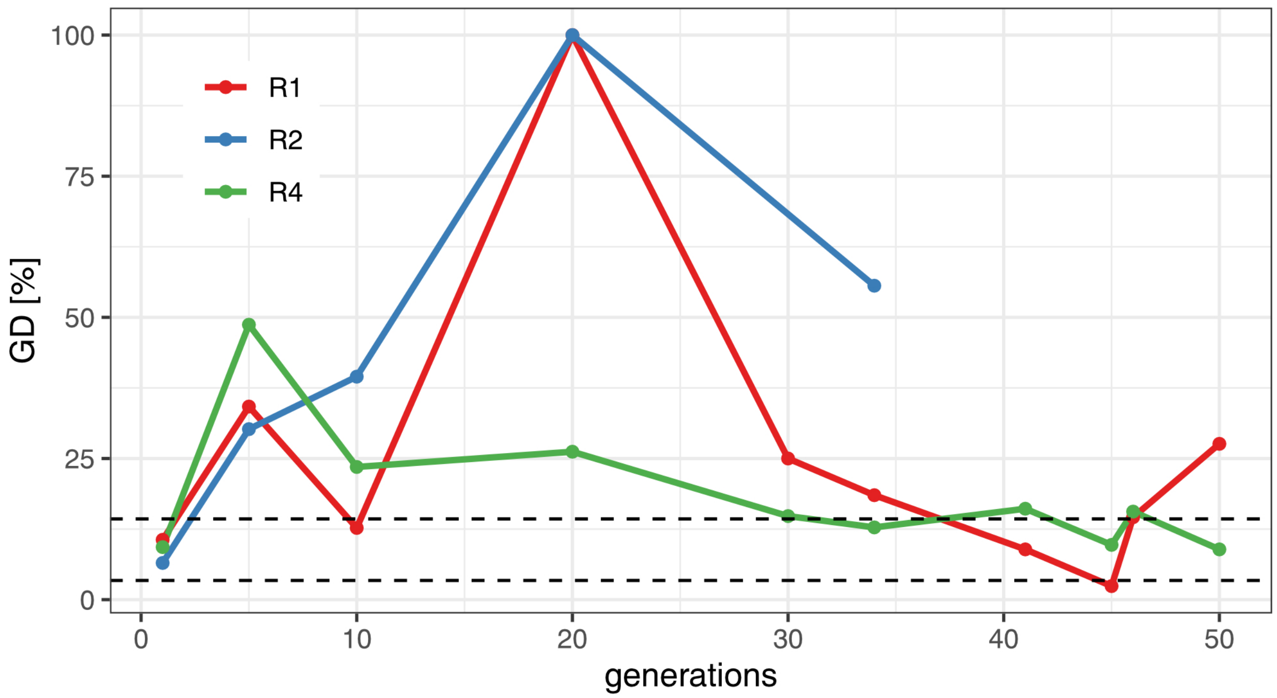


Figure 16: Extend of gonadal dysgenesis (GD) during the experiment. We allowed the experimental populations to lay eggs for 2-3 days, the eggs were reared at 29°C and the percentage of atrophied ovaries was estimated for all three replicates (R1-R4). Dashed lines indicate the range of GD observed for naive flies not having the P-element. Note that insufficient numbers of flies eclosed for replicate 2 after generation 34.

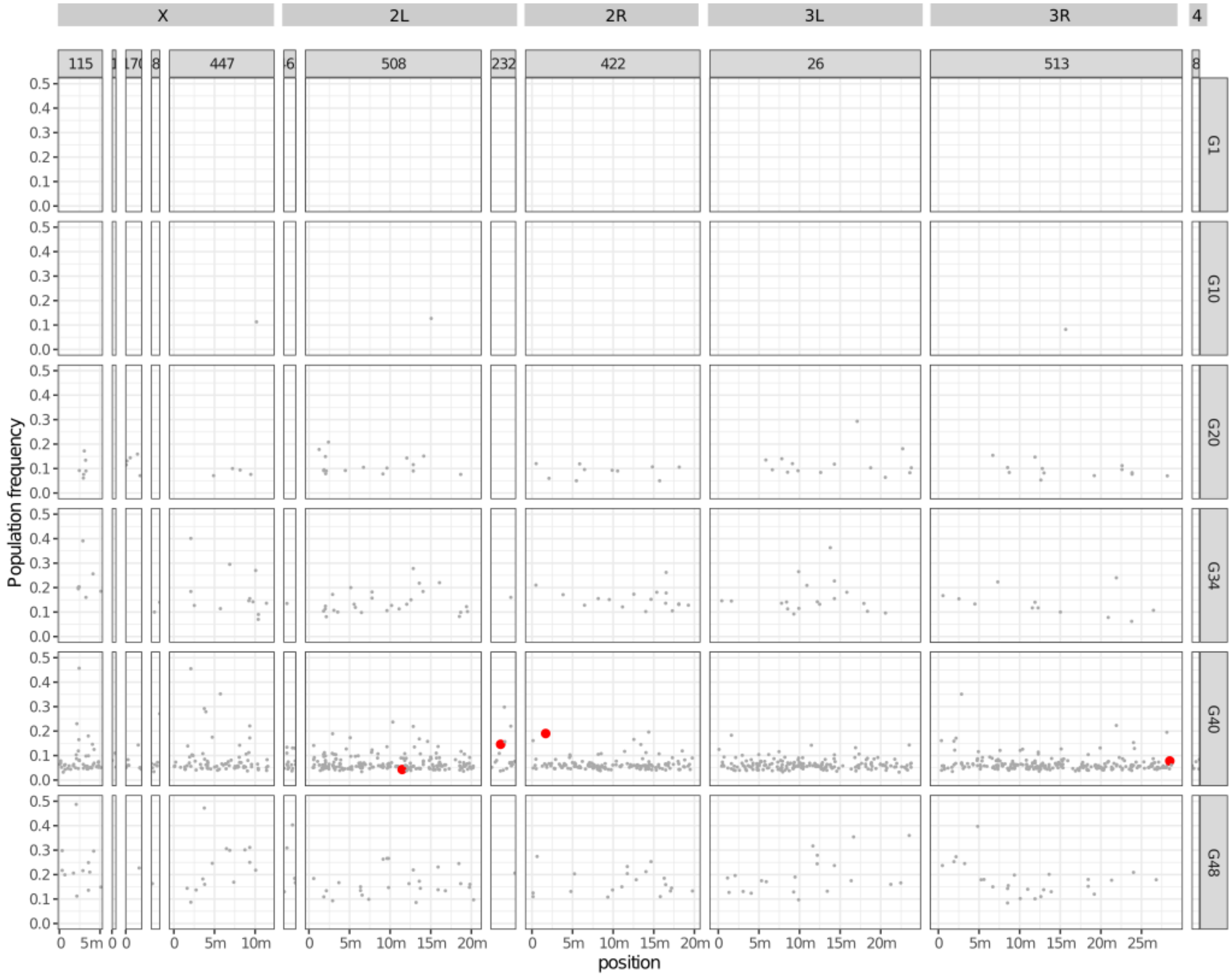


Figure 17: Position and population frequency of P-element insertions in replicate 1. The contigs are in the top panel and the generations in the right panels. P-element insertions in piRNA clusters are shown in red. The likely order of *D. erecta* contigs was inferred by aligning the contigs to the *D. melanogaster* genome using Mauve [Darling et al., 2004]. m million base pairs

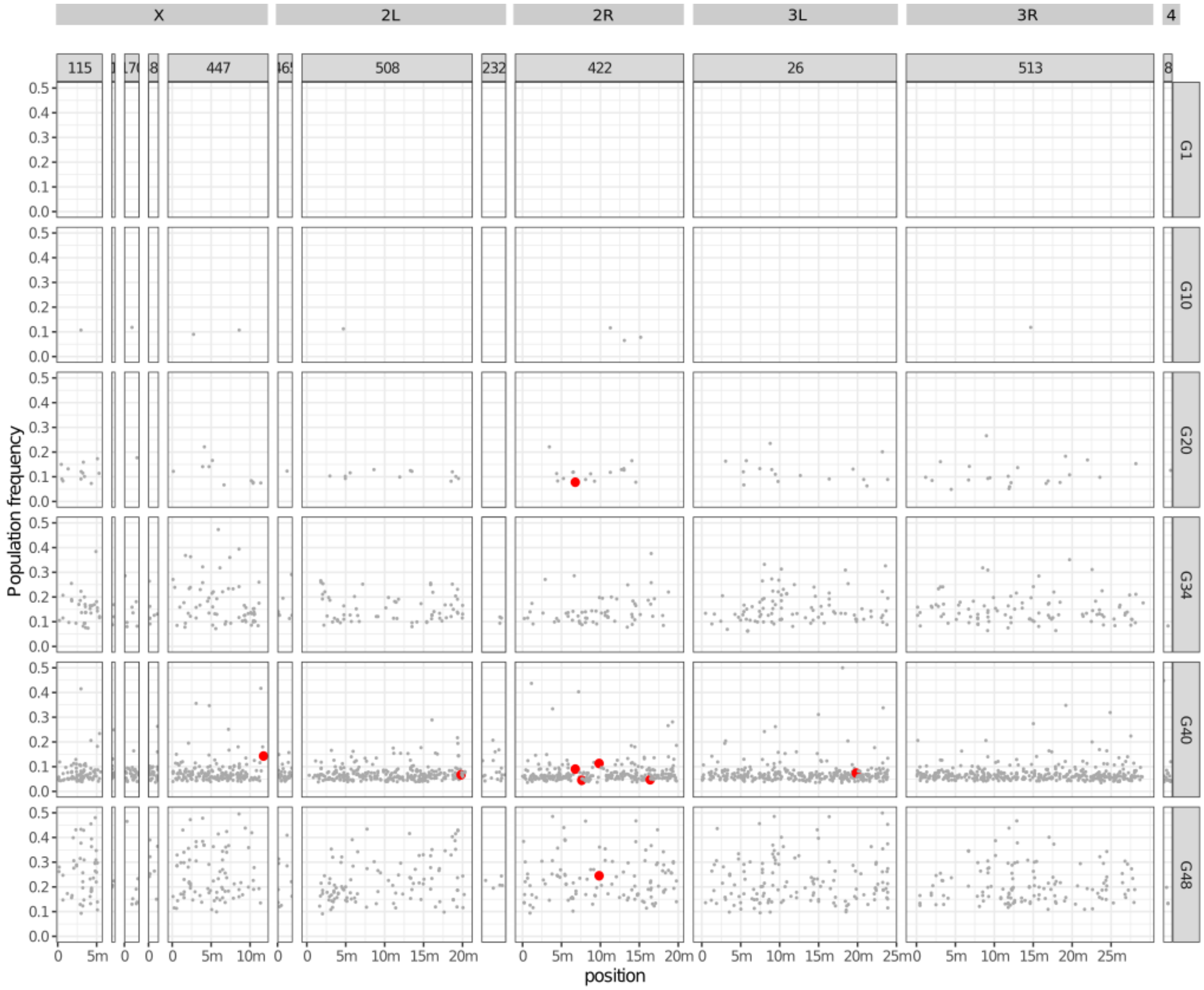


Figure 18: Position and population frequency of P-element insertions in replicate 2.

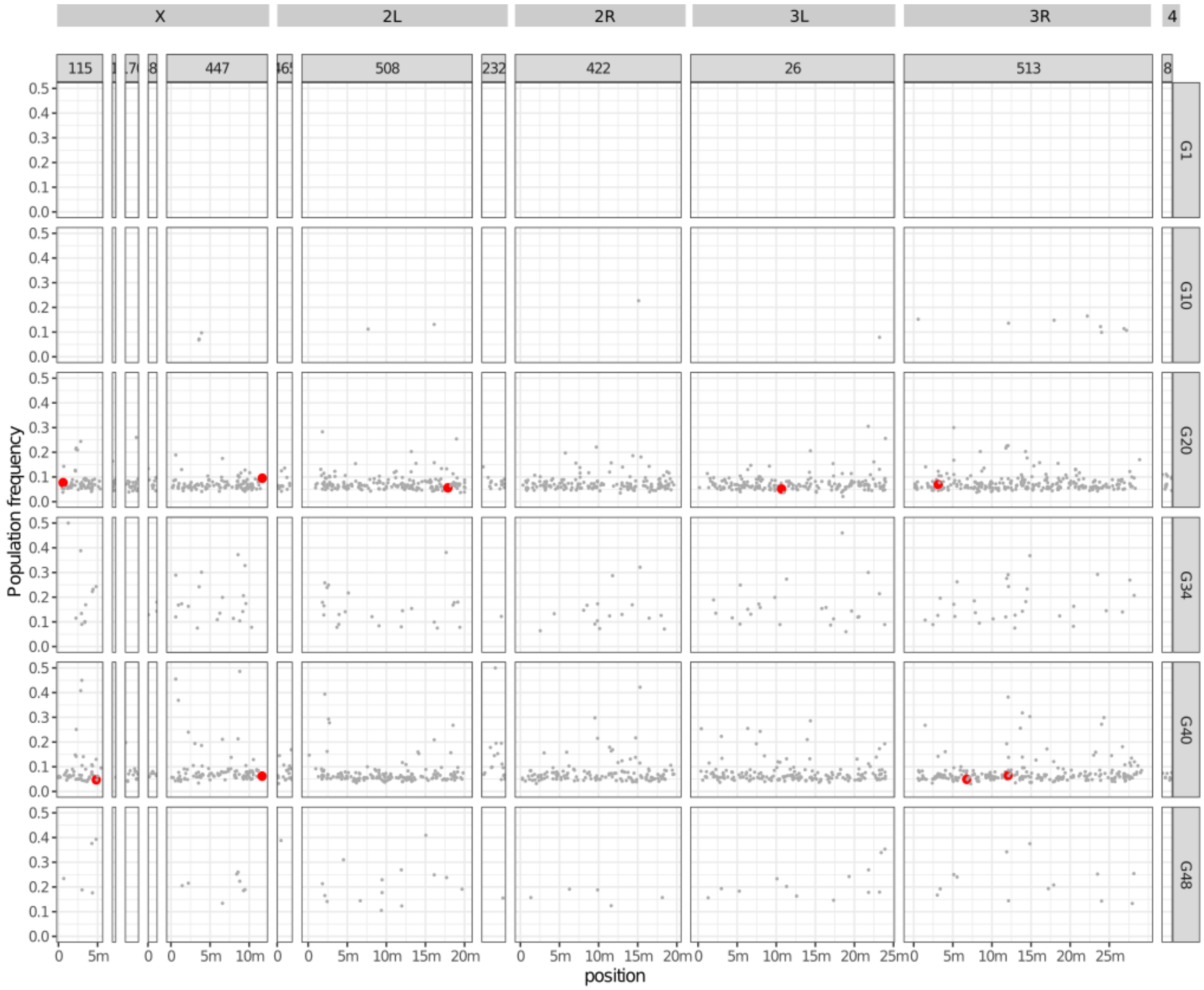


Figure 19: Position and population frequency of P-element insertions in replicate 4.

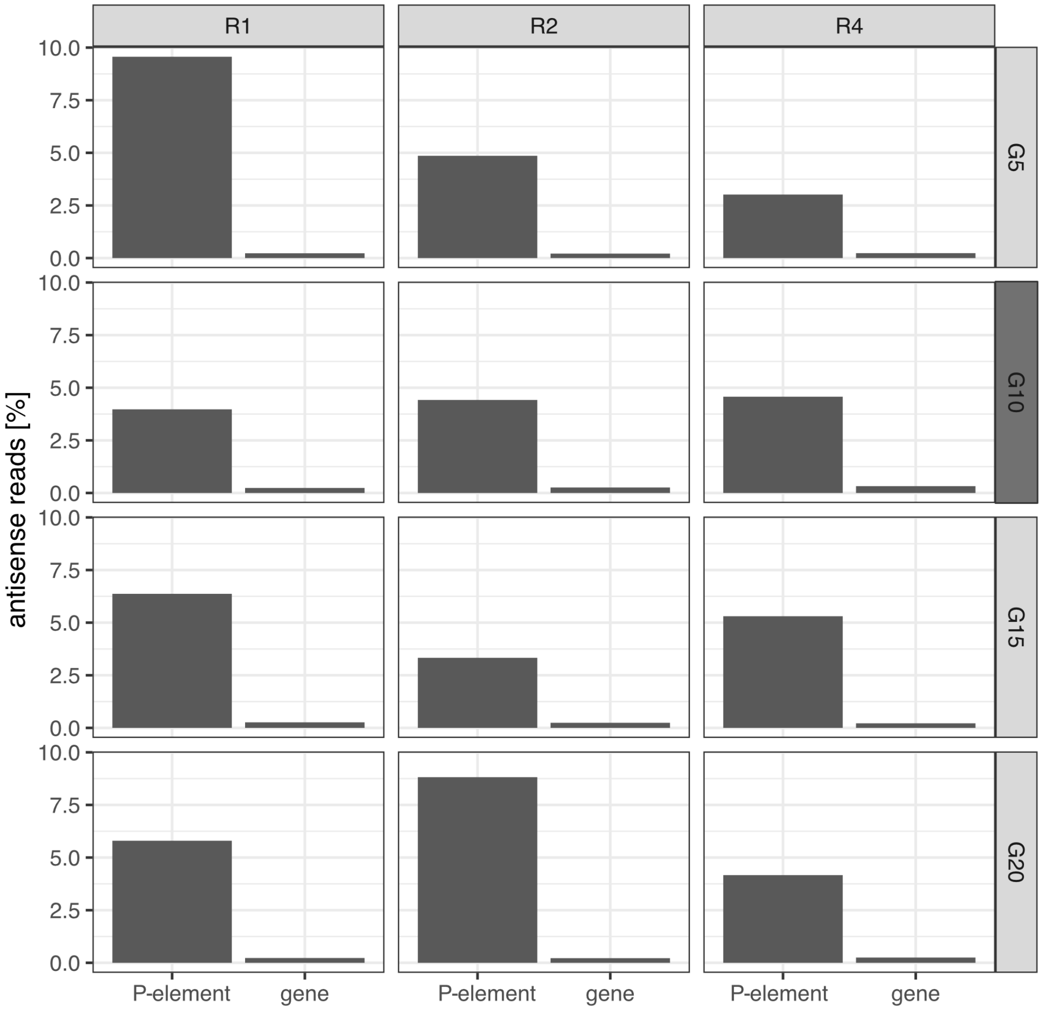


Figure 20: Percentage of antisense reads for the P-element and *D. erecta* transcripts at early generations of the invasion ($\leq$ 20 generations). Stranded RNA-seq data were generated for female flies (light grey panels) and ovaries (dark grey panel).

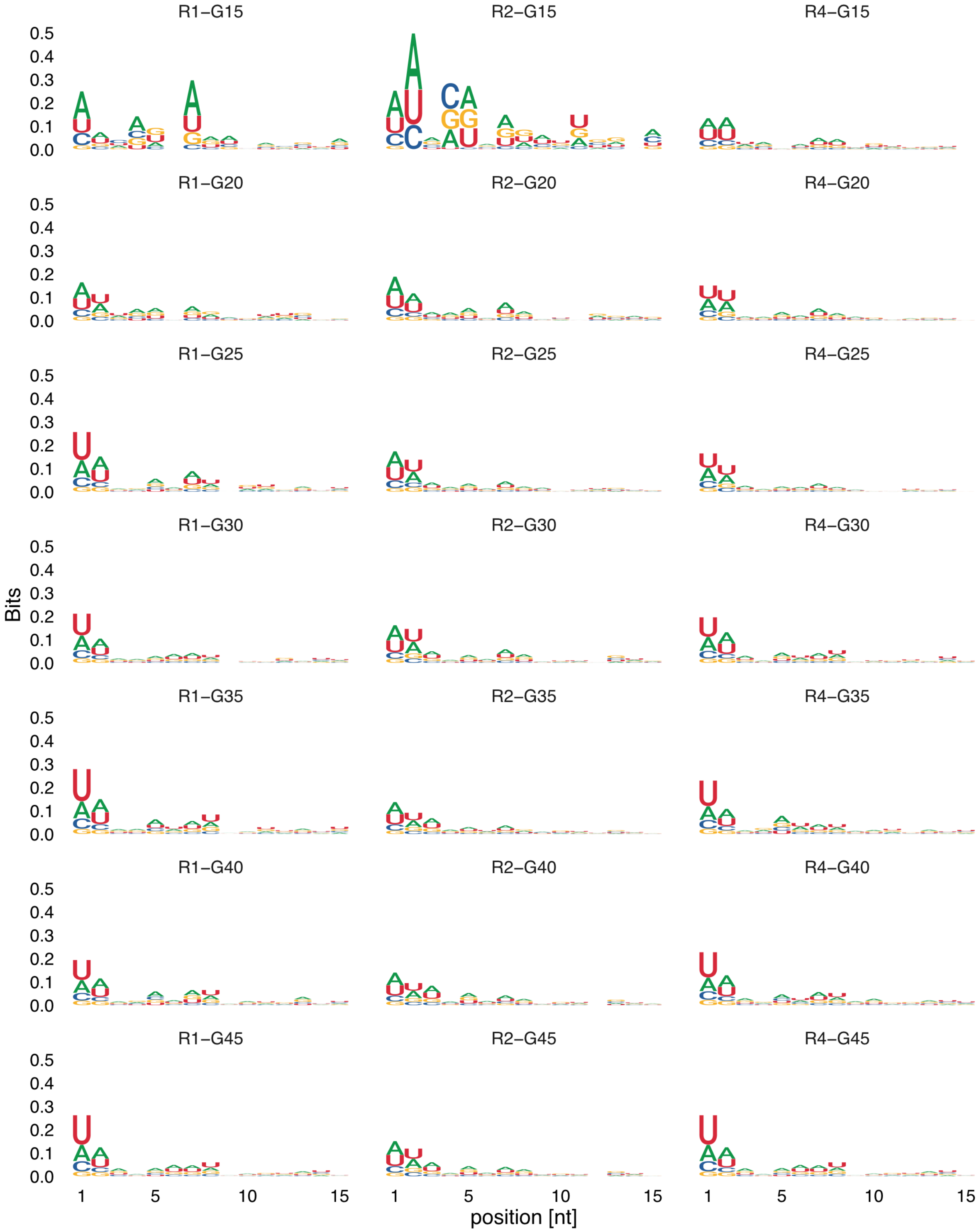


Figure 21: DNA motif of P-element siRNAs (20-22nt). Data are shown for three replicates (R1, R2, R4) and multiple generations during the invasion (G15-G45). Prior to generation 15 the abundance of siRNAs was too small for computing motifs.

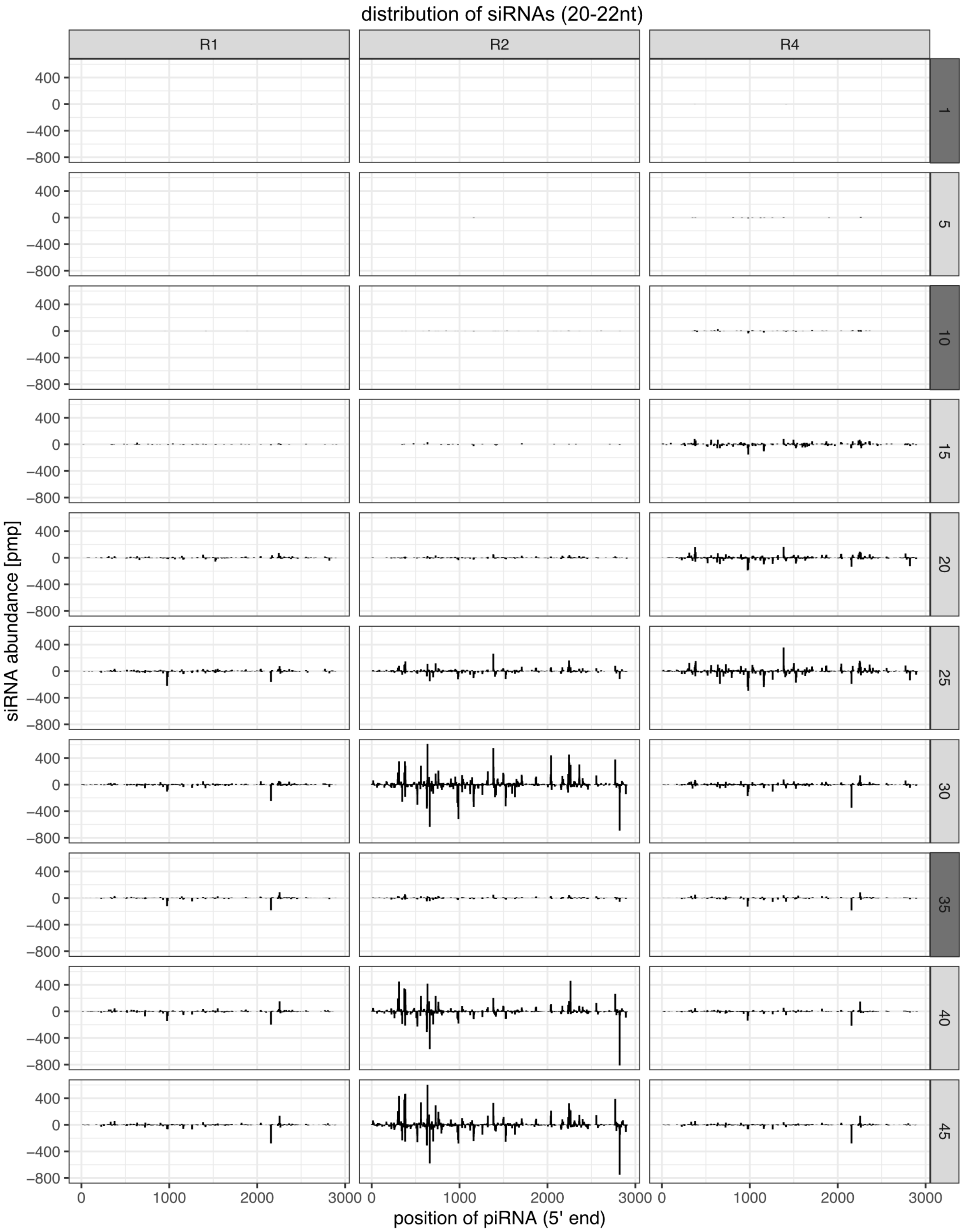


Figure 22: Distribution of siRNAs (20-22nt) along the P-element. Only the 5' positions of siRNAs are shown. The abundance of siRNAs is normalized to one million piRNAs (pmp). Replicates are at the top panel and the generations at the right panel. Sense piRNAs are shown on the positive y-axis and antisense piRNAs on the negative y-axis. We extracted small RNAs either from whole bodies of female flies (light grey panels) or from ovaries (dark grey panels).

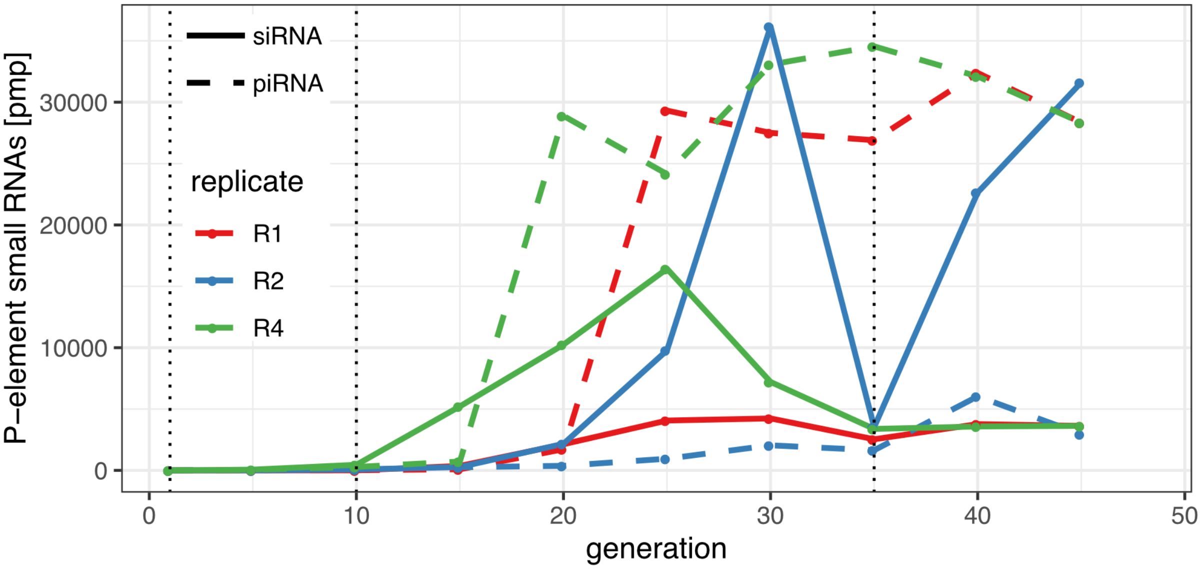


Figure 23: Abundance of siRNAs (20-22nt) during the P-element invasion in the three replicates (solid line). The abundance of piRNAs (23-29nt) is shown as reference (dashed lines). RNA was extracted from whole bodies of female flies except for time points marked by vertical dotted lines where RNA was extracted from ovaries.

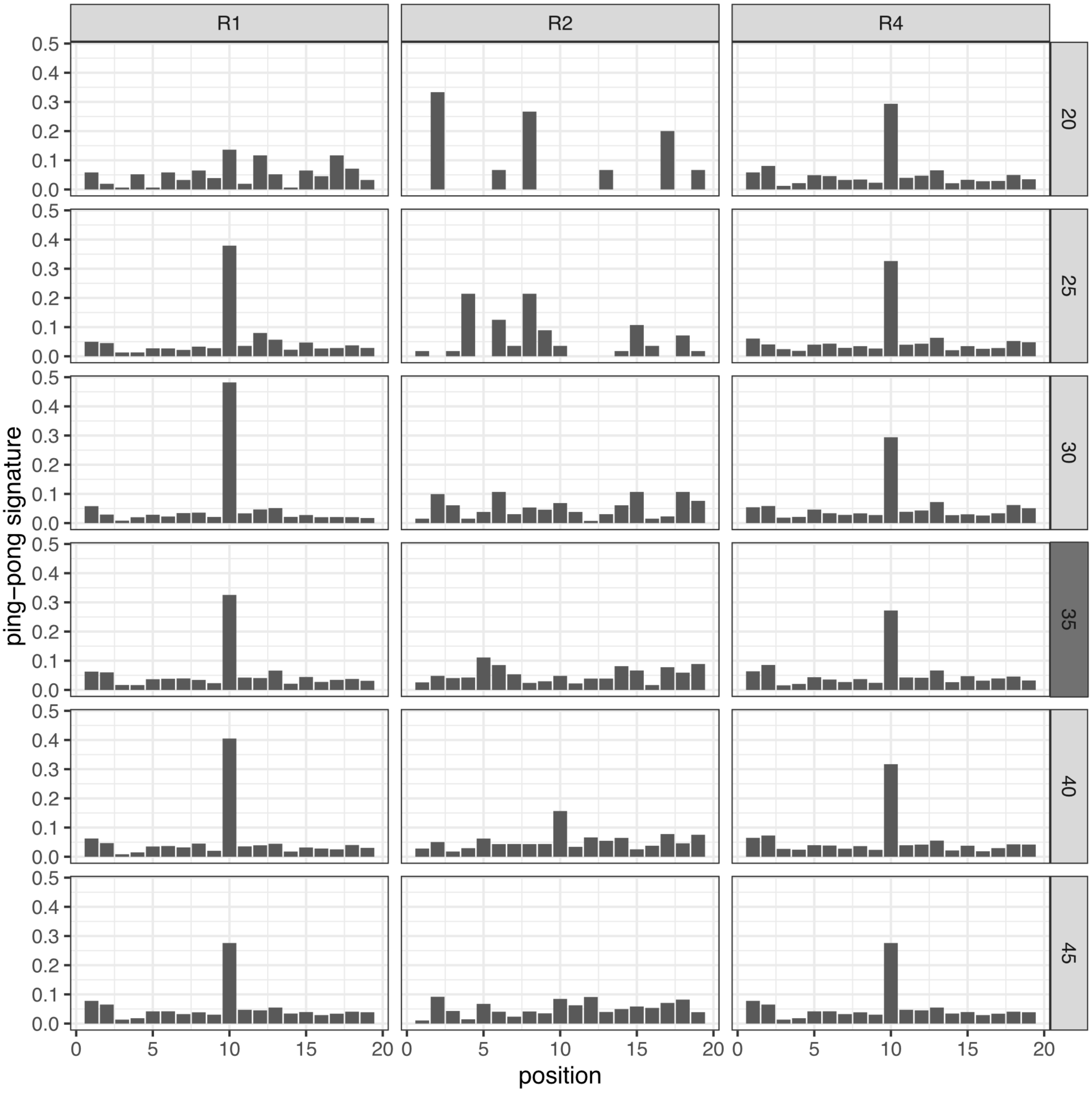


Figure 24: Ping-pong signatures for the P-element during the invasion. Data are shown for different replicates (top panel) and generations during the invasion (right panel). small RNAs were either extracted from whole bodies of female flies (light grey panels) or ovaries (dark grey panels). Due to a low number of piRNAs we could not compute ping-pong signatures for any replicate before generation 20.

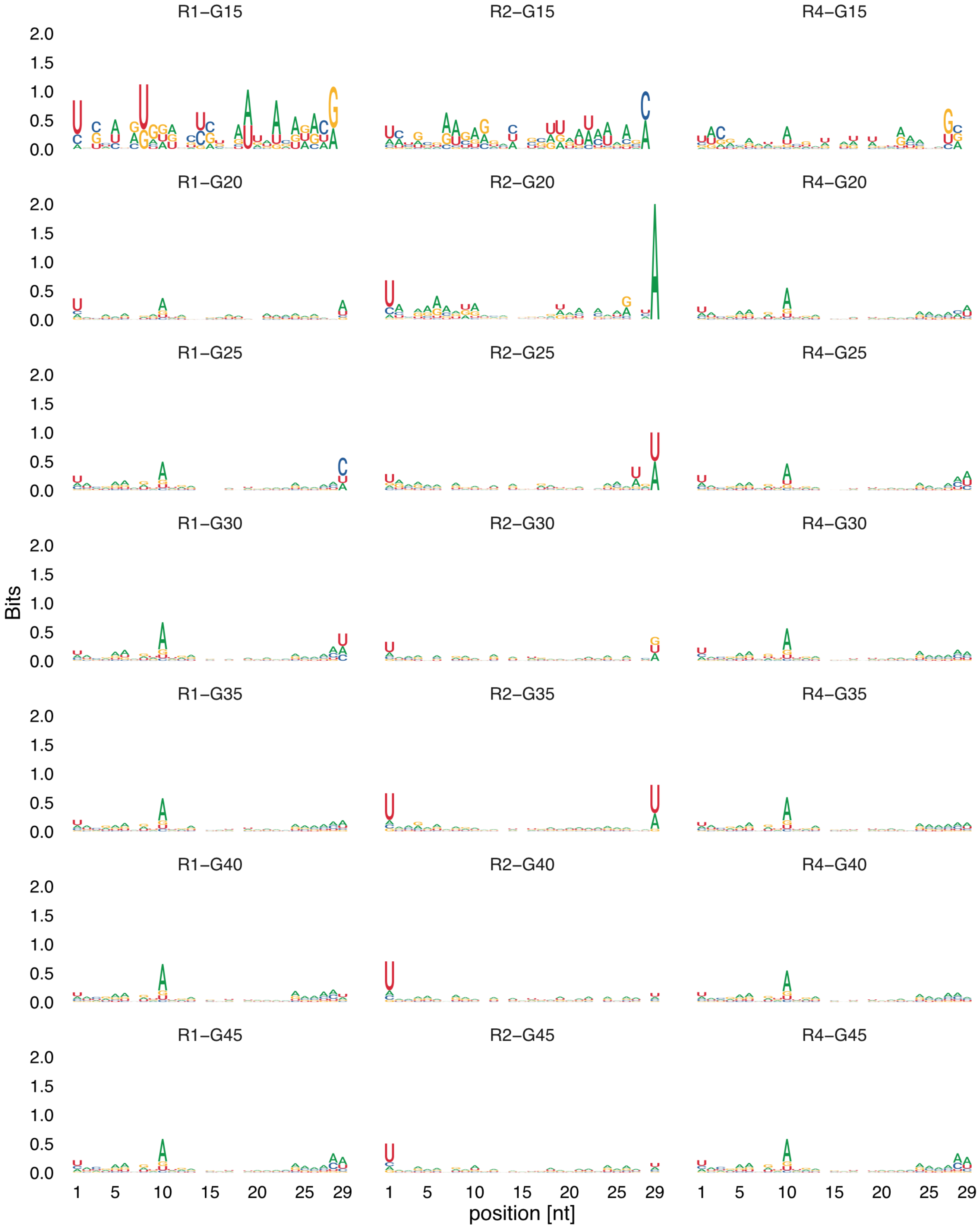


Figure 25: DNA motif of sense piRNAs of the P-element (23-29nt). Data are shown for three replicates (R1, R2, R4) and multiple generations during the invasion (G15-G45).

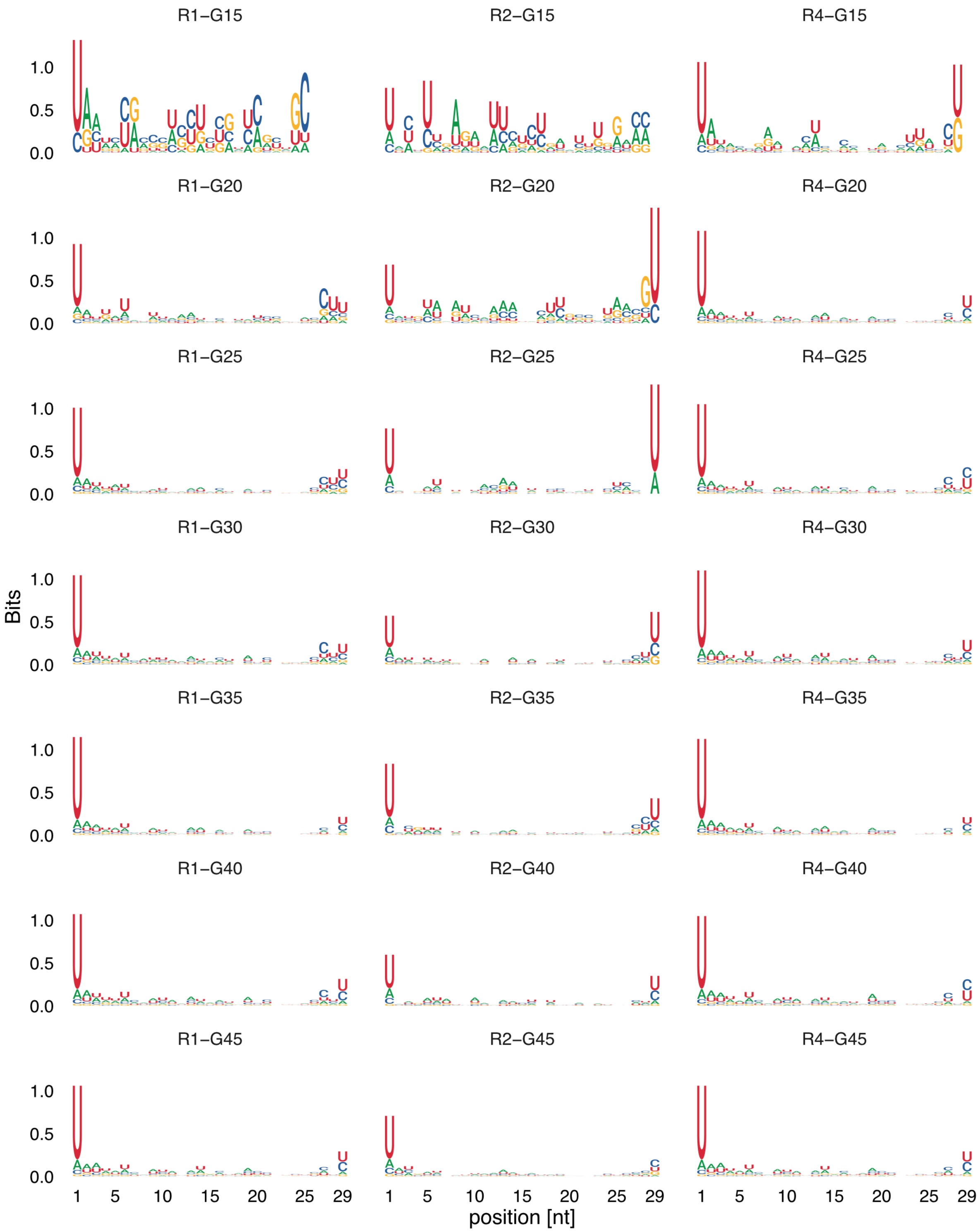


Figure 26: DNA motif of antisense piRNAs of the P-element (23-29nt). Data are shown for three replicates (R1, R2, R4) and multiple generations during the invasion (G15-G45).

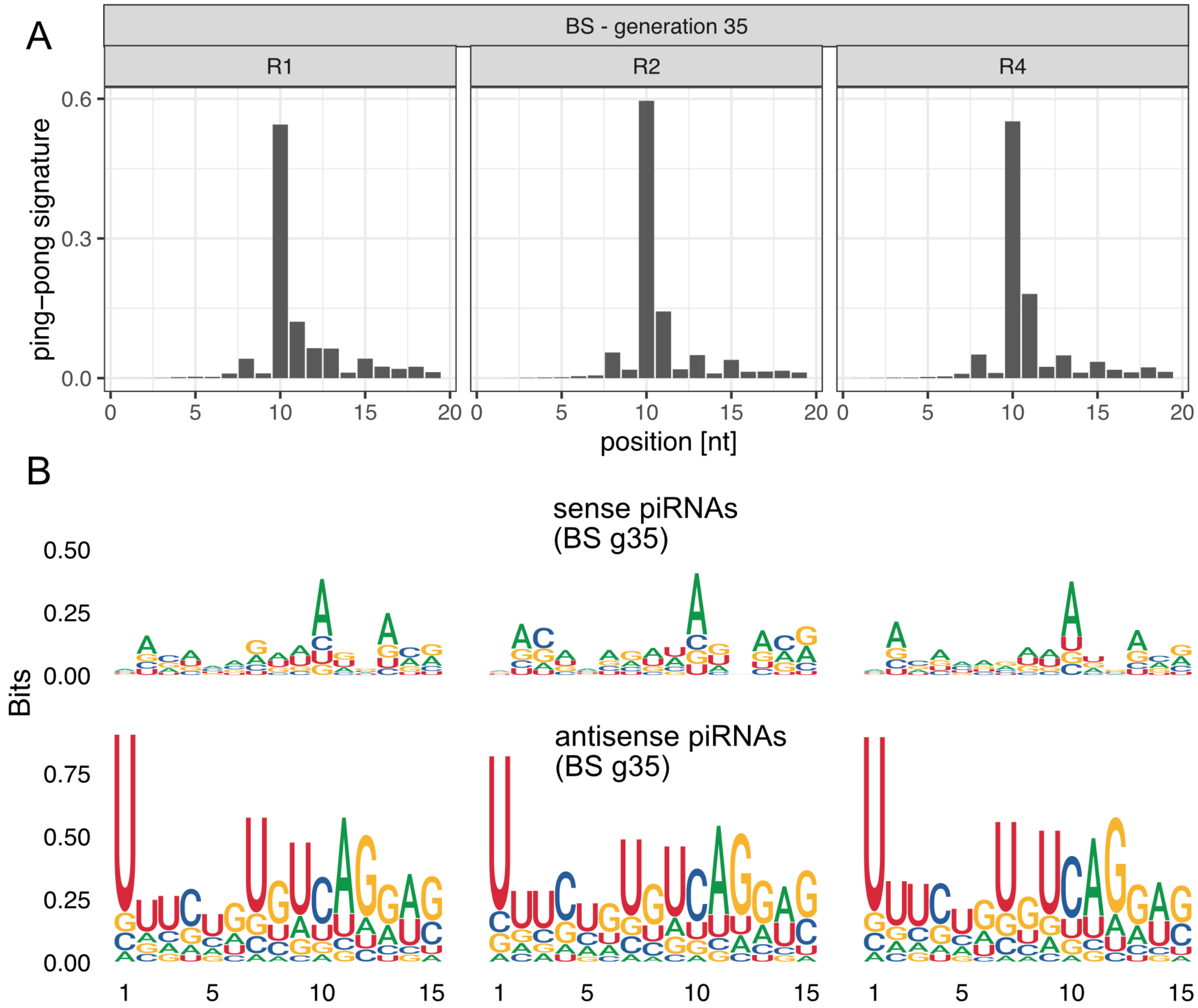


Figure 27: A) The ping-pong signature of BS in ovaries at generation 35. B) Motifs of sense and antisense piRNAs (23-29nt) complementary to BS in ovaries at generation 35.

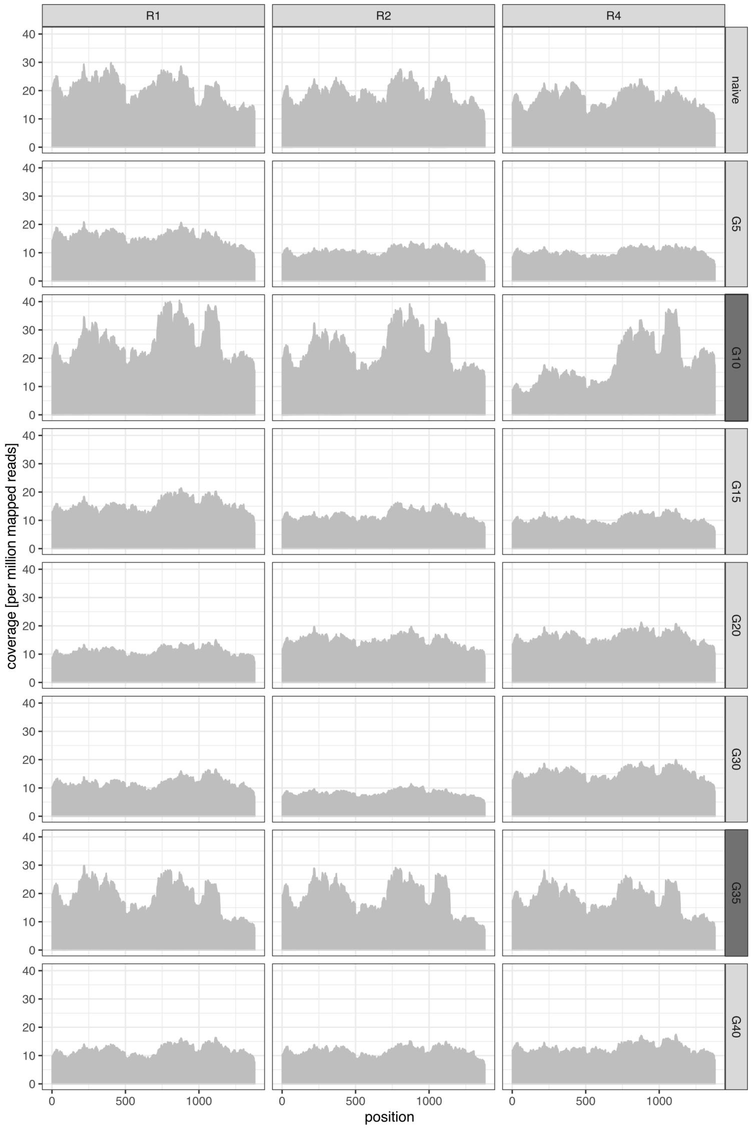


Figure 28: Expression level of FBtr0141271 (the *D. erecta* ortholog of Chk2) in our experimental populations. Results are shown for all replicates (top panel) at different generations (right panel). RNA extracted from whole flies (light grey panel) or ovaries (dark grey panel) was aligned to *D. erecta* transcripts and the coverage was normalized to a million mapped reads. No significant differences in the average coverage of FBtr0141271 were found between replicate 2 and replicates 1,4 (Wilcoxon rank sum test $W = 59$, $p = 0.79$).

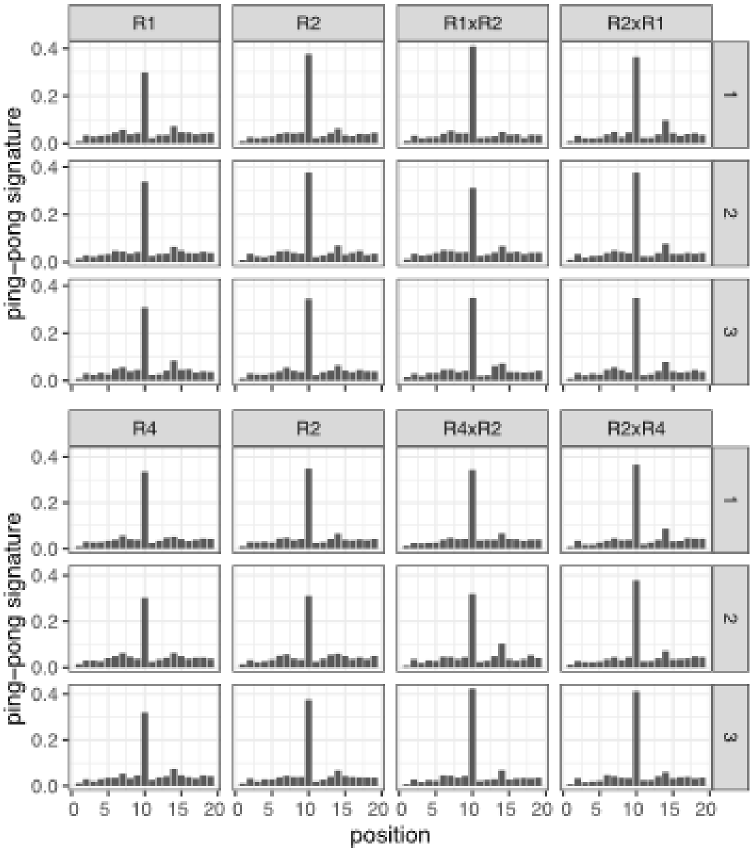


Figure 29: Ping-pong signatures of Quasimodo in the parents (R1, R2, R4) and the offspring (R1xR2, R2xR1, R4xR2, R2xR4) of reciprocal crosses among replicates.

### Supplementary tables

Table 1: Overview of the Pool-Seq data used in this work. For all replicates (rep) and generations (gen) we show the lane ID, the sequencing technology (platform), the read length (rl.), the inner distance (id.) and the number of sequenced paired-end reads in millions. Samples from naive flies not having the P-element are also included.

| rep | gen | lane ID | platform | rl. | id. | reads [M] |
|---|---|---|---|---|---|---|
| naive | - | CEBYBANXX | Illumina HiSeq 2500 | 125 | xx | 13.2 |
| naive | - | CD67CANXX | Illumina HiSeq 2500 | 125 | xx | 19.0 |
| R1 | G1 | CDMKUANXX | Illumina HiSeq 2500 | 125 | 55 | 27.6 |
| R1 | G10 | CDMKUANXX | Illumina HiSeq 2500 | 125 | 52 | 25.9 |
| R1 | G20 | CDMKUANXX | Illumina HiSeq 2500 | 125 | 52 | 28.8 |
| R1 | G34 | CE7UPANXX | Illumina HiSeq 2500 | 125 | 45 | 22.2 |
| R1 | G40 | CEBYBANXX | Illumina HiSeq 2500 | 125 | 60 | 30.1 |
| R1 | G48 | CD67CANXX | Illumina HiSeq 2500 | 125 | 43 | 23.7 |
| R2 | G1 | CDMKUANXX | Illumina HiSeq 2500 | 125 | 44 | 31.7 |
| R2 | G10 | CDMKUANXX | Illumina HiSeq 2500 | 125 | 50 | 25.9 |
| R2 | G20 | CDMKUANXX | Illumina HiSeq 2500 | 125 | 56 | 26.4 |
| R2 | G34 | CE7UPANXX | Illumina HiSeq 2500 | 125 | 39 | 30.8 |
| R2 | G40 | CEBYBANXX | Illumina HiSeq 2500 | 125 | 72 | 25.6 |
| R2 | G48 | CD67CANXX | Illumina HiSeq 2500 | 125 | 34 | 26.9 |
| R4 | G1 | CDMKUANXX | Illumina HiSeq 2500 | 125 | 44 | 27.1 |
| R4 | G10 | CDMKUANXX | Illumina HiSeq 2500 | 125 | 55 | 24.5 |
| R4 | G20 | CDMKUANXX | Illumina HiSeq 2500 | 125 | 57 | 25.8 |
| R4 | G34 | CE7UPANXX | Illumina HiSeq 2500 | 125 | 51 | 23.0 |
| R4 | G40 | CEBYBANXX | Illumina HiSeq 2500 | 125 | 79 | 22.9 |
| R4 | G48 | CD67CANXX | Illumina HiSeq 2500 | 125 | 50 | 18.8 |

Table 2: Abundance and effective transposition rate ($u'$) of the P-element in the experimentally evolving *D. erecta* populations. Data are shown for different replicates (rep) and generations (gen). The abundance of the P-element was estimated as reads per million (rpm) using PoPoolationTE2 [Kofler et al., 2016] and as insertions per haploid genome (ins) using DeviaTE [Weilguny and Kofler, 2019]. t.[M]: total number of reads in million; P: number of reads mapping to the P-element.

| rep | gen | t.[M] | P | rpm | ins | $u'_{rpm}$ | $u'_{ins}$ |
|---|---|---|---|---|---|---|---|
| R1 | 1 | 27.6 | 442 | 8.0 | 0.4 | - | - |
| R1 | 10 | 25.8 | 2,439 | 47.2 | 2.4 | 0.218 | 0.224 |
| R1 | 20 | 28.8 | 18,469 | 320.3 | 15.5 | 0.211 | 0.203 |
| R1 | 34 | 22.1 | 20,510 | 462.5 | 21.8 | 0.027 | 0.025 |
| R1 | 40 | 30 | 29,032 | 482.8 | 22.3 | 0.007 | 0.004 |
| R1 | 48 | 23.6 | 25,205 | 531.9 | 26.9 | 0.012 | 0.023 |
| R2 | 1 | 31.6 | 686 | 10.8 | 0.5 | - | - |
| R2 | 10 | 25.8 | 2,292 | 44.3 | 2.0 | 0.169 | 0.156 |
| R2 | 20 | 26.3 | 17,477 | 331.6 | 15.1 | 0.223 | 0.224 |
| R2 | 34 | 30.7 | 98,445 | 1,599.2 | 92.5 | 0.119 | 0.138 |
| R2 | 40 | 25.5 | 100,053 | 1,954.9 | 125.8 | 0.034 | 0.053 |
| R2 | 48 | 26.8 | 136,585 | 2,543.3 | 151.4 | 0.033 | 0.023 |
| R4 | 1 | 27.1 | 567 | 10.4 | 0.5 | - | - |
| R4 | 10 | 24.4 | 7,340 | 149.8 | 7.1 | 0.344 | 0.342 |
| R4 | 20 | 25.8 | 28,330 | 548.1 | 25.9 | 0.138 | 0.139 |
| R4 | 34 | 22.9 | 28,460 | 619.7 | 31.7 | 0.009 | 0.014 |
| R4 | 40 | 22.8 | 29,645 | 648.3 | 30.1 | 0.008 | -0.008 |
| R4 | 48 | 18.8 | 25,880 | 687.3 | 37.0 | 0.007 | 0.026 |

Table 3: Overview of small RNA data used in this work. For each replicate (r.) we sequenced small RNA at multiple time points (gen.) during the P-element invasions and assessed the abundance of reads mapping to TEs (i.e. siRNAs and piRNAs), miRNAs, tRNAs, rRNAs and mRNAs. For each class of small RNAs we also estimated the fraction of reads mapping to the sense strand. Small RNAs were sequenced either from ovaries (tis.: ov) embrios (tis.: em) or whole bodies (tis.: b). Additionally, we sequenced whole bodies and ovaries of female flies not having the P-element using three replicates (marked with '-' at r. and gen.). t.[M]: total number of reads in million, m.[M]: mapped reads in million, sr: sub-replicate

| | | | | | abundance [%] | | | | | fraction sense[%] | | | | |
|---|---|---|---|---|---|---|---|---|---|---|---|---|---|---|
| r. | gen. | tis. | t. [M] | m.[M] | TE | miRNA | tRNA | rRNA | mRNA | TE | miRNA | tRNA | rRNA | mRNA |
| R1 | 1 | ov | 17.3 | 8.8 | 15.5 | 36.1 | 1.7 | 36.9 | 9.9 | 6.5 | 100.0 | 99.8 | 99.4 | 53.9 |
| R1 | 5 | b | 11.4 | 4.9 | 4.3 | 55.0 | 6.2 | 28.1 | 6.5 | 7.7 | 100.0 | 99.9 | 99.8 | 74.7 |
| R1 | 10 | ov | 18.5 | 8.1 | 16.5 | 28.2 | 6.7 | 37.4 | 11.2 | 6.8 | 100.0 | 99.9 | 99.5 | 54.8 |
| R1 | 15 | b | 10.9 | 4.7 | 5.2 | 67.6 | 5.1 | 17.3 | 4.8 | 8.0 | 100.0 | 99.9 | 99.6 | 59.0 |
| R1 | 20 | b | 11.5 | 4.7 | 4.8 | 76.2 | 3.8 | 12.0 | 3.3 | 10.4 | 100.0 | 99.9 | 99.4 | 53.9 |
| R1 | 25 | b | 9.2 | 3.9 | 5.1 | 72.5 | 2.8 | 15.9 | 3.7 | 9.5 | 100.0 | 99.8 | 99.5 | 56.6 |
| R1 | 30 | b | 15.3 | 4.4 | 5.3 | 66.8 | 4.1 | 19.7 | 4.2 | 9.0 | 100.0 | 99.9 | 99.6 | 60.2 |
| R1 | 35 | ov | 13.1 | 6.0 | 19.7 | 53.0 | 2.1 | 15.3 | 9.9 | 8.5 | 100.0 | 99.7 | 97.9 | 44.0 |
| R1 | 40 | b | 11.3 | 5.0 | 6.8 | 79.1 | 2.9 | 6.7 | 4.5 | 9.2 | 100.0 | 99.8 | 98.2 | 54.0 |
| R1 | 45 | b | 11.5 | 4.8 | 7.2 | 76.4 | 3.3 | 8.6 | 4.5 | 8.0 | 100.0 | 99.9 | 98.6 | 51.6 |
| R2 | 1 | ov | 13.1 | 6.3 | 19.0 | 46.1 | 1.8 | 22.3 | 10.8 | 6.6 | 100.0 | 99.7 | 98.9 | 46.7 |
| R2 | 5 | b | 12.7 | 5.2 | 5.3 | 67.8 | 5.5 | 15.2 | 6.2 | 8.0 | 100.0 | 99.9 | 99.5 | 69.3 |
| R2 | 10 | ov | 10.7 | 5.5 | 9.5 | 23.2 | 3.5 | 55.8 | 8.0 | 7.1 | 100.0 | 99.9 | 99.8 | 59.6 |
| R2 | 15 | b | 9.9 | 3.5 | 3.7 | 71.3 | 4.1 | 16.7 | 4.3 | 9.5 | 100.0 | 99.9 | 99.7 | 66.5 |
| R2 | 20 | b | 12.8 | 5.5 | 4.4 | 74.8 | 4.1 | 12.9 | 3.8 | 10.9 | 100.0 | 99.9 | 99.6 | 61.3 |
| R2 | 25 | b | 12.1 | 5.2 | 4.2 | 73.2 | 4.2 | 14.4 | 4.0 | 10.7 | 100.0 | 99.9 | 99.6 | 64.2 |
| R2 | 30 | b | 13.0 | 6.1 | 3.2 | 67.2 | 7.4 | 18.2 | 4.1 | 11.8 | 100.0 | 99.9 | 99.8 | 73.9 |
| R2 | 35 | ov | 5.4 | 2.5 | 18.9 | 57.1 | 1.5 | 12.4 | 10.1 | 9.2 | 100.0 | 99.7 | 97.5 | 43.3 |
| R2 | 40 | b | 12.8 | 5.7 | 5.4 | 76.1 | 4.9 | 9.2 | 4.4 | 10.1 | 100.0 | 99.9 | 99.1 | 60.0 |
| R2 | 45 | b | 15.3 | 7.1 | 5.6 | 76.6 | 4.1 | 9.5 | 4.3 | 9.7 | 100.0 | 99.9 | 98.9 | 57.2 |
| R4 | 1 | ov | 14.3 | 7.3 | 13.8 | 26.7 | 1.5 | 47.9 | 10.1 | 6.8 | 100.0 | 99.8 | 99.6 | 57.7 |
| R4 | 5 | b | 13.6 | 5.6 | 4.8 | 67.1 | 6.0 | 16.9 | 5.2 | 7.8 | 100.0 | 99.9 | 99.6 | 66.8 |
| R4 | 10 | ov | 13.9 | 8.4 | 2.7 | 10.0 | 1.6 | 82.2 | 3.5 | 10.6 | 100.0 | 99.9 | 100.0 | 75.5 |
| R4 | 15 | b | 10.2 | 4.6 | 2.0 | 58.9 | 4.2 | 30.5 | 4.5 | 11.9 | 100.0 | 99.9 | 99.9 | 79.4 |
| R4 | 20 | b | 10.7 | 4.7 | 7.3 | 75.0 | 4.5 | 8.4 | 4.8 | 8.2 | 100.0 | 99.9 | 98.7 | 54.8 |
| R4 | 25 | b | 10.5 | 4.4 | 5.2 | 72.9 | 3.3 | 14.8 | 3.8 | 10.0 | 100.0 | 99.8 | 99.4 | 54.7 |
| R4 | 30 | b | 13.2 | 6.0 | 7.0 | 77.6 | 3.3 | 7.9 | 4.2 | 8.5 | 100.0 | 99.9 | 98.5 | 50.5 |
| R4 | 35 | ov | 17.8 | 8.0 | 20.8 | 57.4 | 1.3 | 10.6 | 10.1 | 9.4 | 100.0 | 99.5 | 97.4 | 42.3 |
| R4 | 40 | b | 12.3 | 6.0 | 6.4 | 79.5 | 3.4 | 6.3 | 4.4 | 9.7 | 100.0 | 99.9 | 98.7 | 54.9 |
| R4 | 45 | b | 14.1 | 6.6 | 6.8 | 78.1 | 3.6 | 6.8 | 4.7 | 8.2 | 100.0 | 99.9 | 98.4 | 54.1 |
| - | - | b | 9.7 | 4.3 | 7.8 | 77.8 | 3.1 | 6.5 | 4.9 | 7.6 | 100.0 | 99.8 | 97.8 | 49.3 |
| - | - | b | 12.6 | 5.8 | 7.2 | 77.2 | 4.1 | 6.8 | 4.6 | 7.9 | 100.0 | 99.9 | 98.1 | 50.3 |
| - | - | b | 30.6 | 15.1 | 6.4 | 69.6 | 2.9 | 16.7 | 4.5 | 8.0 | 100.0 | 99.8 | 99.4 | 54.2 |
| - | - | ov | 10.9 | 5.1 | 21.1 | 59.9 | 2.0 | 6.4 | 10.7 | 7.9 | 100.0 | 99.7 | 94.2 | 40.3 |
| - | - | ov | 10.2 | 4.9 | 18.7 | 51.7 | 1.8 | 17.7 | 10.1 | 7.9 | 100.0 | 99.7 | 97.9 | 45.2 |
| - | - | ov | 9.4 | 4.2 | 22.7 | 57.7 | 2.3 | 6.1 | 11.2 | 7.7 | 100.0 | 99.7 | 93.9 | 40.9 |
| | - | em | 4.9 | 2.9 | 10.9 | 67.7 | 1.3 | 13.3 | 6.9 | 4.0 | 100.0 | 99.1 | 99.3 | 45.2 |
| R1_sr1 | G70 | b | 9.9 | 4.9 | 8.6 | 76.6 | 5.5 | 5.8 | 3.6 | 8.6 | 100.0 | 100.0 | 99.4 | 47.3 |
| R1_sr2 | G70 | b | 9.6 | 4.7 | 8.4 | 68.2 | 5.6 | 13.8 | 4.0 | 8.9 | 100.0 | 100.0 | 99.8 | 51.6 |
| R1_sr3 | G70 | b | 12.8 | 6.1 | 9.2 | 74.3 | 5.5 | 6.7 | 4.4 | 9.0 | 100.0 | 99.9 | 99.4 | 49.5 |
| R1xR2_sr1 | F1 | b | 9.3 | 4.2 | 7.8 | 73.3 | 6.5 | 8.8 | 3.6 | 10.18 | 100.0 | 100.0 | 99.6 | 53.2 |
| R1xR2_sr2 | F1 | b | 9.1 | 4.3 | 8.7 | 74.6 | 6.5 | 6.8 | 3.5 | 9.6 | 100.0 | 100.0 | 99.6 | 46.5 |
| R1xR2_sr3 | F1 | b | 8.6 | 4.2 | 6.8 | 74.8 | 7.6 | 7.6 | 3.3 | 10.76 | 100.0 | 100.0 | 99.6 | 53.6 |
| R2_sr1 | G67 | b | 10.8 | 5.2 | 7.2 | 75.9 | 5.6 | 8.0 | 3.4 | 10.9 | 100.0 | 100.0 | 99.6 | 48.4 |
| R2_sr2 | G67 | b | 9.4 | 4.5 | 5.9 | 71.7 | 6.7 | 12.5 | 3.2 | 12.3 | 100.0 | 100.0 | 99.8 | 54.2 |
| R2_sr3 | G67 | b | 8.2 | 4.0 | 6.3 | 74.6 | 7.2 | 8.5 | 3.4 | 10.4 | 100.0 | 100.0 | 99.6 | 53.5 |
| R2_sr4 | G67 | b | 7.4 | 3.6 | 6.4 | 70.5 | 9.3 | 10.0 | 3.8 | 9.7 | 100.0 | 100.0 | 99.7 | 57.8 |
| R2_sr5 | G67 | b | 10.0 | 5.1 | 6.5 | 74.9 | 7.1 | 8.1 | 3.5 | 11.9 | 100.0 | 100.0 | 99.7 | 55.4 |
| R2_sr6 | G67 | b | 11.8 | 5.9 | 6.8 | 71.3 | 7.6 | 10.4 | 3.9 | 11.3 | 100.0 | 100.0 | 99.7 | 56.6 |
| R2xR1_sr1 | F1 | b | 11.4 | 6.4 | 3.6 | 43.9 | 17.0 | 30.0 | 5.5 | 11.3 | 100.0 | 100.0 | 99.9 | 83.4 |
| R2xR1_sr2 | F1 | b | 11.0 | 6.1 | 4.9 | 58.3 | 21.4 | 11.0 | 4.4 | 11.1 | 100.0 | 100.0 | 99.8 | 71.6 |
| R2xR1_sr3 | F1 | b | 11.3 | 6.3 | 4.3 | 52.2 | 27.1 | 11.6 | 4.8 | 11.4 | 100.0 | 100.0 | 99.8 | 77.2 |
| R2xR4_sr1 | F1 | b | 8.9 | 4.5 | 5.0 | 72.5 | 9.3 | 9.9 | 3.3 | 10.1 | 100.0 | 100.0 | 99.8 | 58.7 |
| R2xR4_sr2 | F1 | b | 9.8 | 4.9 | 5.7 | 71.1 | 9.7 | 9.8 | 3.7 | 10.7 | 100.0 | 100.0 | 99.8 | 60.7 |
| R2xR4_sr3 | F1 | b | 9.1 | 4.5 | 5.6 | 66.3 | 10.6 | 13.7 | 3.8 | 10.7 | 100.0 | 100.0 | 99.8 | 61.2 |
| R4_sr1 | G70 | b | 9.5 | 4.6 | 7.8 | 71.1 | 7.3 | 9.9 | 3.9 | 8.6 | 100.0 | 100.0 | 99.7 | 52.4 |
| R4_sr2 | G70 | b | 8.5 | 3.9 | 8.8 | 74.4 | 6.2 | 6.7 | 3.9 | 8.3 | 100.0 | 100.0 | 99.5 | 47.6 |
| R4_sr3 | G70 | b | 11.1 | 5.2 | 8.5 | 69.5 | 7.7 | 10.0 | 4.3 | 7.8 | 100.0 | 100.0 | 99.7 | 52.9 |
| R4xR2_sr1 | F1 | b | 6.4 | 2.7 | 7.7 | 72.0 | 9.0 | 7.7 | 3.7 | 11.1 | 100.0 | 100.0 | 99.7 | 50.8 |
| R4xR2_sr2 | F1 | b | 9.2 | 4.4 | 7.2 | 73.9 | 7.7 | 7.7 | 3.5 | 10.2 | 100.0 | 100.0 | 99.7 | 53.2 |
| R4xR2_sr3 | F1 | b | 8.0 | 4.1 | 6.4 | 63.4 | 10.6 | 15.8 | 3.8 | 10.2 | 100.0 | 100.0 | 99.9 | 63.4 |

Table 4: Overview of the RNA-Seq data used in this work. RNA was either extracted from ovaries (ov) or whole bodies of female flies (wf). Data are shown for different replicates (rep.) and generations (gen.) of the experimental populations. We also sequenced three replicates of naive flies not having the P-element. rl: read length, reads: total number of reads in million (including both reads of the paired ends), m.: mapped reads in million, hq.m.: reads mapped with a high mapping-quality in million ($\geq 20$). gene m.: reads mapping to a gene in million, $\%se_{gene}$: percentage of 'gene m.' reads aligning to sense strand, pele m.: reads mapping to the P-element, $\%se_{pele}$: percentage of 'pele m.' aligning to sense strand

| rep. | gen. | tissue | rl | reads | m. | hq.m. | gene m. | $\%se_{gene}$ | pele m. | $\%se_{pele}$ |
|---|---|---|---|---|---|---|---|---|---|---|
| - | naive | wf | 100 | 17.07 | 10.08 | 9.67 | 9.65 | 99.87 | 1 | - |
| - | naive | wf | 100 | 21.27 | 13.57 | 13.02 | 13.00 | 99.87 | 1 | - |
| - | naive | wf | 100 | 22.17 | 14.50 | 13.94 | 13.92 | 99.88 | 0 | - |
| R1 | 5 | wf | 100 | 58.72 | 39.60 | 37.91 | 37.84 | 99.86 | 82 | 91.11 |
| R1 | 10 | ov | 100 | 70.74 | 45.40 | 41.32 | 41.23 | 99.86 | 1763 | 96.97 |
| R1 | 15 | wf | 100 | 61.88 | 44.74 | 42.81 | 42.72 | 99.83 | 1672 | 93.93 |
| R1 | 20 | wf | 100 | 72.57 | 51.42 | 49.41 | 49.31 | 99.85 | 7062 | 94.69 |
| R1 | 30 | wf | 100 | 41.03 | 28.94 | 27.76 | 27.70 | 99.85 | 3327 | 87.12 |
| R1 | 35 | ov | 100 | 83.79 | 52.48 | 51.00 | 50.85 | 99.80 | 7531 | 83.94 |
| R1 | 40 | wf | 100 | 56.17 | 40.18 | 38.73 | 38.65 | 99.84 | 3849 | 88.52 |
| R2 | 5 | wf | 100 | 52.82 | 37.33 | 35.97 | 35.90 | 99.86 | 326 | 95.60 |
| R2 | 10 | ov | 100 | 78.33 | 50.41 | 46.72 | 46.61 | 99.83 | 291 | 97.00 |
| R2 | 15 | wf | 100 | 53.05 | 37.88 | 36.37 | 36.29 | 99.83 | 2537 | 96.91 |
| R2 | 20 | wf | 100 | 51.24 | 34.98 | 33.70 | 33.62 | 99.86 | 8207 | 91.17 |
| R2 | 30 | wf | 100 | 71.36 | 51.35 | 49.02 | 48.89 | 99.83 | 20539 | 92.02 |
| R2 | 35 | ov | 100 | 85.42 | 54.40 | 52.96 | 52.80 | 99.84 | 39917 | 93.72 |
| R2 | 40 | wf | 100 | 51.54 | 36.79 | 35.37 | 35.28 | 99.86 | 21325 | 93.65 |
| R4 | 5 | wf | 100 | 57.66 | 41.14 | 39.56 | 39.47 | 99.83 | 471 | 97.31 |
| R4 | 10 | ov | 100 | 73.43 | 43.94 | 40.86 | 40.74 | 99.78 | 2559 | 96.93 |
| R4 | 15 | wf | 100 | 58.40 | 42.43 | 40.60 | 40.52 | 99.85 | 6824 | 94.82 |
| R4 | 20 | wf | 100 | 73.02 | 49.39 | 47.52 | 47.41 | 99.85 | 6576 | 96.04 |
| R4 | 30 | wf | 100 | 46.35 | 32.79 | 31.49 | 31.43 | 99.87 | 3743 | 95.78 |
| R4 | 35 | ov | 100 | 79.75 | 52.30 | 51.00 | 50.89 | 99.86 | 6549 | 94.86 |
| R4 | 40 | wf | 100 | 60.70 | 43.15 | 41.74 | 41.65 | 99.86 | 5317 | 95.29 |

Table 5: Effect of piRNAs on the expression and splicing of the P-element. The expression level of the P-element (expr. in rpkm), the splicing level of the three introns (IVS1, IVS2, IVS3 in rpm) and the abundance of P-element piRNAs (ppm) are shown. Data are shown for all replicates (r.) at different generations (g.). Estimates of TE copy numbers (TEs) for generations 5, 15 and 30 are based on exponential interpolations (based on the transposition rate; see above) between the two closest neighbouring data points. For each independent variable (expr., IVS1, IVS2, IVS3) we generated a linear model where TEs and piRNAs served as explanatory variables. TE copy numbers were significantly positively correlated with all four independent variables ($p < 0.001$). piRNAs showed a significant negative correlation with the splicing level of IVS1 ($p = 0.0021$) and IVS3 ($p = 0.0015$) but did not show a significant correlation with the splicing level of IVS2 ($p = 0.34$) nor the expression of the P-element ($p = 0.067$).

| r. | g. | piRNAs | TEs | expr. | IVS1 | IVS2 | IVS3 |
|---|---|---|---|---|---|---|---|
| 1 | 5 | 14.8 | 0.89 | 0.83 | 0.18 | 0.08 | 0.00 |
| 1 | 15 | 90.7 | 6.15 | 12.80 | 1.25 | 1.45 | 0.20 |
| 1 | 20 | 1724.7 | 15.46 | 46.60 | 5.50 | 6.34 | 0.66 |
| 1 | 30 | 27500.0 | 19.79 | 39.20 | 4.39 | 4.53 | 0.03 |
| 1 | 40 | 32387.0 | 22.33 | 32.20 | 3.68 | 3.28 | 0.00 |
| 2 | 5 | 49.7 | 0.97 | 3.17 | 0.40 | 0.40 | 0.13 |
| 2 | 15 | 255.5 | 5.51 | 22.60 | 2.77 | 2.77 | 0.13 |
| 2 | 20 | 372.5 | 15.12 | 79.60 | 10.26 | 8.72 | 1.63 |
| 2 | 30 | 2041.8 | 55.08 | 139.00 | 21.77 | 12.87 | 1.27 |
| 2 | 40 | 6031.1 | 125.80 | 202.00 | 35.42 | 16.42 | 3.07 |
| 4 | 5 | 50.8 | 1.62 | 3.96 | 0.46 | 0.51 | 0.02 |
| 4 | 15 | 721.8 | 13.57 | 55.10 | 6.74 | 7.24 | 0.80 |
| 4 | 20 | 28886.0 | 25.91 | 44.60 | 3.93 | 5.37 | 0.20 |
| 4 | 30 | 33070.1 | 29.77 | 38.50 | 4.30 | 5.03 | 0.00 |
| 4 | 40 | 32101.9 | 30.13 | 41.30 | 5.08 | 5.03 | 0.00 |

Table 6: Extent of gonadal dysgenesis during the P-element invasion. Results are shown for all three replicates (rep) at different generations (gen). Naive flies not having the P-element (marked with '-' in rep and gen) were also analyzed. We estimated the number of flies having clearly visible ovarioles (normal), weakly visible ovarioles (weak), and no discernible ovarioles (dysgenic) at 29°C. Note that we did not obtain any viable females for some generations of R2. The percentage of dysgenic ovaries (HD) is computed as $100 * (dysgenic + (weak/2))/(normal + weak + dysgenic)$

| rep | gen | normal | weak | dysgenic | dysgenic [%] |
|---|---|---|---|---|---|
| R1 | 1 | 28 | 3 | 2 | 10.6 |
| R2 | 1 | 28 | 2 | 1 | 6.5 |
| R4 | 1 | 65 | 6 | 4 | 9.3 |
| R1 | 5 | 73 | 0 | 38 | 34.2 |
| R2 | 5 | 30 | 0 | 13 | 30.2 |
| R4 | 5 | 37 | 5 | 35 | 48.7 |
| R1 | 10 | 61 | 2 | 8 | 12.7 |
| R2 | 10 | 35 | 5 | 22 | 39.5 |
| R4 | 10 | 26 | 0 | 8 | 23.5 |
| R1 | 20 | 0 | 0 | 23 | 100.0 |
| R2 | 20 | 0 | 0 | 27 | 100.0 |
| R4 | 20 | 15 | 1 | 5 | 26.2 |
| R1 | 30 | 24 | 0 | 8 | 25.0 |
| R2 | 30 | - | - | - | - |
| R4 | 30 | 76 | 3 | 12 | 14.8 |
| R1 | 34 | 42 | 4 | 8 | 18.5 |
| R2 | 34 | 7 | 2 | 9 | 55.6 |
| R4 | 34 | 34 | 0 | 5 | 12.8 |
| R1 | 41 | 25 | 1 | 2 | 8.9 |
| R2 | 41 | - | - | - | - |
| R4 | 41 | 94 | 5 | 16 | 16.1 |
| R1 | 45 | 83 | 0 | 2 | 2.4 |
| R2 | 45 | - | - | - | - |
| R4 | 45 | 60 | 1 | 6 | 9.7 |
| R1 | 46 | 67 | 1 | 11 | 14.6 |
| R2 | 46 | - | - | - | - |
| R4 | 46 | 92 | 0 | 17 | 15.6 |
| R1 | 50 | 41 | 2 | 15 | 27.6 |
| R2 | 50 | - | - | - | - |
| R4 | 50 | 25 | 1 | 2 | 8.9 |
| - | - | 23 | 0 | 1 | 4.2 |
| - | - | 18 | 0 | 3 | 14.3 |
| - | - | 28 | 0 | 1 | 3.4 |

Table 7: Overview of ONT long-read data used in this work. For each run we show the mean, median and N50 of the read length. Data are shown for different replicates (rep.) and generations (gen.)

| rep. | gen. | run | flowcell | output [Gb] | reads [m.] | N50 | mean | median |
|---|---|---|---|---|---|---|---|---|
| R1 | G20 | 1 | R9.4.1 | 14.66 | 2.47 | 12,968 | 5,917 | 2,891 |
| R2 | G18 | 1 | R9.4.1 | 10.68 | 2.04 | 11,346 | 5,224 | 2,723 |
| R2 | G21 | 1 | R9.4.1 | 11.43 | 2.21 | 7,566 | 5,157 | 4,121 |
| R2 | G21 | 2 | R9.4.1 | 6.88 | 1.8 | 5,508 | 3,724 | 2,793 |
| R2 | G21 | 3 | R9.4.1 | 2.40 | 0.36 | 9,646 | 6,611 | 5,087 |
| R2 | G26 | 1 | R9.4.1 | 5.66 | 0.96 | 8,773 | 5,875 | 4,445 |
| R2 | G51 | 1 | R9.4.1 | 3.65 | 0.73 | 9,575 | 4,954 | 3,002 |
| R2 | G51 | 2 | R9.4.1 | 25.5 | 6.55 | 7,833 | 3,901 | 2,205 |
| R4 | G25 | 1 | R9.4.1 | 10.7 | 1.4 | 12,239 | 7,610 | 4,739 |
| R4 | G51 | 1 | R9.4.1 | 15.9 | 3.85 | 8,037 | 4,137 | 2,442 |

Table 8: Overview of P-element insertions in piRNA clusters that are supported by at least two ONT reads at early generations of the experimental populations ($g. \leq 26$). Reads from different ONT libraries (runs) of the same sample were merged. For each insertion we show the contig, the position (*pos*), the maximum number of bases aligning to the P-element (*sup*; the length of the P-element is 2,907bp) and the orientation of the P-element insertion (*ori*). Given the coverage (*cov*) and the number of reads ($r$) supporting a P-element insertion the population frequency of the insertion ($f = r/cov$) can be computed. The coverage was inferred from IGV and the number of reads supporting an insertion was corrected ($+x$ in column $r$) if a read was missed by our automated approach (for very long ONT reads with central P-element insertions, minimap2 may miss supplementary alignments to the P-element; these P-element insertions are instead reported as long insertions ). For 5kb windows around each insertion we also show the expression level of piRNAs (*expr* uniquely mapping piRNAs per million piRNAs), the strand bias (*s.b*: -1 all piRNAs are antisense, 1 all piRNAs are sense, 0 equal amounts of piRNAs are sense and antisense) and the degree of maternal transmission (*m.t*: 0 no piRNAs are found in the embryo, 1 equal amounts of piRNA in ovaries and embryo). Note that the low population frequency ($f$) of the insertions is not compatible with a model assuming that a fixed cluster insertions is responsible for silencing the P-element around generation 20 in replicates 1 and 4 (R1, R4). rep. replicate

| rep. | g. | contig | pos | cov | r | f | sup | ori | expr | s.b | m.t |
|---|---|---|---|---|---|---|---|---|---|---|---|
| R1 | 20 | contig_232 | 1,319,408 | 73 | 6+2 | 0.11 | 2,021 | fwd | 39.40 | 0.62 | 0.51 |
| R1 | 20 | contig_26 | 428,194 | 92 | 2 | 0.02 | 2,437 | rev | 809.94 | -0.88 | 0.06 |
| R1 | 20 | contig_422 | 1,690,050 | 203 | 3 | 0.01 | 2,907 | fwd | 1049.45 | 0.82 | 0.61 |
| R1 | 20 | contig_422 | 9,830,474 | 95 | 3 | 0.03 | 2,907 | fwd | 2291.91 | -0.21 | 0.06 |
| R2 | 18 | contig_508 | 579,832 | 62 | 7+2 | 0.14 | 2,389 | rev | 9.11 | -0.47 | 0.84 |
| R2 | 21 | contig_232 | 3,074,029 | 180 | 2 | 0.01 | 2,825 | fwd | 225.86 | -0.96 | 0.02 |
| R2 | 21 | contig_508 | 579,833 | 134 | 5+2 | 0.05 | 1,559 | rev | 9.11 | -0.47 | 0.84 |
| R4 | 25 | contig_513 | 12,838,058 | 91 | 3+1 | 0.04 | 1,434 | fwd | 1128.25 | 1.00 | 0.37 |
| R4 | 25 | contig_513 | 28,883,839 | 64 | 3 | 0.05 | 2,613 | rev | 24.30 | -0.47 | 0.24 |

Table 9: Overview of reads supporting P-element insertions inside (*obs.ci*) and outside of piRNA clusters (*non.ci*) in our ONT long-read libraries. Based on the shotgun model each diploid individual should carry about four P-element insertions in a piRNA cluster, i.e. two insertions per haploid genome [Kofler et al., 2018, Kofler, 2019]. Given that the average coverage (*cov*) approximates the number of haploid genomes present in a sample, the number of cluster insertions expected under the shotgun model can be computed ($exp.ci = 2 * cov$). Note that, in all samples, the number of P-element insertions in piRNA clusters is lower than expected under the shotgun model ($obs.ci/exp.ci < 1$).

| rep | gen | run | cov | non.ci | obs.ci | exp.ci | obs./exp |
|---|---|---|---|---|---|---|---|
| R1 | G20 | 1 | 90.5 | 1043 | 18 | 181.0 | 0.10 |
| R2 | G18 | 1 | 53.2 | 685 | 12 | 106.5 | 0.11 |
| R2 | G21 | 1 | 58.5 | 1394 | 4 | 116.9 | 0.03 |
| R2 | G21 | 2 | 31.8 | 854 | 3 | 63.6 | 0.05 |
| R2 | G21 | 3 | 12.1 | 450 | 7 | 24.2 | 0.29 |
| R2 | G26 | 1 | 28.8 | 1267 | 6 | 57.7 | 0.10 |
| R2 | G51 | 1 | 12.4 | 1239 | 17 | 24.8 | 0.69 |
| R2 | G51 | 2 | 127.2 | 14758 | 170 | 254.4 | 0.67 |
| R4 | G25 | 1 | 63.7 | 1619 | 8 | 127.5 | 0.06 |
| R4 | G51 | 1 | 70.9 | 2521 | 26 | 141.8 | 0.18 |

Table 10: Normalized number of reads aligning to *Drosophila* viruses in the small RNA libraries [rpm]. We sequenced small RNAs for each replicate (r.) at multiple generations (gen.). Small RNAs were sequenced either from ovaries (tis.: ov) or whole bodies of female flies (tis.: b). Additionally, we sequenced whole bodies and ovaries of naive flies not having the P-element (marked with '-' at r. and gen.). Data are shown for the five most abundant viruses across the libraries. 1: MG969167 Mauternbach Nudivirus, 2: MZ852356 picorn-like virus, 3: BlastFreeCandidate48, 4: Gosford Narnavirus, 5: MF893259 Teise virus segment 1). No significant differences in the abundance of any of these viruses was found between replicate 2 and replicates 1+4 (Wilcoxon rank sum tests; $p \geq 0.198$

| r. | gen. | tis. | 1 | 2 | 3 | 4 | 5 |
|---|---|---|---|---|---|---|---|
| R1 | 1 | ov | 53.32 | 171.63 | 1.01 | 15.15 | 7.98 |
| R1 | 5 | b | 127.44 | 26.34 | 47.65 | 17.72 | 76.28 |
| R1 | 10 | ov | 19.99 | 9.67 | 0.96 | 23.1 | 18.68 |
| R1 | 15 | b | 98.14 | 5.66 | 7.85 | 18.12 | 29.05 |
| R1 | 20 | b | 125.65 | 0.47 | 95.27 | 17.87 | 11.42 |
| R1 | 25 | b | 111.64 | 2.41 | 69.14 | 17.56 | 17.08 |
| R1 | 30 | b | 48.59 | 18.16 | 41.72 | 14.56 | 10.8 |
| R1 | 35 | ov | 100.27 | 13.27 | 0.52 | 42.62 | 5.16 |
| R1 | 40 | b | 24.91 | 0.21 | 55.41 | 25.74 | 9.72 |
| R1 | 45 | b | 29.93 | 2.45 | 57.12 | 14.75 | 12.23 |
| R2 | 1 | ov | 39.71 | 245.37 | 0.52 | 15.16 | 7.11 |
| R2 | 5 | b | 42.1 | 2.03 | 47.43 | 20.94 | 32.12 |
| R2 | 10 | ov | 639.87 | 11.87 | 0.42 | 16.87 | 60.95 |
| R2 | 15 | b | 75.24 | 6.41 | 5.64 | 16.4 | 27.67 |
| R2 | 20 | b | 136.03 | 3.81 | 54.22 | 17.19 | 14.34 |
| R2 | 25 | b | 347.03 | 2.96 | 64.39 | 19.17 | 16.48 |
| R2 | 30 | b | 123.75 | 3.99 | 26.39 | 18.81 | 25.24 |
| R2 | 35 | ov | 103.95 | 17.88 | 1.21 | 36.97 | 5.76 |
| R2 | 40 | b | 178.61 | 0.46 | 44.73 | 19.22 | 16.77 |
| R2 | 45 | b | 61.52 | 2.22 | 29.83 | 20.25 | 14.59 |
| R4 | 1 | ov | 68.07 | 575.14 | 0 | 14.54 | 6.18 |
| R4 | 5 | b | 80.64 | 0.55 | 28.03 | 20.77 | 33.16 |
| R4 | 10 | ov | 293.01 | 9.49 | 1.87 | 12.36 | 21.99 |
| R4 | 15 | b | 187.66 | 16.69 | 10.56 | 12.9 | 34.68 |
| R4 | 20 | b | 59.82 | 3.31 | 78.04 | 20.55 | 12.18 |
| R4 | 25 | b | 122.56 | 2.01 | 63.14 | 17.36 | 12.99 |
| R4 | 30 | b | 69.05 | 4.89 | 26.32 | 20.34 | 14.35 |
| R4 | 35 | ov | 62.97 | 40.51 | 1.05 | 42.86 | 2.55 |
| R4 | 40 | b | 40.57 | 0.64 | 89.58 | 25.75 | 10.99 |
| R4 | 45 | b | 97.99 | 4.3 | 44.64 | 19.24 | 13.18 |
| - | - | b | 41.89 | 2.4 | 92.69 | 22.27 | 10.11 |
| - | - | b | 50.12 | 1.22 | 24.52 | 25.45 | 10.51 |
| - | - | b | 155.05 | 10.3 | 47.3 | 22.66 | 25.04 |
| - | - | ov | 29.87 | 4.43 | 0.39 | 42.47 | 5.21 |
| - | - | ov | 48.22 | 13.69 | 0.58 | 32.37 | 3.4 |
| - | - | ov | 25.75 | 2.97 | 0.26 | 44.51 | 3.05 |

Table 11: P-element polymorphisms in the experimental populations. The coverage (cov) and allele frequency of the minor allele (%) are shown for polymorphisms having a minimum frequency of 5% and a minimum allele count of 2 in at least one generation (G) of the experiment. rep replicate, ma major allele, mi minor allele

| | | | | G1 | | G10 | | G20 | | G34 | | G40 | | G48 | |
|---|---|---|---|---|---|---|---|---|---|---|---|---|---|---|---|
| rep | pos | ma | mi | cov | % | cov | % | cov | % | cov | % | cov | % | cov | % |
| 1 | 571 | G | A | 26 | 7.69 | 123 | 0.00 | 869 | 0.23 | 840 | 0.12 | 1260 | 0.00 | 836 | 0.00 |
| 1 | 1342 | G | T | 24 | 8.33 | 106 | 0.00 | 786 | 0.00 | 877 | 0.00 | 1159 | 0.00 | 1005 | 0.10 |
| 1 | 1758 | T | A | 20 | 10.00 | 110 | 0.00 | 861 | 0.12 | 911 | 0.11 | 1251 | 0.00 | 1065 | 0.19 |
| 2 | 100 | T | G | 34 | 5.88 | 83 | 0.00 | 743 | 0.67 | 4919 | 0.69 | 4910 | 0.49 | 7083 | 0.37 |
| 2 | 1924 | T | A | 32 | 6.25 | 107 | 0.00 | 708 | 0.00 | 3274 | 0.21 | 3384 | 0.06 | 3586 | 0.03 |
| 2 | 2399 | A | G | 30 | 10.00 | 83 | 0.00 | 661 | 0.00 | 4706 | 0.08 | 5148 | 0.04 | 6832 | 0.00 |
| 2 | 2800 | A | T | 24 | 8.33 | 95 | 0.00 | 671 | 0.30 | 4692 | 0.06 | 5216 | 0.06 | 6445 | 0.02 |
| 4 | 570 | C | T | 21 | 9.52 | 327 | 0.00 | 1276 | 0.16 | 1213 | 0.00 | 1244 | 0.00 | 994 | 0.00 |
| 4 | 1193 | A | T | 36 | 5.55 | 285 | 0.00 | 1222 | 0.00 | 1222 | 0.00 | 1340 | 0.00 | 961 | 0.10 |
| 4 | 1304 | C | T | 15 | 13.33 | 291 | 0.00 | 1157 | 0.09 | 1145 | 0.00 | 1145 | 0.00 | 860 | 0.00 |
| 4 | 1904 | A | T | 19 | 10.53 | 297 | 0.67 | 1142 | 0.18 | 1166 | 0.34 | 1157 | 0.00 | 1019 | 0.00 |
